# Characterization of TelE, a T7SS LXG effector exhibiting a conserved C-terminal glycine zipper motif required for toxicity

**DOI:** 10.1101/2022.09.07.506920

**Authors:** Wooi Keong Teh, Yichen Ding, Francesca Gubellini, Alain Filloux, Claire Poyart, Michael Givskov, Shaynoor Dramsi

**Affiliations:** Singapore Centre for Environmental Life Sciences Engineering, Nanyang Technological University, Singapore; Present address: National Public Health Laboratory, Singapore; Institut Pasteur, Unité de Microbiologie Structurale, Paris, France; Centre for Bacterial Resistance Biology, Department of Life Sciences, Imperial College London, London SW7 2AZ, UK; Université de Paris, Assistance Publique Hôpitaux de Paris, Service de Bactériologie, Centre National de Référence des Streptocoques, Groupe Hospitalier Paris Centre site Cochin, Paris, France; Institut Pasteur, Université Paris Cité, CNRS UMR6047, Biology of Gram-positive Pathogens Unit, Paris 75015 France.; Centre National de la Recherche Scientifique (CNRS) UMR2001, Paris, France; Costerton Biofilm Centre, Department of Immunology and Microbiology, University of Copenhagen, Denmark

**Keywords:** *Streptococcus gallolyticus*, Type VIIb secretion system, pore-forming toxin, LXG family toxin, glycine zipper

## Abstract

*Streptococcus gallolyticus* subsp. *gallolyticus (SGG)* is an opportunistic bacterial pathogen strongly associated with colorectal cancer. Here, through comparative genomics analysis, we demonstrated that the genetic locus encoding the Type VIIb Secretion System (T7SSb) machinery is uniquely present in *SGG* in two different arrangements. *SGG* UCN34 carrying the most prevalent T7SSb genetic arrangement was chosen as the reference strain. To identify the effectors secreted by this secretion system, we inactivated the essC gene encoding the motor of this machinery. Comparison of the proteins secreted by UCN34 WT and its isogenic ΔessC mutant revealed six T7SSb effector proteins, including the expected WXG effector EsxA and three LXG-containing proteins. In this work, we characterized an LXG-family toxin named herein TelE displaying pore-forming activity. Seven homologs of TelE harboring a conserved glycine zipper motif at the C-terminus were identified in different *SGG* isolates. Scanning mutagenesis of this motif showed that the glycine residue at position 470 was crucial for TelE pore-forming activity. Unlike other pore-forming toxins commonly antagonized by a membrane protein, TelE activity was antagonized by a small protein TipE belonging to the DUF5085 family. Overall, we report herein a unique *SGG* T7SSb effector exhibiting a pore-forming activity against non-immune bacteria.

**IMPORTANCE:** In this study, 38 clinical isolates of *Streptococcus gallolyticus* subsp*. gallolyticus* (*SGG*) were sequenced and a genetic locus encoding the Type VIIb secretion system (T7SSb) was found conserved and absent from 16 genomes of the closely related *S. gallolyticus* subsp. *pasteurianus (SGP)*. The T7SSb is a *bona fide* pathogenicity island. Here, we report that the model organism *SGG* strain UCN34 secretes six T7SSb effectors. One of the six effectors named TelE displayed a strong toxicity when overexpressed in *Escherichia coli*. Our results indicate that TelE is a pore forming toxin whose activity can be antagonized by a non-canonical immunity protein named TipE. Overall, we report a unique toxin-immunity protein pair and our data expand the range of effectors secreted through T7SSb.

## INTRODUCTION

*Streptococcus gallolyticus* subsp*. gallolyticus* (*SGG*) is a commensal bacterium in the rumen of herbivores. The first complete genome of strain UCN34 provided clear insights for its adaptation to the rumen (1). *SGG* is also an opportunistic pathogen causing septicemia and endocarditis in the elderly who often have concomitant colon tumours (2-9). *SGG* was formerly known as *S. bovis* biotype I belonging to the large *S. bovis* / *equinus* complex, which is comprised of several opportunistic pathogens and non-pathogenic bacteria. In this complex, *SGG* is assigned into a clade with two other closely related subspecies named *S. gallolyticus* subsp. *pasteurianus* (referred to hereafter as *SGP*) and *S. gallolyticus* subsp. *macedonicus* (referred to hereafter as *SGM*) (10, 11). In general, *SGP* is considered less pathogenic than *SGG*, mainly causing neonatal meningitis (12, 13), whereas *SGM* is generally recognized as a safe non-pathogenic strain (14, 15).

The Type VIIb Secretion System (T7SSb) is a specialized secretion system exploited by Firmicutes to secrete various substrates or effectors involved in host immune system modulation or bacterial competition (16-20). T7SSb is distantly related to the Type VII secretion system originally discovered in *Mycobacterium tuberculosis.* These two evolutionarily related secretion systems have two components in common; a membrane-bound ATPase (FtsK/SpoEIII, EssC/EccC) constituting the motor of the secretion machinery, and a secreted effector named EsxA/ESAT6 belonging to the WXG100 family (21). In Firmicutes, the T7SSb core machinery is composed of 4 additional proteins namely EssA, EsaA, EsaB and EssB. This specialized secretion system is involved in the secretion of a variety of effectors including (i) WXG100 proteins such as EsxA and EsxB which are short alpha-helical proteins with a characteristic WXG motif in the middle of the sequence (22); (ii) proteins of approximately 100 amino acids without a WXG motif (named WXG100-like proteins) such as EsxC and EsxD (23, 24); (iii) polymorphic toxins containing a N-terminal LXG domain and a variable C-terminal domain with antibacterial activity such as TelA, TelB, TelC, TelD, TspA and EsxX (18, 25-28). The LXG domain is predicted to adopt an alpha-helical structure that is reminiscent of the WXG100 proteins. Previous studies revealed that these LXG-containing effectors function as a NADase toxin (TelB) (18), a lipid II phosphatase (TelC) (18), nucleases (EsaD, YobL,, YxiD) (16, 27, 29), or a membrane depolarizing toxin (TspA) (25).

Importantly, the T7SSb of *SGG* strain TX20005 was shown to contribute to the development of colon tumors in a murine model (30). Here, using comparative genomics, we identified that the T7SSb locus residing in a 26-kb genomic island is uniquely present in *SGG* but neither in *SGP* nor in *SGM*. Subsequently, we found that six proteins were secreted through the T7SSb machinery of the reference strain UCN34, including the WXG100 effector EsxA, two hypothetical proteins, and three LXG family proteins including two TelC homologs. One of the unknown LXG family protein was characterized herein and named TelE. We show that TelE is a pore-forming protein whose activity is antagonized by a non-canonical immunity protein.

## RESULTS

### A T7SSb locus specific to *SGG*

To identify the specific pathogenic trait(s) of *SGG*, comparative genomics was performed on newly sequenced 13 *SGP* and 38 *SGG* clinical isolates obtained from the National Reference Center of Streptococci in France. Phylogenetic analysis of these newly sequenced genomes, together with the complete genomes of *SGG* and *SGP* publicly available on NCBI GenBank assigned *SGG* and *SGP* into two distinct clades, confirming their correct classification **(Fig. S1a)**. BLASTn-based comparative genomic analysis of the reference genome UCN34 together with the representative *SGG* and *SGP* genomes uncovered a genomic island unique to *SGG* **(Fig. 1A, Fig. S1b**). This genomic island contains the six genes (*esxA*, *essA*, *esaB*, *essB*, *essC* and *esaA*) encoding the core components of the Type VIIb Secretion System (T7SSb) machinery **(Fig. 1B, Fig. S1b)**.

**Figure 1.**
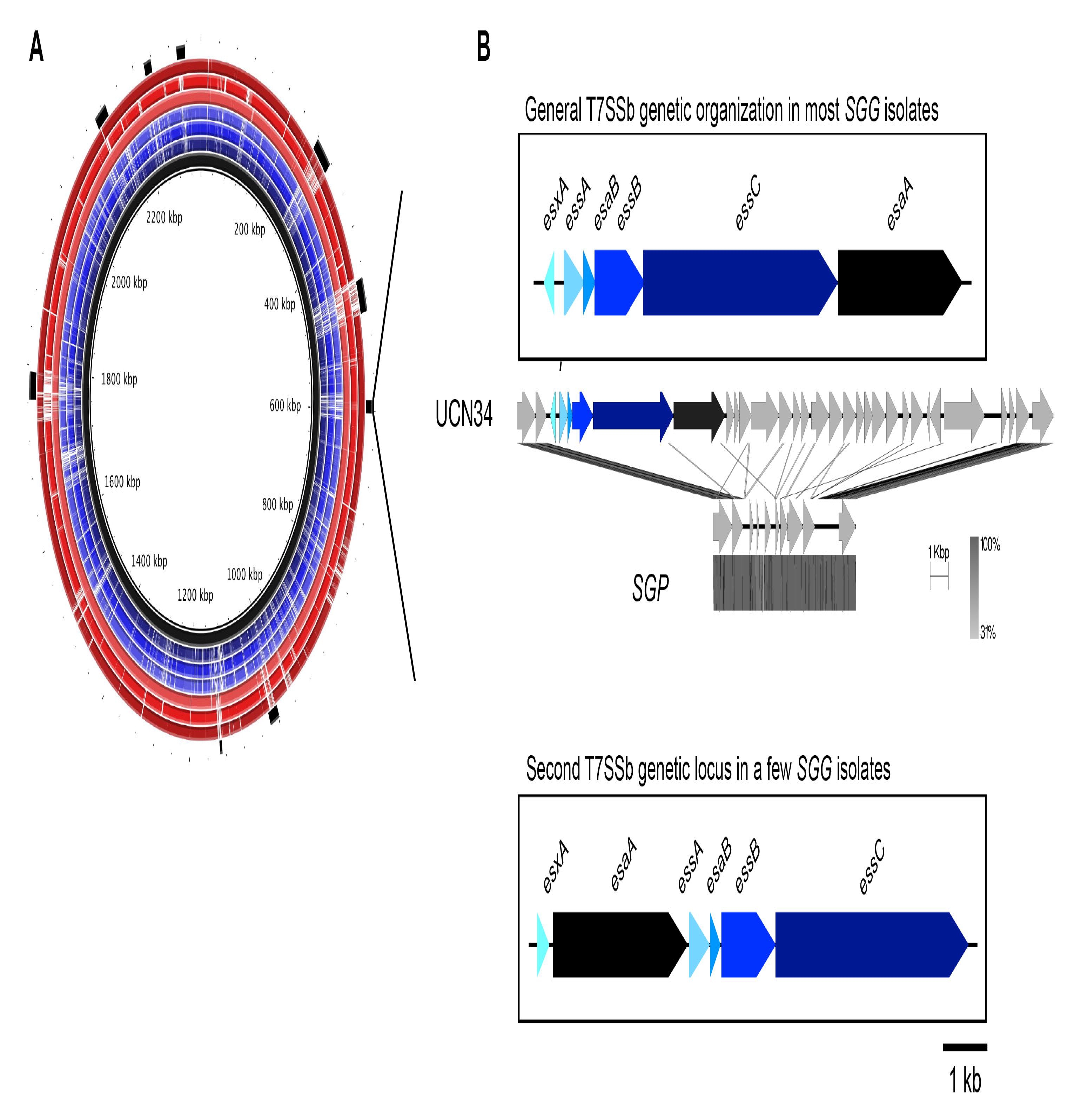
A T7SSb locus specific to *SGG*. (A) An overview of the BLASTn-based comparative genomics analysis. From the innermost circle, the reference genome *SGG* UCN34 (black ring); followed by *SGP* isolates (Gallo1, Gallo4, Gallo6, blue rings); and *SGG* isolates (Gallo37, Gallo43 and Gallo51, red rings). The outermost black blocks represent the genomic islands in UCN34 genome predicted by IslandViewer 4. White spaces within each ring represent a ≤50% similarities of the respective genomic regions to UCN34. (B) Genetic organization of the two types of T7SSb found in *SGG* clinical isolates and absent in *SGP*. Each T7SSb gene encoding the secretion machinery was color-coded and labelled accordingly.

An extended protein homology search of each T7SSb core components across all the newly sequenced genomes confirmed the conservation of T7SSb in *SGG* genomes and its absence in the phylogenetically closest relatives *SGP* and *SGM* **(Fig. S1b)**. A closer inspection of the T7SSb genetic locus uncovered two different gene arrangements in the *SGG* isolates sequenced so far. In a vast majority of *SGG* isolates, the T7SSb locus is arranged in the order of *esxA/essA-esaB-essB-essC- esaA*, as in our reference strain UCN34. Notably, *esxA* is transcribed in the opposite direction from the other five genes *essA-esaB-essB-essC-esaA,* a unique feature distinct from other Firmicutes. In contrast, in a small minority of *SGG* isolates including the US isolate TX20005, the T7SSb genes are arranged in a different order of *esxA-esaA-essA-esaB-essB-essC* with an evident shuffling of *esaA*, and the six genes transcribed in the same direction **(Fig. 1B, Fig. S1b**). A detailed comparison of the genetic organization of the entire T7SSb locus in the representative *SGG* strains UCN34 and TX20005 is shown in **Fig. S2**. In addition to a different gene organization of the core machinery, the set of putative T7SSb effectors is different and appears to be strain specific.

### *SGG* UCN34 secretes 6 T7SSb effectors

In order to identify the T7SSb effectors, we inactivated the T7SSb machinery, through in-frame deletion of *essC* gene encoding the main ATPase, in the strain UCN34. We next compared the secretome of the UCN34 WT strain to that of the Δ*essC* mutant. This approach identified 6 proteins that were differentially expressed **(Fig. 2A**, **Table S2**) including the ubiquitous WXG100 effector (EsxA/Gallo_0553) whose level was about 38-fold less abundant in the Δ*essC* mutant, confirming that the Δ*essC* mutant is a *bona fide* T7SSb-deficient strain. In addition, five proteins namely Gallo_0559, Gallo_0560, Gallo_0562, Gallo_1068 and Gallo_1574 were detected only in the secretome of the UCN34 WT but not in the Δ*essC* mutant.

**Figure 2.**
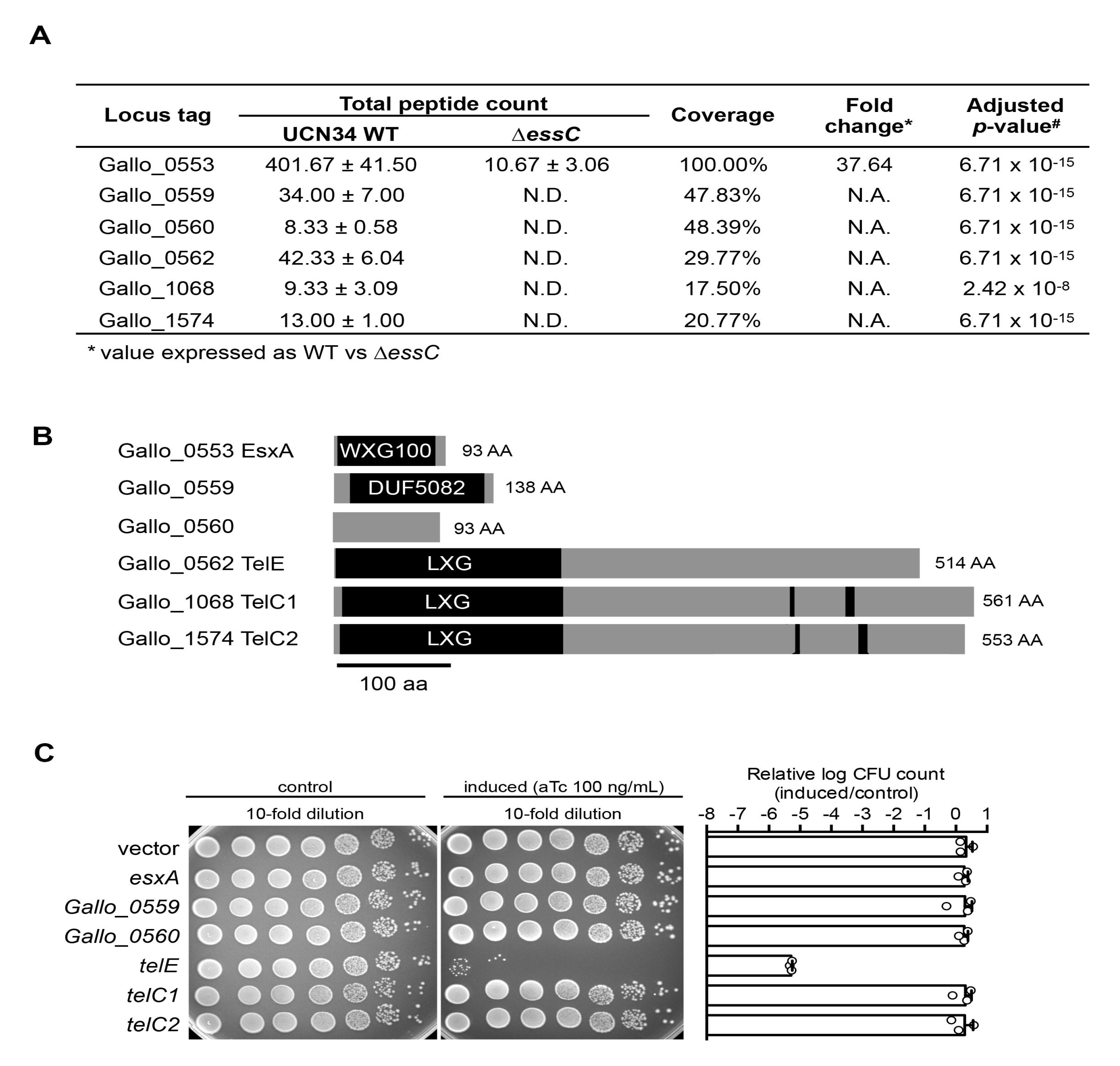
A *SGG* T7SSb effector exhibits toxicity in *E. coli*. (A) List of proteins that were significantly more abundant in the secretome of the UCN34 wildtype than the Δ*essC* mutant. The total peptide count of each protein was expressed as means ± standard deviation from three biological replicates. N.D., not detected; N. A., not applicable. (B) The domain architecture of the six proteins identified in (A). Both Gallo_1068 and Gallo_1574 are homologous to the lipid II phosphatase TelC. The conserved aspartate-rich motifs essential for TelC activity are indicated by black boxes. (C) Viability of the bacterial cells harboring either an empty vector (pTCVerm-P_tetO_) or a vector encoding the full-length *telE* with the expression inducible by anhydrotetracycline (aTc). Bar chart on the right showed the logarithm-transformed CFU count of the *telE*-expressing cells in relative to the control. Error bars represent mean + standard deviation (n = 3).

Prediction of the putative function of these effectors by *in silico* SMART analysis revealed that Gallo_0559 contains a domain of unknown function DUF5082, Gallo_0560 as a TIGR04197 family protein, whereas Gallo_0562, Gallo_1068 and Gallo_1574 contain the prototypical N-terminal LXG domain (pfam04740) found in other T7SSb effectors **(Fig. 2B)**. Protein homology search indicated that Gallo_1068 and Gallo_1574 are homologous (>98% coverage, >40% similarity) to the *Streptococcus intermedius* TelC effector that functions as a lipid II phosphatase (18). Similar to TelC, both Gallo_1068 and Gallo_1574 contain an intact C-terminal aspartate-rich motif critical for TelC enzymatic activity. We thus propose to rename Gallo_1068 and Gallo_1574 as TelC1 and TelC2, respectively **(Fig. 2B)**. The third LXG-containing effector (Gallo_0562) is a protein of 514 amino acids that we propose to rename TelE in accordance with the gene nomenclature (<u>T</u>oxin exported by <u>E</u>sx with <u>L</u>XG domain) proposed for *S. intermedius* (18). Of note, 4 out of 6 proteins identified in the secretome of UCN34 WT are encoded in the T7SSb locus and three are encoded by genes likely transcribed as an operon, namely *gallo_0559*, *gallo_0560* and *telE (gallo_0562)*.

### TelE is toxic when overexpressed in *Escherichia coli*

LXG family proteins are polymorphic toxins implicated in interbacterial competition (18). However, we failed to observe any role of *SGG* T7SSb in interbacterial competition under standard laboratory conditions. We reasoned that this could be due to the low expression of T7SSb effectors as reflected by the low total peptide counts in the proteomics data **(Fig. 2A)** and decided to test their potential antibacterial activity through heterologous and inducible expression in *Escherichia coli* DH5α, as reported for other T7SSb effectors such as TspA and TelD (25, 28). As shown in **Fig. 2C**, overexpression of *telE*, but none of the other T7SSb effectors, exhibited antibacterial activity in *E. coli* **(Fig. 2C)**.

TelE is encoded in the vicinity of the T7SSb core machinery locus **(Fig. 3A)**. This prompted us to examine the prevalence and the conservation of TelE among the *SGG* clinical isolates. As shown in **Fig. S3**, TelE homologs were found in over 90% of *SGG* isolates. Based on their primary amino acid sequences, TelE homologs can be categorized into 7 variants named TelE1 (initial TelE found in UCN34) through TelE7. Notably, multiple copies of TelE homologs can be found in a few *SGG* genomes. Some of these TelE homologs are N-terminally truncated, resembling the orphan modules commonly seen in polymorphic toxins (31) (**Fig. S3**). To assess the antimicrobial activity of these TelE homologs, each TelE variant was expressed in *E. coli.* As shown in **Fig. 3B**, all the TelE variants exhibited a strong toxicity upon induction in *E. coli*. Interestingly, a multiple sequence alignment of these seven TelE variants uncovered a highly conserved sequence motif at the C-terminus (**Fig. S4**). This motif, rich in small hydrophobic amino acids alternating with conserved glycine residues, is similar to the glycine zipper motif (GxxxGxxxG) found in many membrane proteins (32) **(Fig. 3C)**. Three highly conserved glycine residue were identified at positions 458, 470 and 474, and three less-conserved glycine residues were found at positions 466, 478 and 480 **(Fig. 3C and 3D)**. Single replacement of each glycine residue to a slightly bulkier valine residue demonstrated that those at positions 466, 470 and 474 were key for TelE activity in *E. coli* **(Fig. 3E)**, with the replacement of the glycine residue at position 470 showing the strongest impact on TelE activity **(Fig. 3E)**. Together, these data showed that nearly all the *SGG* isolates sequenced as to date encode a functional TelE homolog whose primary sequence varies from one isolate to another, but all 7 TelE variants contain a conserved glycine zipper motif at the C-terminus essential for TelE antimicrobial activity.

**Figure 3.**
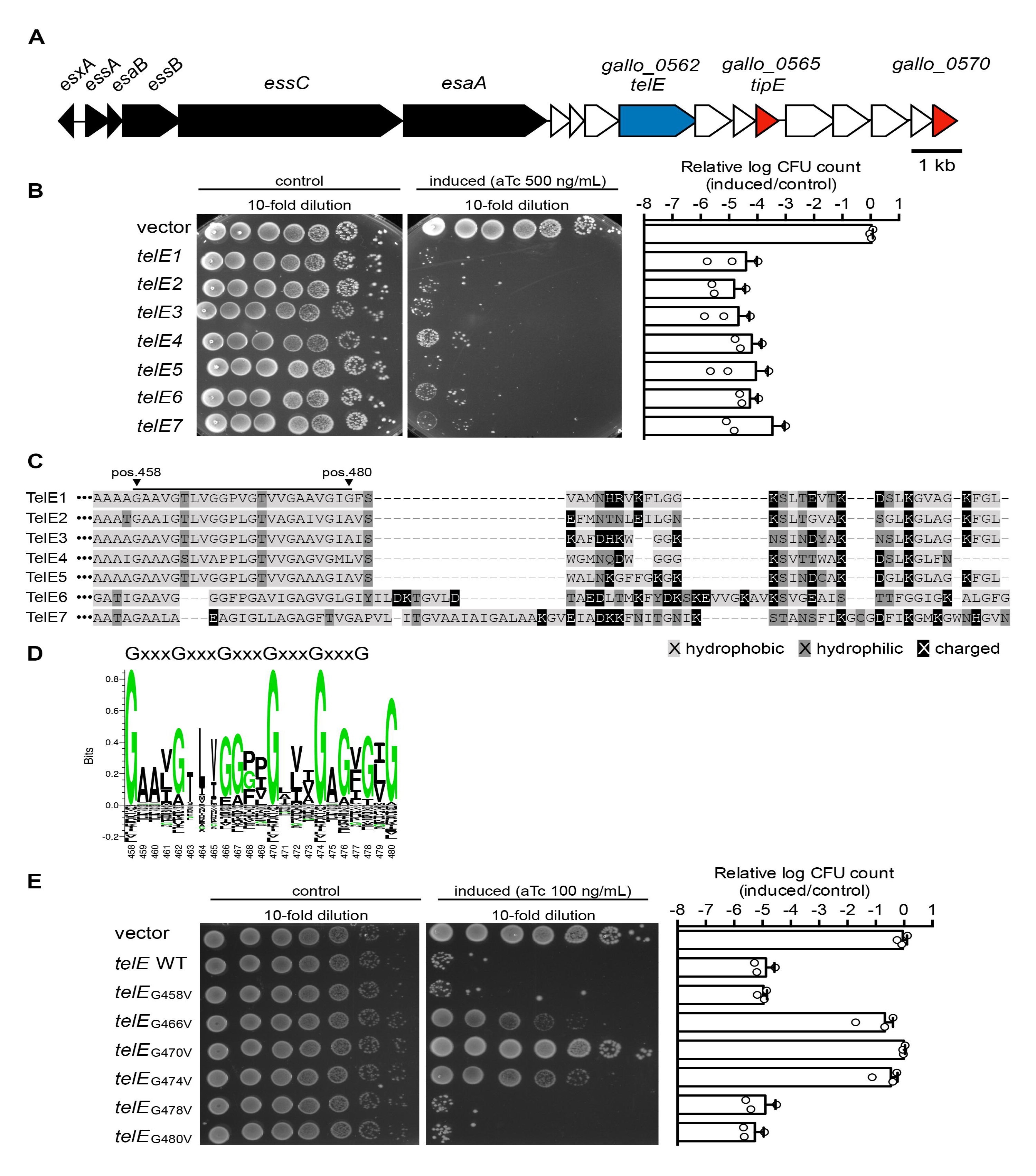
T7SSb Effector TelE contains a C-terminal glycine zipper motif essential for its activity. (A) The genomic region encoding the T7SSb core components, TelE and TipE in *SGG* UCN34. Black, T7SSb core components; blue, TelE (LXG domain-containing protein); red, DUF5085 domain-containing protein); white, hypothetical proteins. (B) Viability of the *E. coli* cells harboring either an empty vector ((pTCVerm-P_tetO_) or a vector encoding various *telE* variants (*telE1* to *telE7*) with the expression inducible by anhydrotetracycline (aTc). Bar chart on the right showed the logarithm-transformed CFU count of the *telE*-expressing cells in relative to the control. Error bars represent mean + standard deviation (n = 3). (C) Alignment of the C-terminus sequence of the *telE* variants expressed in (B) revealed a conserved region (underlined) enriched with hydrophobic residues. (D) Sequence logo of the underlined region in (C) generated from all aligned TelE variants identified a sequence motif highly similar to glycine zipper motif (GxxxGxxxG). The number at the bottom of the sequence logo represents the position of the respective amino acid residue in reference to the TelE sequence from UCN34. Shrunk residues (*i.e.* residues at position 462 to 465) reflected the presence of gaps in the sequence alignment. (E) Viability of the *E. coli* cells harboring either an empty vector (pTCVerm-P_tetO_) or a vector encoding the mutated *telE* with the expression inducible by anhydrotetracycline (aTc). Bar chart on the right showed the logarithm-transformed CFU count of the *telE*-expressing cells in relative to the control. Error bars represent mean + standard deviation (n = 3).

### TelE is a pore-forming toxin

Structural modelling of TelE indicates a high number of alpha helices and the C-terminal domain of TelE exhibit a small stretch of amino acids weakly homologous to bacteriocin IIc and glycine zipper motif **(Fig. 4A)**. Alpha-helical structures with glycine zipper motifs are the two main characteristics of membrane proteins with a high propensity to form homo-oligomeric ion channels (32), suggesting that TelE might have a pore-forming activity. To investigate this possibility, we used propidium iodide, a large molecule (668 Da) that is conventionally used to assess the presence of membrane pores or membrane disruption in bacterial cells (33, 34). Propidium iodide can only enter the cellular cytoplasmic space through membrane lesions or large membrane pores to subsequently bind to intracellular DNA and emit a red fluorescence signal. Using time-lapse fluorescence microscopy, we observed a gradual increase of red fluorescence signal in *E. coli* cells expressing TelE, but not in *E. coli* cells carrying the empty vector **(Fig. 4B)**, indicating that TelE expression resulted in an influx of propidium iodide into the cells. Intriguingly, this influx was preceded immediately by shrinkage of the bacterial cell visible by phase contrast microscopy **(Fig. 4C)**. Similar morphological changes have been observed on cells experiencing water loss or cytoplasmic content leakage (35, 36), which can be due to the excessive presence of membrane pores. Our microscopy observations thus indicate that TelE can form membrane pores in *E. coli* cells. To monitor TelE expression, a chimeric TelE variant C-terminally fused with superfolder green fluorescent protein (TelE-sfGFP) was constructed and GFP fluorescence signals were measured to quantify TelE expression **(Fig. 4D)**. Of note, TelE-sfGFP retains TelE toxicity in *E. coli* cells (**Fig. S5A**). The non-toxic version of TelE (TelE_G470V_-sfGFP) was constructed in parallel. As shown in **Fig. 4D**, an increase of GFP fluorescence was seen upon induction of TelE expression over time from 10 to 60 min. TelE_G470V_-sfGFP fluorescence level was always higher compared to TelE-sfGFP. Induction of TelE-sfGFP in *E. coli* for 60 min led to a 2-log decrease in C.F.U counts as compared to the non-induced strain or the strain expressing the mutant version of *telE,* namely TelEG470V-gfp **(Fig. 4D)**. This result demonstrates that TelEG470V punctual mutant is produced as well as TelE and that its non-toxicity is not due to protein degradation.

**Figure 4.**
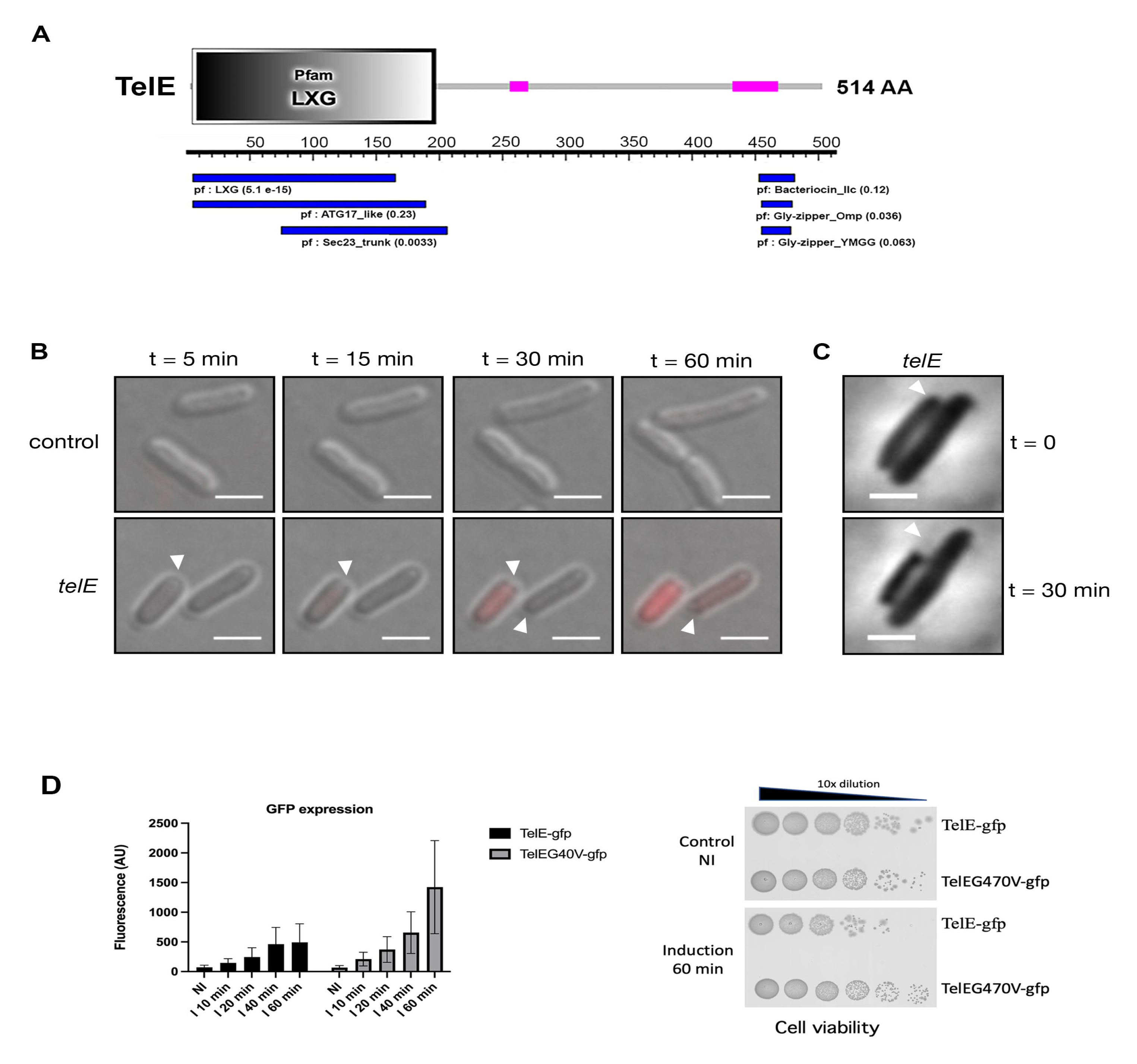
TelE is a pore-forming toxin. (A) The Pfam domains found in TelE protein using SMART (upper part) and motif search from KEGG Gene databases (lower part). The number in parenthesis next to the Pfam domain indicates the E-value score. (B) Differential interference contrast (DIC) micrograph of *E. coli* cells harboring an empty vector or a vector encoding *telE* with the expression inducible by anhydrotetracycline (aTc). The physiological changes of the *E. coli* cells following protein induction were documented at the indicated time. Cells with membrane lesions were stained red by propidium iodide. Arrows indicate the region with visible cell shrinkage. Scale bar, 2 µm. (C) Phase contrast micrograph of *telE*-expressing *E. coli* documented at the indicated time. Arrows indicate the region with visible cell shrinkage. Scale bar, 2 µm. (D) Monitoring of TelE expression through translational reporter constructs TelE-sfGFP or TelE_G470V_-sfGFP expressed in *E. coli* under the control of PtetO promoter. GFP fluorescence signal was measured at different time points following anhydrotetracyclin addition (I 10min-I 60 min) and compared to the initial non-induced (NI) culture. Cell viability was also monitored in parallel and shown for the 60 min induction time point.

### The cognate immunity protein TipE antagonizes TelE toxicity

The activity of LXG family toxins such as TelA, TelB and TelC can be antagonized by cognate immunity proteins located downstream of LXG-encoding gene (18). To identify the immunity protein able to counteract TelE toxicity in *E. coli*, each gene located downstream from *telE* was cloned into a second compatible plasmid (pJN105) in which gene expression was induced by arabinose. We found that only the co-expression of the third gene downstream from *telE* (*gallo_0565*) rescued *E. coli* from TelE intoxication **(Fig. 5A, Fig. S5A)**. This gene encodes a 151 amino acids-long protein renamed herein as TipE (<u>T</u>el immunity <u>p</u>rotein), which harbor a domain of unknown function DUF5085. An additional DUF5085 domain-containing protein (Gallo_0570), sharing ∼30% protein sequence similarity with TipE, was found five genes downstream of TipE **(Fig 3A)**. However, co-expression of this protein did not protect *E. coli* from TelE intoxication, suggesting a specific interaction between TipE and TelE. An AlphaFold prediction model of the TelE-TipE complex is shown in **Fig. 5B** (TelE in green and TipE in orange). The same model colored with the confidence value is shown in **Fig. S6**. The predicted interaction of TipE with the region including the Glycine-zipper motif of TelE (in yellow) fits well with the inhibitory effect of TipE on TelE.

**Figure 5.**
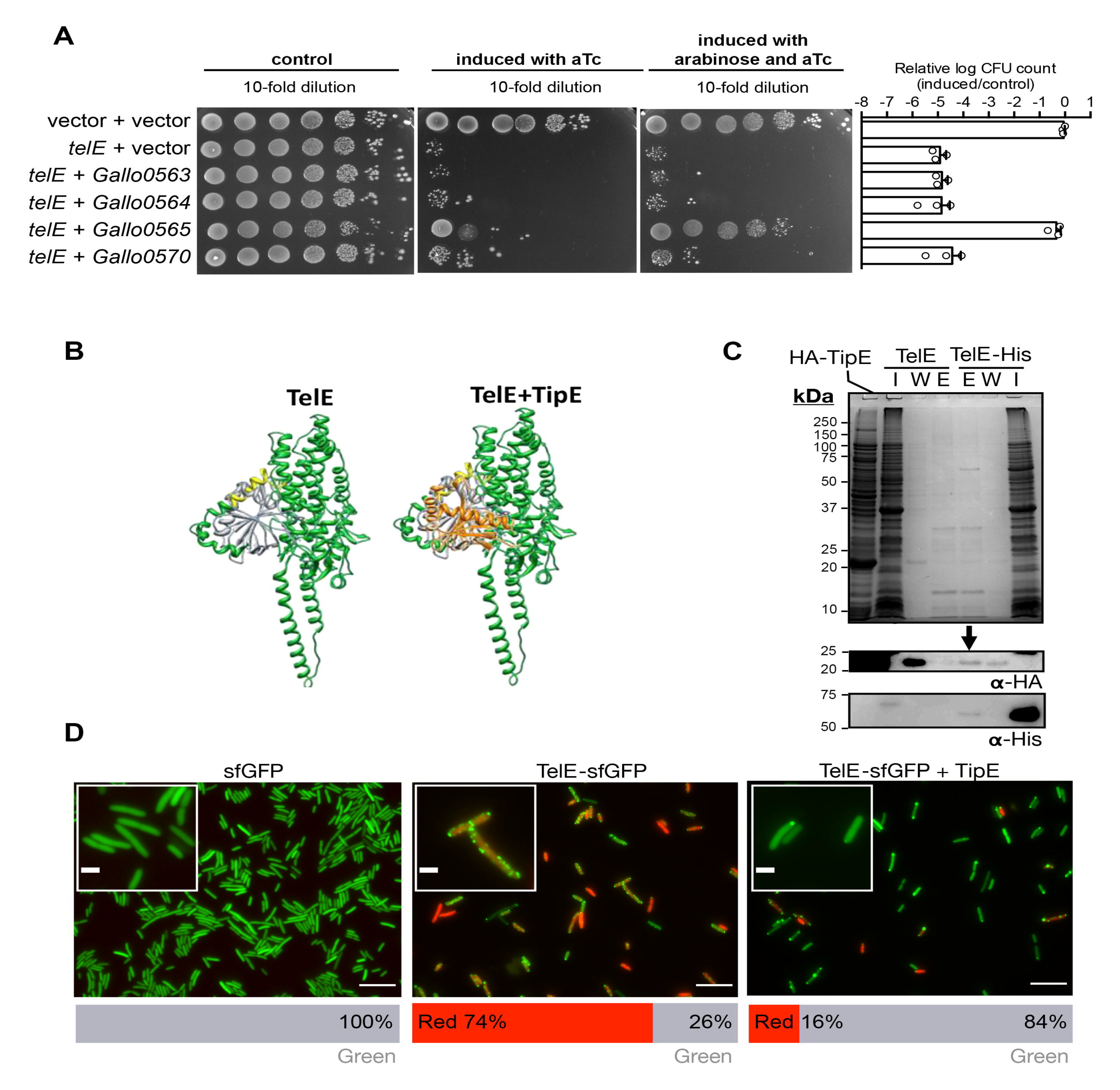
The cognate immunity protein TipE antagonizes TelE toxicity. (A) Viability of *E. coli* cells harboring two vectors (pTCVerm-P_tetO_ and pJN105-P_BAD_) for single expression of *telE* inducible by anhydrotetracycline (aTc), or co-expression of *telE* and the indicated gene candidates with the expression inducible by arabinose. Bar chart on the right showed the logarithm transformed CFU count of the cells co-expressing two genes in relative to the control. Error bars represent mean + standard deviation (n = 3). (C) AlphaFold modelling of monomeric TelE (in green) without or with TipE (in orange). TipE is predicted interacting with the TelE zipper motif shown in yellow. (C) Immunoblot with anti-His or anti-HA on the proteins eluted from the TelE-bound nickel-immobilized resins incubated with cellular lysate containing HA-TipE. The TelE-free resins subjected to the same treatment werem used as the control. I, input (cellular lysate containing HA-TipE); W, wash fraction; E, eluted fraction. (D) Fluorescence micrograph of *E. coli* cells expressing superfolder GFP (sfGFP), TelE-sfGFP, or co-expressing TipE and TelE-sfGFP. Cells with membrane lesions were stained red by propidium iodide. Scale bar 10 µm; 2 µm for inlets. Bar chart (bottom) showed the quantification of cells fluoresced green or red (n ≈ 300).

To demonstrate that TipE interacts with TelE, we constructed a functional, recombinant TelE tagged with a hexahistidyl tagged at the C-terminus (TelE-His) and a recombinant TipE tagged with the hemagglutinin tag (HA-TipE) at the N-terminus to facilitate the detection of these proteins using an anti-His and anti-HA monoclonal antibody, respectively. Of note, addition of a hexahistidyl tagged at the C-terminus of TelE reduces its toxicity towards *E. coli* as compared to the same tag added at the N-terminus (**Fig. S5C**). Co-expression of HA-TipE clearly increased TelE-His level as shown in **Fig. S5B**. Next, a protein pulldown experiment was carried out using nickel-immobilized resin that is able to retain TelE-His. Incubation of TelE-bound resin with cellular lysate containing HA-TipE captured a significant amount of TipE, which was detected through immunoblot using anti-HA specific antibodies following TelE elution **(Fig. 5C)**. Importantly, when using non-tagged TelE as control, HA-TipE was not retained on the free nickel affinity column **(Fig. 5C)**, indicating that TipE indeed interacts with TelE.

Subsequently, we monitored TelE expression in living cells by observing the *E. coli* cells expressing TelE-sfGFP under a fluorescence microscopy. Expectedly, TelE expression in *E. coli* cells resulted in cellular influx of propidium iodide **(Fig. 5D)**. However, instead of uniformly distributed throughout the cells as the native sfGFP **(Fig. 5D)**, TelE-sfGFP was found in small foci distributed around the cell body **(Fig. 5D)**. In contrast, in *E. coli* cells co-expressing TelE-sfGFP and TipE, TelE lost its distribution as foci in a majority of cells **(Fig. 5D)**. Notably, a small cell population co-expressing TelE-sfGFP and TipE remained stained red by propidium iodide, consistent with our earlier observation that TipE only partially rescued cells from TelE intoxication **(Fig. 5B)**. Taken together, these data showed that TipE is a *bona fide* immunity protein to TelE, counteracting TelE activity through protein-protein interaction.

## DISCUSSION

In this study, we show that a T7SSb cluster is a putative pathogenicity island conserved in all the clinical isolates of *Streptococcus gallolyticus subsp. gallolyticus* (*SGG*). Interestingly, two different T7SSb genetic arrangements can be found in *SGG* clinical isolates and a very different set of putative effectors as shown by comparing closely UCN34 and TX20005 (**Fig. S2**). The most prevalent T7SS machinery arrangement found in UCN34 resembles those found in *Streptococcus intermedius*, whereas the T7SS of a small minority of *SGG* isolates like TX20005 is identical to those of *Staphylococcus aureus* (16, 18) **(Fig. S1b)**. Of note, a second copy of the gene *essC* encoding the motor ATPase is present in TX20005 (**Fig. S2**). Six T7SSb effectors were identified experimentally in strain UCN34, including the prototypical EsxA/ESAT-6, and three LXG-containing proteins. Amongst the three LXG effectors, two encoded outside of the T7SSb locus are homologs of TelC, a lipid II phosphatase shown to inhibit peptidoglycan biosynthesis in Gram-positive bacteria (18, 37). The third LXG effector of unknown function is unique to *SGG* and is encoded in vicinity of the locus encoding T7SSb machinery. Here, we showed that this LXG-effector named TelE is toxic when overexpressed in *E. coli*. Similar to other LXG effectors, TelE toxicity in *E. coli* can be neutralized by a cognate immunity protein named TipE.

TipE is an atypical immunity protein. In other pore-forming toxins similarly containing glycine zipper motifs such as CdzC/D from *Caulobacter crescentus* and Tse4 from *Pseudomonas aeruginosa*, the cognate immunity proteins are multi-pass transmembrane proteins residing predominantly in the inner membrane (33, 38).

Similarly, an inner membrane protein protects the *E. coli* cells expressing the *S. aureus* membrane depolarizing T7SSb toxin TspA (25). Remarkably, the six-pass transmembrane protein Gallo_0563 encoded by the gene located immediately downstream of *telE* did not protect *E. coli* from TelE intoxication **(Fig. 5A, 5B)**, and our data showed that only TipE that contains no transmembrane domain is able to block TelE activity through direct interaction. Notably, TipE is a DUF5085 family protein predicted related to T7SS in the Pfam database. Based on our data, we propose herein that DUF5085 family proteins could be a new class of immunity protein to T7SSb toxins.

Interestingly, a diverse pool of TelE variants (TelE to TelE7) exist in the genomes of *SGG* isolates, all harboring the conserved C-terminal glycine zipper motif essential for TelE activity. Glycine zipper motif is a common feature in membrane proteins displaying pore-forming activity. In addition to the aforementioned CdzC/D and Tse4, glycine zipper motif is also found in the vacuolating toxin VacA from *Helicobacter pylori*, the prion protein (PrP) and the amyloid beta peptide (A*β*) (32), many of which display pathological roles. For instance, VacA is involved in gastric ulceration caused by *H. pylori*, whereas PrP and A*β* are associated with prion and Alzheimer’s disease, respectively (32). In line with previous observations (32, 33, 39), mutation of the key glycine residue in this motif abolishes TelE activity. Since the glycine residues of the motif are usually positioned in the inner part of membrane pores, we speculate that change of the small glycine to a bulkier valine residue may affect the oligomerization and/or pore opening of TelE, as demonstrated for other glycine zipper toxins (32).

In addition to TelE, we discovered two additional yet to be characterized T7SSb effectors namely Gallo_0559 and Gallo_0560 located immediately upstream of TelE and probably encoded in the same operon. Bioinformatic analysis showed that Gallo_0559 is a DUF5082 family protein whereas Gallo_0560 is a TIGR01497 family protein. Proteins belonging to these two families were previously identified in *S. intermedius* as WxgA and WxgC, respectively (18). WxgA and WxgC are chaperone proteins essential for the secretion of LXG family proteins (Tel) in *S. intermedius* (18). These proteins were suggested as WXG100-like proteins (18), but WxgA and another chaperone protein WxgB are 40% longer than the canonical WXG100 proteins. These chaperones have not been detected in the *S. intermedius* bacterial supernatant and were henceforth hypothesized to disassociate from Tel proteins prior to or during secretion (18). Here we found that these probable Wxg-like chaperones were detected in the supernatant of *SGG* UCN34 WT but not in the Δ*essC* mutant, indicating that the secretion of these proteins is T7SSb-dependent, and that they may remain associated with LXG toxins post secretion. Further investigation on the potential interaction of these two proteins with TelE may provide insight into T7SSb effectors secretion and delivery. Klein et al. showed that one or two LXG-associated proteins located upstream of LXG-effector are involved in the correct targeting of the LXG effector to the Type VII secretion machinery.

The three LXG family toxins identified in this study expands our knowledge on the antibacterial arsenal possessed by *SGG*. It is noteworthy that *SGG* also produces bacteriocins such as gallocin A exhibiting antimicrobial activity against gut commensals such as *E. faecalis* (40), which together with the newly identified Tel toxins, are likely the arsenal essential for *SGG* survival in the highly dynamic and populated human gut environment.

Finally, our findings may have further implications in understanding the pathogenicity of *SGG*. Indeed, it was shown in *SGG* strain TX20005 that T7SSb is a key component contributing to *SGG* gastrointestinal tract colonization and cancer-promoting capability (30). Of note, TX20005 belongs to the small minority of *SGG* isolates displaying a different T7SSb gene locus arrangement together with a different repertoire of effectors as compared to UCN34, and the impact of these differences on the pathogenic outcome is currently unclear. The contribution of pathogenicity associated islands in bacterial virulence is well established. For instance, the CagA toxin, VacA toxin and Type IV secretion system encoded in the *Helicobacter pylori* pathogenicity island (41, 42), or the colibactin synthase genes encoded in the *E. coli* polyketide synthase (*pks*) genotoxic island (43-45) are key to the progression of gastric cancer and colorectal cancer, respectively. Of particular interest is VacA that also contains a glycine zipper motif essential for its function causing ulceration and gastric lesions in murine model (46). Whether TelE could also alter host cells is a perspective for future studies.

## MATERIALS AND METHODS

### Bacterial culture conditions

All bacterial strains and plasmids used in this study are listed in **Table S1**. Cultures of *E. coli* strains were prepared from single colonies in lysogeny broth (LB) (BD) and incubated overnight at 37°C, 200 rpm, whereas cultures of *Streptococcus agalactiae* and *Streptococcus gallolyticus* strains were prepared from single colonies in brain heart infusion broth (BHI) (Acumedia) and incubated statically overnight at 37°C. When the use of growth medium with minimal amount of peptides were necessary, overnight culture of *SGG* were prepared from single colonies in M9YEG broth (1X M9 minimal salts supplemented with 0.5% yeast extract and 1.0% of glucose) (47) and incubated statically at 37°C. When appropriate, antibiotics were supplemented at the following concentrations: kanamycin, 50 µg/mL for Gram-negative bacteria and 500 µg/mL for Gram-positive bacteria; gentamicin, 20 µg/mL for *E. coli*; erythromycin, 2 µg/mL for Gram-positive bacteria.

### Genetic techniques

Standard molecular cloning procedures as previously described (48) were followed to construct plasmids for gene deletion and gene expression. All primers used in this study were synthesized by Integrated DNA Technology and listed in **Table S1**. The enzymes Q5 DNA polymerase and restriction enzymes were obtained from New England Biolabs, whereas T4 ligase was obtained from Thermo Fisher Scientific. All genetic constructs were delivered into chemically transformed *E. coli* DH5⍰ prepared as described (49) and verified by Sanger sequencing performed at 1st BASE. For gene expression plasmids construction, the corresponding open reading frames with the native ribosomal binding sites were cloned downstream of inducible promoters such as P_tetO_ (anhydrotetracycline-inducible) and P_BAD_ (arabinose-inducible).

To construct plasmids for gene deletion, two ∼500bp DNA fragments corresponding to the upstream and downstream region of the gene of interest were first PCR amplified. The resulting two DNA fragments were subsequently spliced by overlap extension PCR and ligated into pG1 plasmid at *Sma*I restriction site. The constructed plasmids were transformed into *E. coli* and later transformed into *S. gallolyticus* as described in the transformation section below. Subsequently, *SGG* in-frame chromosomal deletion mutants were generated by allelic exchange as previously described (50).

To generate plasmids encoding TelE with a specific residue substitution, *telE* was PCR-amplified into two fragments corresponding to the 5’ and 3’ regions by using primers with the desired mutation incorporated. These two fragments were spliced into a full-length *telE* open reading frame by overlap extension PCR and ligated into pTCV*erm*- P_tetO_ at the *Bam*HI*/Sph*I restriction site.

Plasmid encoding TelE with the C-terminus fused with superfolder GFP was constructed by splicing the PCR amplicons corresponding to the *telE* open reading frame without a stop codon and the superfolder GFP open reading frame that has been codon-optimized for *SGG* (gene synthesized at GenScript).

### Transformation

*E. coli* DH5⍰ was chemically transformed as described in (49). *E. faecalis* and *S. agalactiae* were transformed by electroporation as previously described (51-53). *S. agalactiae* carrying the respective plasmids were used for conjugal transfer into *SGG* as described previously (50). *SGG* was also naturally transformed using a newly established method, by supplementing the log-phase culture grown in M9YEG with 10 µM of the competence inducing peptide (ITGWWGL) outlined in the previous study (54), at least 5 µg of plasmid DNA, 10 µM of magnesium chloride, 10 µM of magnesium sulfate and 1 µM of ferrous (II) chloride. The mixture was incubated for an hour at 37°C, followed by three hours at 30°C, before being plated onto BHI agar supplemented with erythromycin.

### Genomic DNA isolation

*S. gallolyticus* genomic DNA was extracted using either Wizard Genomic DNA Purification Kit or Wizard SV Genomic DNA Purification Kit (Promega). The extraction was performed according to the manufacturer’s instructions, except that 20U to 50U of mutanolysin (Sigma-aldrich) was added at the lysozyme treatment step. When necessary, an additional phenol:chloroform clean up step was performed to remove the excessive impurities in the samples.

### Genomic DNA sequencing and assembly

High integrity DNA were sequenced to at least a coverage of 100X on the Illumina Miseq platform (reagent kit v3, 2 x 300bp) at the in-house next-generation sequencing facility in the Singapore Centre for Environmental Life Sciences Engineering. The paired-end sequencing reads were trimmed and assembled *de novo* into contigs on the CLC Genomics Workbench 10.0 (Qiagen). These assembled contigs were subsequently uploaded to the RAST server (55-57) for open reading frame prediction and gene annotation.

### Phylogenetic analysis

A core genome-based alignment was performed on Parsnp version 1.2 integrated in the Harvest suite (58), with *SGG* UCN34 (GenBank accession no. FN597254) served as the reference genome. The other *S. gallolyticus* complete closed genomes available on NCBI, *i.e. SGG* ATCC43143 (GenBank accession no. NC_017576), ATCC BAA_2069 (GenBank accession no. NC_015215) and DSM 16831 (GenBank accession no. NZ_CP018822); *SGP* ATCC43144 (GenBank accession no. NC_015600), NCTC13784 (GenBank accession no. NZ_LS483462) and WUSP067 (GenBank accession no. NZ_CP039457); and two additional draft genomes – *SGM* ACA-DC-206 (unpublished genome) and the *SGG* TX20005 (NCBI Assembly ID GCA_000146525) - were also included in the analysis for a clear distinction of these closely related strains. The resulted alignment was subsequently visualized on Gingr (integrated in the Harvest suite) and used for phylogenetic inference on FigTree2 based on approximate-maximum-likelihood algorithm (59).

### Comparative genomics

A genome-wide comparison among *SGG* UCN34, three representative *SGG* clinical isolates, and three representative *SGP* was performed using BLAST Ring Image Generator v0.95 based on BLASTn search (60). Genomic islands of the *SGG* UCN34 were downloaded from IslandViewer 4(61). NCBI tBLASTn program was used to identify the protein homologs of the T7SS core components across all the *SGG* and *SGP* clinical isolates. Easyfig v2.2.2 was used to visualize the region of interest and to perform pairwise comparisons on the protein sequences using tBLASTx program (62).

### Comparative proteomics on the *SGG* secretomes

Single colonies of *SGG* UCN34 WT and the Δ*essC* mutant were inoculated into 20 mL of M9YEG and grown to OD_600nm_ 3.0 corresponding to the early stationary phase of growth. The culture supernatant was collected and filtered through 0.22 µm filter units. 15 mL of the filtered culture supernatant was first concentrated using Amicon Ultra 15 mL Centrifugal Filters (cut-off 10 kDa, Merck), and the proteins were precipitated out of the solution using trichloroacetic acid (TCA) as previously described (63). The samples were further processed and analyzed at the Taplin Mass Spectrometry Facility. Briefly, precipitated proteins were first treated with trypsin (Promega) in 50 mM ammonium bicarbonate solution, dried in a speed-vac and were subsequently reconstituted in HPLC solvent A (2.5% acetonitrile, 0.1% formic acid). Peptides were loaded using a Famos auto sampler (LC Packings) into a nano-scale reverse-phase HPLC fused silica capillary column (100 µm inner diameter x ∼30 cm length) packed with 2.6 µm C18 spherical silica beads and eluted with increasing concentrations of solvent B (97.5% acetonitrile, 0.1% formic acid). Eluted peptides were subjected to electrospray ionization and entered into an LTQ Orbitrap Velos Pro ion-trap mass spectrometer (Thermo Fisher Scientific). Peptides were detected, isolated, and fragmented to produce a tandem mass spectrum of specific fragment ions for each peptide. Peptide sequences (and hence protein identity) were determined by comparing the fragmentation pattern acquired by the software program Sequest (Thermo Fisher Scientific) (64) to the protein sequences on protein databases. All databases include a reversed version of all the sequences and the data was filtered to between a one and two percent peptide false discovery rate.

Downstream data analysis was performed in R studio using the Differential Enrichment analysis of Proteomics Data 1.10.0 package (65). Proteins that were not detected across all the three replicates of either the UCN34 WT or the Δ*essC* mutant were excluded from the analysis. Next, the total peptide count of the remaining proteins (n = 162) was scaled and variance stabilized (66) before the missing values (*i.e.* peptide count = 0) were imputed with the assumption that the missing values are missing not at random (MNAR), using the Deterministic Minimum Imputation (MinDet) strategy (10^-4^ quantile) (67). The MinDet-imputed data were subsequently used for differential expression analysis, with a cut off of false discovery rate of 5%, and a log_2_ fold change of 1. The derived data set was plotted using the EnhancedVolcano package (68).

### Bacterial toxicity experiments

For an intracellular expression of the *SGG* T7SSb effectors, *E. coli* DH5⍰ or GM48 were transformed with pTCVerm-P_tetO_ plasmid series (see **Table S1** for the plasmids used in this study). Overnight cultures of these strains were serially diluted in tenfold increments and 5 µL of each dilution was spotted onto LB agar supplemented with an appropriate amount of antibiotics and inducer (kanamycin, 50 µg/mL; anhydrotetracycline, 100 to 500 ng/mL). Agar plates incubated overnight at 37°C were documented on the Gel Doc XR+ Documentation System (Bio-Rad).

For the identification of the TelE cognate immunity protein, the *E. coli* DH5⍰ carrying pTCVerm-P_tetO_::TelE was transformed with compatible plasmids of either pJN105 (vector control) or pJN105 containing each individual open reading frames found downstream of *telE*. The expression of these open reading frames was driven by arabinose-inducible promoter (P_BAD_). Overnight cultures of these strains were first 50x-diluted into sterile LB broth supplemented with antibiotics and incubated at 37°C with agitation at 200 rpm for 1.5 hours. These bacterial cultures at the early log-phase of growth were supplemented with arabinose to a final concentration of 0.5%, and incubated further for 3 hours, before being subjected to serial dilution and spotted onto LB agar supplemented with an appropriate amount of antibiotics and inducer (kanamycin, 50 µg/mL; gentamicin, 20 µg/mL; anhydrotetracycline, 100 ng/mL; arabinose, 0.5%). Agar plates incubated overnight at 37°C were documented on the Gel Doc XR+ Documentation System (Bio-Rad).

### *In silico* TelE prediction and homologs identification

The TelE amino acid sequence from the reference strain *SGG* UCN34 (protein ID CBI13054.1) was uploaded onto the web-based I-TASSER server (69-71) for protein structural prediction using the default settings. The same amino acid sequence was also used to identify TelE homologs among the clinical isolates using the NCBI tBLASTn program. Hit proteins were considered as TelE homologs if significant similarities (e-value <10^-11^) were identified at the N terminus, C terminus, or the full-length protein.

### Multiple sequence alignment (MSA) and sequence logo generation

MSA was performed on the online MAFFT server version 7 (https://mafft.cbrc.jp/alignment/server/) with E-INS-i iterative refinement method that assumes all the sequences shared the same conserved motifs (72, 73). Regional sequence logo (probability-weighted Kullback-Leibler logo type) was generated on web-based Seq2Logo 2.0 server (http://www.cbs.dtu.dk/biotools/Seq2Logo/), with the Heuristics clustering method applied (74).

### Fluorescence microscopy imaging

For time-lapse microscopy imaging, log-phase *E. coli* (OD_600nm_ ∼ 1.0 Abs) cultured in LB was mixed with propidium iodide (1:1000 diluted from 20 mM stock solution, Thermo Fisher Scientific) and anhydrotetracycline at a final concentration of 500 ng/mL (Sigma Aldrich), and spotted onto an 1% (w/v) low melting agarose pad (bioWORLD). Samples were immediately imaged with the microscope Axio Observer 7 fitted with Plan Apochromat 40x/0.95 dry lens for phase contrast imaging, or Plan Apochromat 100x/1.4 oil lens for differential interference contrast imaging (Carl Zeiss), with temperature maintained at 37°C, and aided with Definite Focus.2 for autofocus. When necessary, samples were excited with filter wheel 555/30 to detect red fluorescence signals.

To image *E. coli* expressing TelE-sfGFP with or without TipE, overnight cultures were diluted 50x into sterile LB broth, incubated at 37°C with 200 rpm for 1.5 hours to grow to early log-phase. Samples were first induced with 0.5% arabinose for 1.5 hours and were subsequently supplemented with anhydrotetracycline to a final concentration of 500 ng/mL and propidium iodide (1:1000 diluted from 20 mM stock solution, Thermo Fisher Scientific), and incubated further for 2 hours at 37°C without agitation. Following incubation, samples were spotted onto an 1% low melting agarose pad (bioWORLD), and imaged with the microscope Axio Observer Z1 fitted with Plan Neofluar 100x/1.3 oil lens, with excitation at 488 nm (GFP) and 535 nm (PI).

### Flow-cytometry

To analyze TelE-gfp expression, 3 mL of bacterial culture in LB medium was prepared (initial OD around 0.1) from fresh agar plates and grown until reaching an OD_600_ = 0.4-0.5. At that time, we removed an aliquot of 500 µL where no inducer was added (non-induced sample or NI). Anhydrotetracycline at 200 ng/mL was added to the rest of the culture and 500 µL aliquots were removed at various time points from 30 to 90 min, and Hoescht 33342 was directly added to the medium (1/1000 dilution), the tubes were left open and incubated 10 min under gentle agitation. After washing and resuspension in PBS, samples were acquired on a MACSQuant YGV Analyzer Apparatus (Milteyni Biotec) and data were analyzed using FlowJo^TM^10 software. GraphPad Prism 9 was used for statistical analysis.

### Protein expression, purification and pull-down

Protein samples were in general prepared as follow: overnight cultures of *E. coli* BL21(DE3) harboring the respective protein expression pET28b plasmids were 50x diluted into sterile LB broth supplemented with 50 µg/mL of kanamycin and grown to an optical density of ∼0.7 Abs measured at 600 nm. Subsequently, cultures were added with isopropyl β-D-1-thiogalactopyranoside (IPTG) to a final concentration of 100 µM and incubated for an additional 3 hours at 37°C, 200 rpm. Following incubation, cells were harvested by centrifugation at 8,000 xg for 10 minutes, and the resulted cell pellets were stored at −80°C (for protein purification). When needed, cells were resuspended in 100 mM HEPES pH 7.5 and lysed by sonication (10s on-off pulse cycles, at 20 or 40% amplitude, Sonics Vibracell VCX750). Unbroken cells were removed by centrifugation at 8,000 xg for 10 minutes at 4°C, and the resulted supernatant was ultracentrifuged (Beckman Coulter XPN100) at 35,000 rpm for 1 hour at 4°C to separate the soluble (supernatant) and membrane proteins (pellet).

For protein purification, the membrane pellets were resuspended to a final concentration of 100 mg/mL in Buffer A (100 mM HEPES pH 7.5, 500 mM NaCl and 1% n-dodecyl-β-D-maltoside (DDM)) and incubated for 1 hour at 4°C aided with top-down rotation. The solubilized membrane proteins were subsequently supplemented with imidazole to a final concentration of 50 mM and incubated with nickel-immobilized HisLink resins (Promega) that had been pre-calibrated in Buffer B (100 mM HEPES pH 7.5, 500 mM NaCl, 50 mM imidazole and 0.1% DDM), for 30 minutes at 4°C with top-down rotation. Proteins-bound HisLink resins were washed with 30X resin volume of ice-cold Buffer B, and finally rinsed with ice-cold buffer containing 100 mM HEPES pH 7.5, 500 mM NaCl, 500 mM imidazole and 0.1% DDM to elute the proteins.

For protein pulldown, the proteins-bound HisLink resins were incubated with cellular lysate containing HA-TipE prepared in Buffer B for 20 minutes at 4°C aided with top-down rotation. The resins were subsequently washed and the bound proteins were eluted as described above.

### SDS-PAGE and Western blot

Protein samples were mixed with an equal amount of 2X Laemmli reducing buffer and boiled for 10 minutes before being subjected to SDS-PAGE analysis on 12% polyacrylamide gels. Proteins were stained with InstantBlue solution (Expedeon) for visualization or were transferred onto PVDF membranes using iBlot Dry Blotting System (Thermo Fisher Scientific) at 20V for 7 minutes. Following protein transfer, the PVDF membranes were immediately blocked with 1X casein blocking buffer (Sigma-Aldrich), and were probed with either horseradish peroxidase (HRP)- conjugated primary antibodies against His-tag (Qiagen) or with rabbit primary antibodies against HA-tag (Sigma-Aldrich) followed by anti-rabbit HRP-conjugated secondary antibodies. Membranes were thoroughly rinsed with PBS containing 0.1% Tween 20 between the incubations. Chemiluminescent signals were detected using Amersham ImageQuant 800 System (GE Healthcare) after a brief incubation of the PVDF membranes with Immobilon Forte Western Chemiluminescent HRP Substrates (EMD Millipore). For reblotting with a different antibody after image acquisition, the membranes were incubated with Restore PLUS Western Blot Stripping Buffer (Thermo Fisher Scientific) for 30 minutes at room temperature, rinsed thoroughly before being blocked with casein blocking buffer and probed with antibodies as described above.

## Supporting information

All supplemental material

## ACKNOWLEDGEMENT

We are very grateful to Alexandra Doloy and Nicolas Dmytruk for providing us with the collection of S. *gallolyticus* clinical isolates. We thank Ross Tomaino from Taplin Mass Spectrometry Facility for the mass-spectrometry analysis, Daniela Megrian-Nunez for her help with AlphaFold and Bruno Périchon for his help in the comparative analysis of T7SSb genetic locus shown in Fig.S1b. We thank Tarek Msadek for critical reading of the manuscript. This work was supported by the National Research Foundation and Ministry of Education Singapore under its Research Centre of Excellence Program (SCELSE). S. Dramsi would like to acknowledge the support of the Institut National contre le Cancer (INCA, grant PLBIO16-025) and from the French Government’s Investissement d’Avenir program, Laboratoire d’Excellence Integrative Biology of Emerging Infectious Diseases (grant no. ANR-10-LABX-62-IBEID).

