## Supplementary material for "Characterization of TelE, a T7SS LXG effector exhibiting a conserved C-terminal glycine zipper motif required for toxicity": All supplemental material

**SUPPLEMENTARY FIGURES LEGEND**

**Fig. S1a. Phylogenetic tree of the clinical isolates sequenced in this study.** Phylogenetic inference of the isolates was performed using approximate-maximum-likelihood algorithm based on the core-genome alignment using Parsnp version 1.2. Scale bar represents substitution per variant site within the core genome. The isolates sequenced in this study were labeled in black font, whereas the reference strains *SGP* (NCTC13784, ATCC43144 and WUSP067) and *SGG* (DSM16831, ATCC43143, BAA2069 and UCN34) with complete genomes available on NCBI GenBank, and a draft genome of *SGM* (strain ACA-DC-206) were labeled in blue font. Genomes used for comparative genomics analysis as shown in Figure 1A were highlighted with darker color shades. Bootstrap value for all the major branches ranges from 0.97 to 1. Genomes labelled with asterisk contain a different T7SSb gene locus arrangement than the UCN34 reference strain.

**Fig. S1b. Comparison of the T7SSb locus in several *SGG* and *SGP* clinical isolates**. (A) A pairwise nucleotide sequence comparison of the genomic region of interest. Each of the T7SSb genes (arranged in the order of *esxA, essA, esaB, essB, essC, esaA*) was represented by colored arrows. Light gray arrows represent other open reading frames in this genomic region. Gray shading shows the similarities between the sequences. (B) Summary of BLAST search using NCBI tBLASTn program, queried against the protein sequences of each T7SSb components (EsxA, EssA, EsaB, EssB, EssC, EsaA) from the reference genome UCN34. The identification of homologs was based on RAST annotation, or significant hits from the BLAST search. (C) The general gene arrangement of the T7SSb gene locus in *SGG*,

*S. intermedius* and *S. aureus*. Each T7SSb genes was color-coded and labelled accordingly. The open reading frames flanked by *esxA* and *essA* in the *S. intermedius* B196 were excluded in this figure for a clear presentation of the T7SSb gene locus.

**Figure S2**. **Detailed comparison of the whole T7SSb locus in two representative strains UCN34 and TX20005.**

**Figure S3. The prevalence of TelE homologs among *SGG* isolates.** The phylogenetic inference of the *SGG* isolates was shown on the left. Genomes in black font represent the clinical isolates sequenced in this study, whereas the genomes in blue font represent the complete genomes publicly available. Scale bar represents substitution per variant site within the core genome. Table on the right summarized the distribution of TelE homologs among *SGG* isolates. Based on the C-terminal sequence (shown in Figure 3B), the identified TelE homologs were broadly categorized into 7 TelE variants and named as TelE1 to TelE7, where TelE1 refers to the reference TelE from strain UCN34. Bootstrap values of all the major branches are 1.

**Figure S4. Multiple sequence alignment of various TelE subtypes.** Amino acid sequences of TelE1 to TelE7 subtypes were aligned using online MAFFT server version 7 with E-INS-i iterative refinement method that assumes all the sequences shared the same conserved motifs. The aligned sequences were viewed using MView (75) available on EMBL-EBI.

**Figure S5. TipE counteracts TelE toxicity.** (A) Viability of *E. coli* cells harboring an empty vector (pTCVerm-$P_{tetO}$) or a vector encoding *telE* fused with epitope tags or superfolder GFP with the expression inducible by anhydrotetracycline (aTc). Bar chart on the right showed the logarithm-transformed CFU count of the cells expressing the *telE* variants in relative to the control. Error bars represent mean + standard deviation ($n \geq 3$). (B) Growth kinetics of the *E. coli* BL21(DE3) harboring either the empty vector or a vector for IPTG-inducible expression of *telE*-His or co-expression of *telE*-His and HA-*tipE.* Arrow indicates the time point when the gene expression was induced. (C) Total protein collected from the respective lysates was resolved by SDS-PAGE and stained with Coomassie blue. I, insoluble lysate; S, soluble lysate. Arrow indicates the protein band corresponding to TelE and TipE, respectively.

**Fig. S6 An Alpha Fold Model of the TelE-TipE interaction.** A representative AlphaFold2 prediction of TelE-TipE complex showing the pLDDT confidence score (scale from 0 to 100). Based on this metric, the model is colored in blue where the per-residue confidence is high (pLDDT > 90), in cyan when good (pLDDT > 70), in yellow when low (pLDDT > 50), and in orange when very low (pLDDT < 50) (76). To better distinguish the two subunits in the complex, TipE (left side) is in the 'mesh' PyMOL representation, while TelE is depicted in the 'cartoon' form (right side). An inset on the top highlights the glycine 470 of TelE that is surrounded by TipE in the model.

#### Fig. S1a

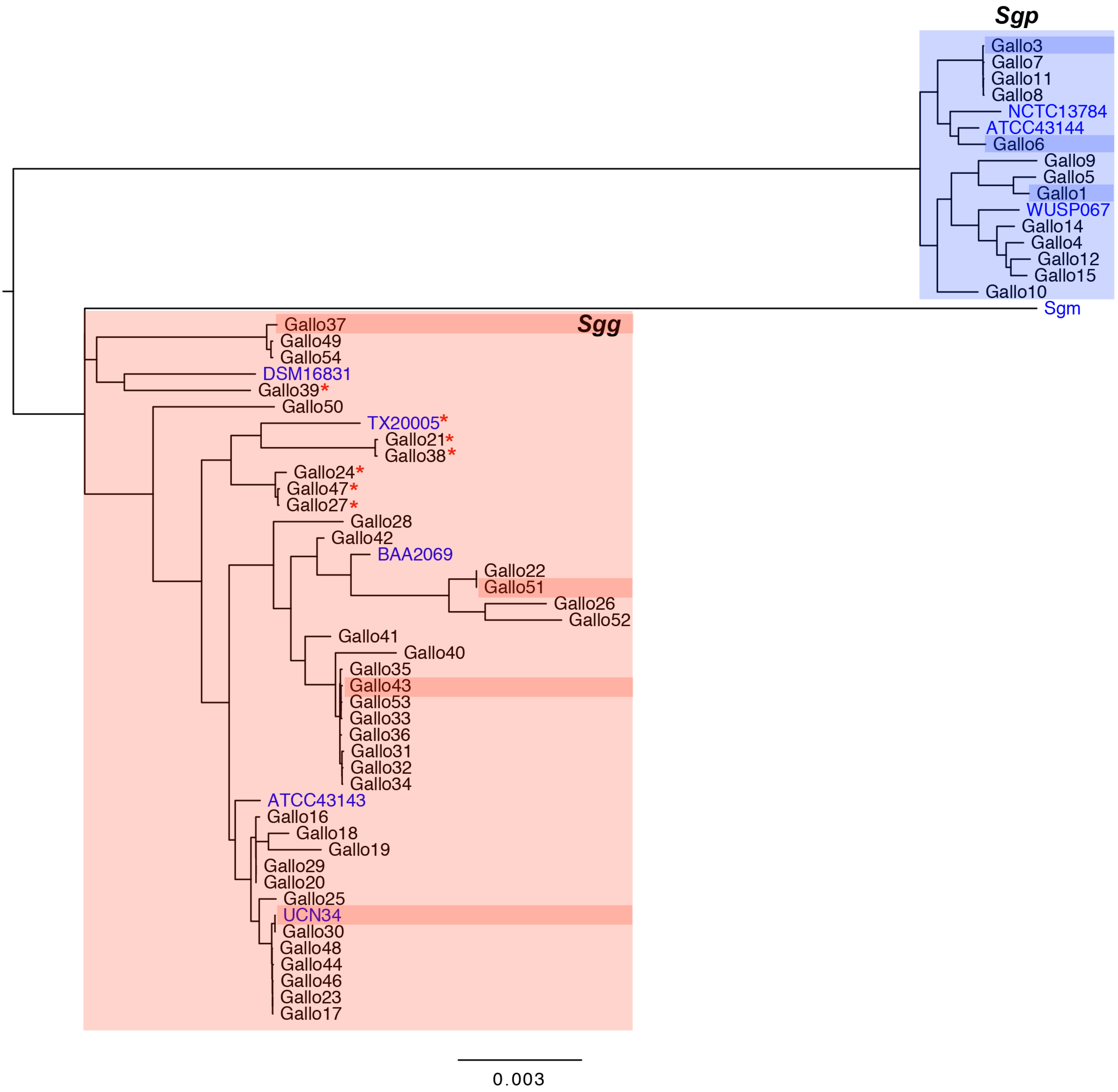
Sgp
Gallo3
Gallo7
Gallo11
Gallo8
NCTC13784
ATCC43144
Gallo6
Gallo9
Gallo5
Gallo1
WUSP067
Gallo14
Gallo4
Gallo12
Gallo15
Gallo10
Sgm
Gallo37
Sgg
Gallo49
Gallo54
DSM16831
Gallo39*
Gallo50
TX20005*
Gallo21*
Gallo38*
Gallo24*
Gallo47*
Gallo27*
Gallo28
Gallo42
BAA2069
Gallo22
Gallo51
Gallo26
Gallo52
Gallo41
Gallo40
Gallo35
Gallo43
Gallo53
Gallo33
Gallo36
Gallo31
Gallo32
Gallo34
ATCC43143
Gallo16
Gallo18
Gallo19
Gallo29
Gallo20
Gallo25
UCN34
Gallo30
Gallo48
Gallo44
Gallo46
Gallo23
Gallo17
0.003

### Figure S1b

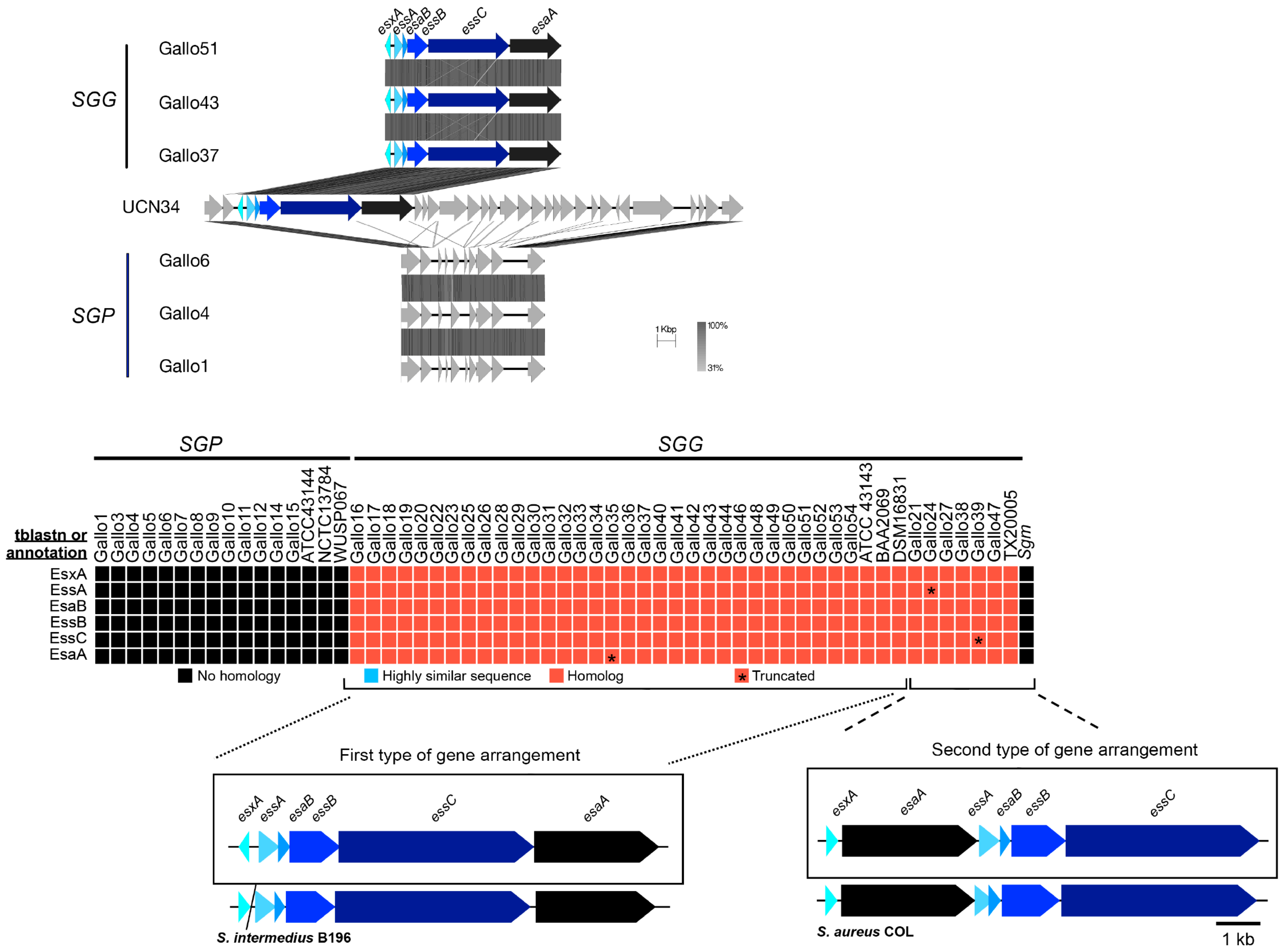
SGG
SGP
Gallo51
Gallo43
Gallo37
UCN34
Gallo6
Gallo4
Gallo1
esxA
essA
esaB
essB
essC
esaA
1 Kbp
100%
31%
tblastn or annotation
EsxA
EssA
EsaB
EssB
EssC
EsaA
Gallo1 Gallo3 Gallo4 Gallo5 Gallo6 Gallo7 Gallo8 Gallo9 Gallo10 Gallo11 Gallo12 Gallo14 Gallo15 ATCC43144 NCTC13784 WUSP067
Gallo16 Gallo17 Gallo18 Gallo19 Gallo20 Gallo22 Gallo23 Gallo25 Gallo26 Gallo28 Gallo29 Gallo30 Gallo31 Gallo32 Gallo33 Gallo34 Gallo35 Gallo36 Gallo37 Gallo40 Gallo41 Gallo42 Gallo43 Gallo44 Gallo46 Gallo48 Gallo49 Gallo50 Gallo51 Gallo52 Gallo53 Gallo54 ATCC 43143 BAA2069 DSM16831 Gallo21 Gallo24 Gallo27 Gallo38 Gallo39 Gallo47 TX20005 Sgm
No homology
Highly similar sequence
Homolog
Truncated
First type of gene arrangement
S. intermedius B196
Second type of gene arrangement
S. aureus COL
1 kb

Fig. S2

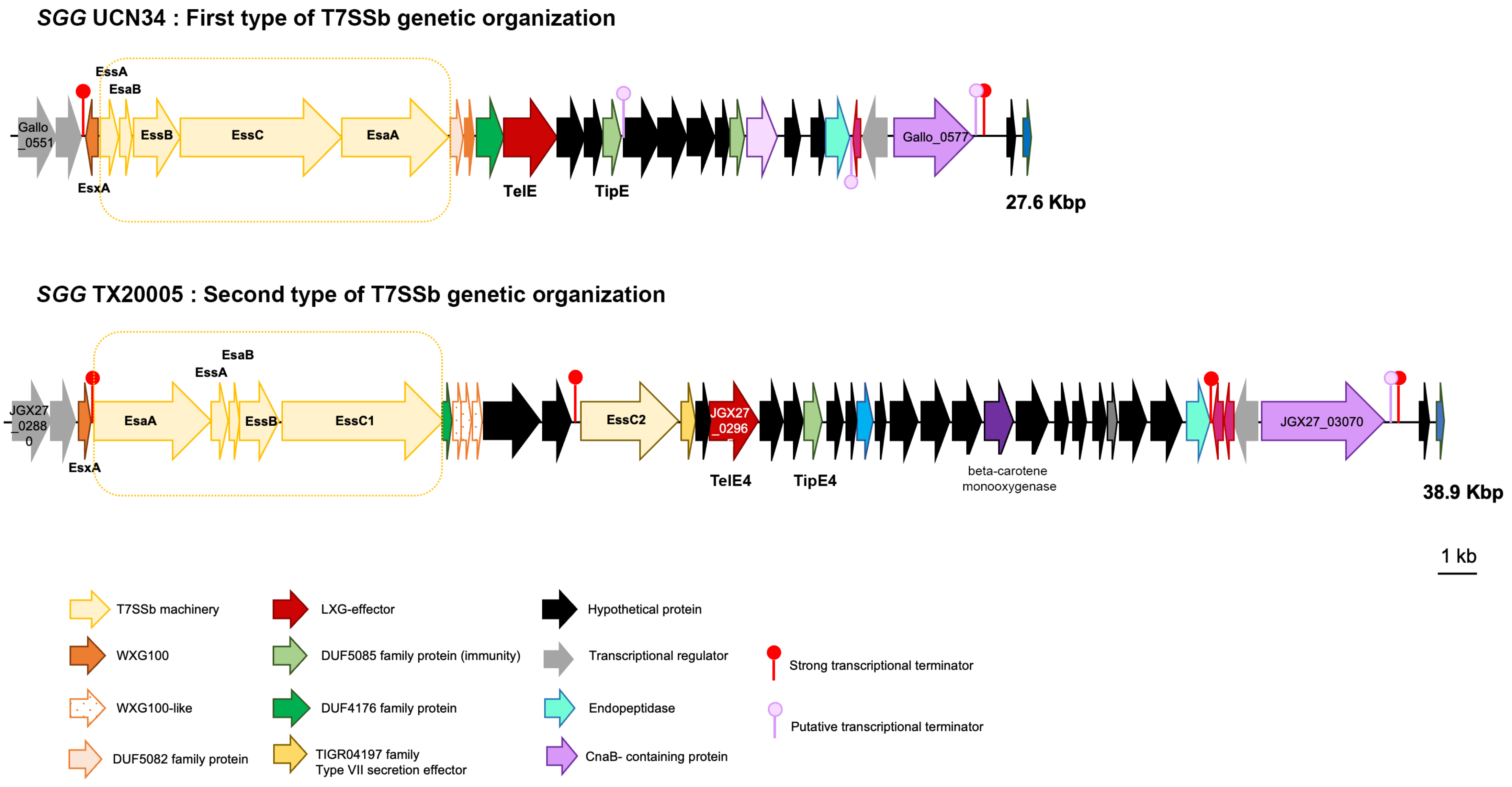
SGG UCN34 : First type of T7SSb genetic organization
Gallo_0551
EssA
EsaB
EsxA
EssB
EssC
EsaA
TelE
TipE
Gallo_0577
27.6 Kbp
SGG TX20005 : Second type of T7SSb genetic organization
JGX27_02880
EsxA
EsaA
EssA
EsaB
EssB
EssC1
EssC2
JGX27_0296
TelE4
TipE4
beta-carotene monooxygenase
JGX27_03070
38.9 Kbp
1 kb
T7SSb machinery
WXG100
WXG100-like
DUF5082 family protein
LXG-effector
DUF5085 family protein (immunity)
DUF4176 family protein
TIGR04197 family Type VII secretion effector
Hypothetical protein
Transcriptional regulator
Endopeptidase
CnaB- containing protein
Strong transcriptional terminator
Putative transcriptional terminator

#### Figure S3

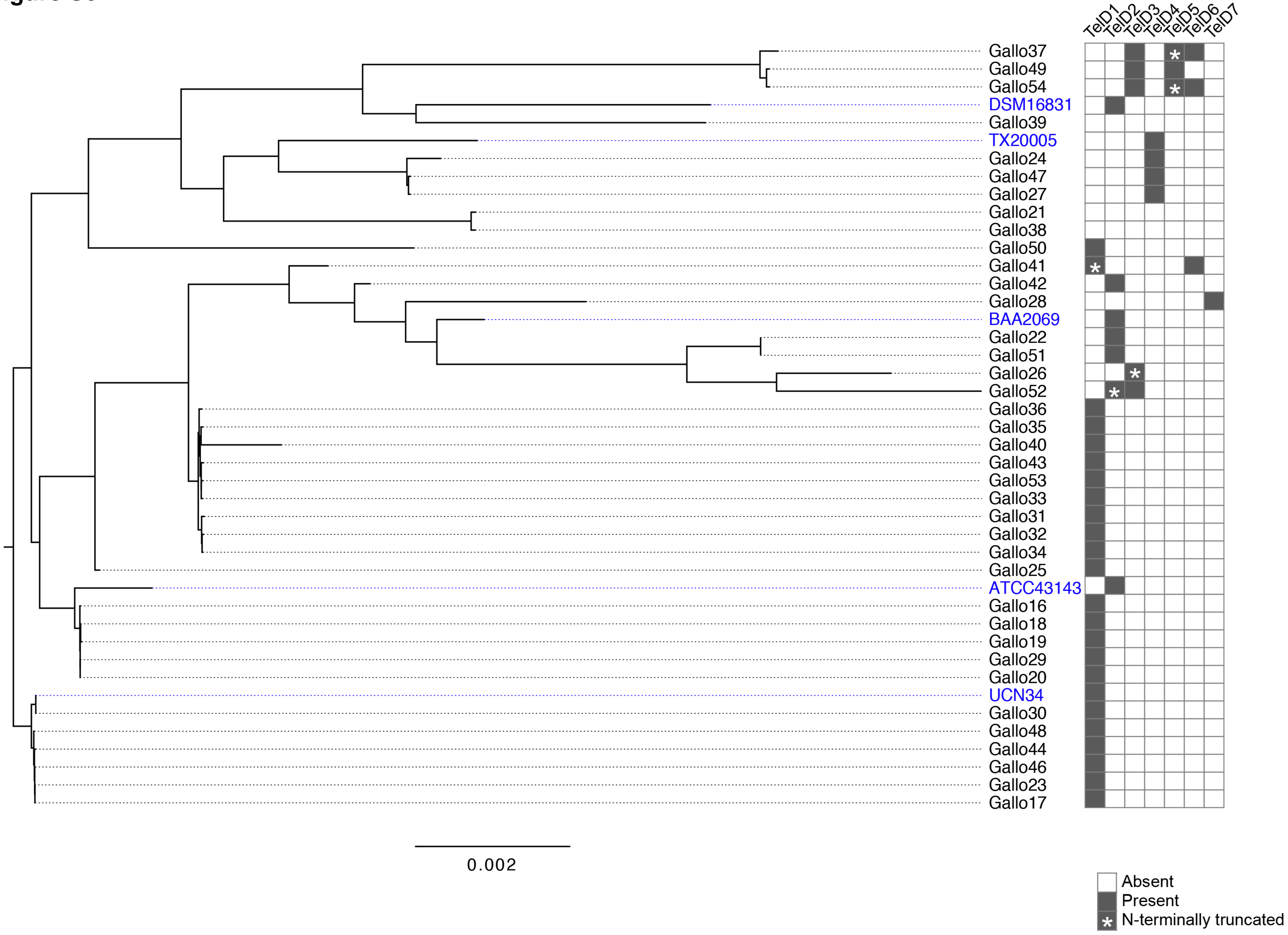
TelD1
TelD2
TelD3
TelD4
TelD5
TelD6
TelD7
Gallo37
Gallo49
Gallo54
DSM16831
Gallo39
TX20005
Gallo24
Gallo47
Gallo27
Gallo21
Gallo38
Gallo50
Gallo41
Gallo42
Gallo28
BAA2069
Gallo22
Gallo51
Gallo26
Gallo52
Gallo36
Gallo35
Gallo40
Gallo43
Gallo53
Gallo33
Gallo31
Gallo32
Gallo34
Gallo25
ATCC43143
Gallo16
Gallo18
Gallo19
Gallo29
Gallo20
UCN34
Gallo30
Gallo48
Gallo44
Gallo46
Gallo23
Gallo17
0.002
Absent
Present
N-terminally truncated

### Figure S4

```
             cov     pid   1 [        .         .         .         .         :         .         .         .         .        1 100
1 TelE1 100.0% 100.0%     M-KIKMSEVLSQKESLASSISSAKSQLESASRKLKSTADSDALQGNVKEAIDNKITNYQVPLLTNYVNSLAVISQGYDNLISTFKETVSETSDSAIIDTD
2 TelE2 100.0%  95.5%     M-KIKMSEVLSQKESLASSISSAKSQLESASRKLKSTADSDALQGNVKEAIDNKITNYQVPLLTNYVNSLAVISQGYDNLISTFKETVSETSDSAIIDTD
3 TelE3  99.6%  95.1%     M-KIKMSEVLSQKESLASSISSAKSQLESASRKLKSTADSDALQGNVKEAIDNKITNYQVPLLTNYVNSLAVISQGYDNLISTFKETVSETSDSAIIDTD
4 TelE4  88.3%  28.6%     MVKMSLGNSTSQGSSTSSISSSRISAYNGAISSLNSFIGASNLQGDAYSSAKSYASTVLIPLIQGAI----LLSEAVSNAVTTFPNRYTSEVAGESLDSE
5 TelE5  99.4%  95.3%     M-KIKMSEVLSQKESLASSISSAKSQLESASRKLKSTADSDALQGNVKEAIDNKITNYQVPLLTNYVNSLAVISQGYDNLISTFKETVSETSDSAIIDTD
6 TelE6  95.3%  57.8%     M-KIKMSEVLSQKESLASSISSAKSQLESASRKLKSTADSDALQGNVKEAIDNKITNYQVPLLTNYVNSLAVISQGYDNLISTFKETVSETSDSAIIDTD
7 TelE7  95.7%  52.1%     M-KIKMSEVLSQKESLASSISSAKSQLESASRKLKSTADSDALQGNVKEAIDNKITNYQVPLLTNYVNSLAVISQGYDNLISTFKETVSETSDSAIIDTD

             cov     pid 101          .         .         .         .         :         .         .         .         .        2 200
1 TelE1 100.0% 100.0%     VLQQMVDKFDSPLEQLKT-CSDEINKAIDEVADIV--------SLTKVT----TADAISEFKSAKKVLTNTIKDMGIFNSTVFTSEAGDILEEQNTQLSS
2 TelE2 100.0%  95.5%     VLQQMVDKFDSPLEQLKT-CSDEINKAIDEVADIV--------SLTKVT----TADAISEFKSAKKVLTNTIKDMGIFNNTVFTSEAGDILEEQNTQLSS
3 TelE3  99.6%  95.1%     ILQQMVDKFDSPLEQLKT-CSDEINKAIDEVADIV--------SLTKVT----TADAISEFKSAKKVLTNTIKDMGVFNSTVFTSEAGDILEEQNTQLSS
4 TelE4  88.3%  28.6%     VLEAQIATYQAAYDRSSAWLTSEMSKKVLNESSILRAQQSMARNLGKVTELQEKLDKLMAFNASSPSLFDEISGL--------YSAVSQGLSQVNNSFSS
5 TelE5  99.4%  95.3%     VLQQMVDKFDSPLEQLKT-CSDEINKAIDEVADIV--------SLTKVT----TADAISEFKSAKKVLTNTIKDMGIFNSTVFTSEAGDILEEQNTQLSS
6 TelE6  95.3%  57.8%     VLQQMVDKFDSPLEQLKT-CSDEINKAIDEVADIV--------SLTKVT----TADAISEFKSAKKVLTNTIKDMGIFNSTVFTSEAGDILEEQNTQLSS
7 TelE7  95.7%  52.1%     ILQQMVDKFDSPLEQLKT-CSNEINKAIDEVADIV--------SLTKVT----TADAISEFKSAKKVLTNTIKDMGIFNNTVFTSEAGDILEEQNTQLSS

             cov     pid 201          .         .         .         .         :         .         .         .         .        3 300
1 TelE1 100.0% 100.0%     LSDLGSSSYTSKKAKEFYKDSTFKAEVKEVKSVVNGKARRDQLKNALAKNLYDSKYSGYMLETSGLEVKAGILITS------------------------
2 TelE2 100.0%  95.5%     LSDLGSSSYTSKKAKDFYKDSTFKAEVKEVKSVVNGKARRDQLKNALAKNLYDSKYSGYMLETSGLEVKAGILITS------------------------
3 TelE3  99.6%  95.1%     LSDLGSSSYTSKKAKDFYKDSTFKAEVKEVKSVVNGKARRDQLKNALAKNLYDSKYSGYMLETSGLEVKAGILITS------------------------
4 TelE4  88.3%  28.6%     YN--GTFSLPSQEELAWTTTITTAWEEREANS---------------ASEEFDETIAN-IAEESGLTVEEVMAYVLY-----------------------
5 TelE5  99.4%  95.3%     LSDLGSSSYTSKKAKEFYKDSTFKAEVKEVKSVVNGKARRDQLKNALAKNLYDSKYSGYMLETSGLEVKAGILITS------------------------
6 TelE6  95.3%  57.8%     LSDLGSSSYTGKKAKDFYKDSTFKAEVKEVKSVVNGKARRDQLKNALAKNLYDSKYSGYMMETS---LPTPIMLTL------------------------
7 TelE7  95.7%  52.1%     LSDLGSSSYTSKKAKDFYKDSTFKAEVKEVKSVVNGKARRDQLKNALAKNLYDSKYSGYMMEQTRLEVTVASVTTPYLVGRKYYKKAKKWATSAYAEHRF

             cov     pid 301          .         .         .         .         :         .         .         .         .        4 400
1 TelE1 100.0% 100.0%     ---------TKKSIGKGKKVVKILDA-TDKALKQK-K------FIGGRELM------------VDSKGRVKVGERF------LYKKPSKGKTKT----VH
2 TelE2 100.0%  95.5%     ---------TKKSIGKGKKVVKILDA-TDKALKQK-K------FIGGRELT------------VDSKGRVKVGERF------LYKKPSKGKTKT----VH
3 TelE3  99.6%  95.1%     ---------TKKSIGKGKKVVKILDA-TDKALKQK-K------FIGGRELT------------VDSKGRVKVGERF------LYKKPSKGKTKT----VH
4 TelE4  88.3%  28.6%     ---------GKKPISYTKKAQKILAT-AIKYQKGK-A------FLKGKEIT------------VEK----------------VYAGTSRESTRVRWNGKP
5 TelE5  99.4%  95.3%     ---------TKKSIGKGKKVVKILDA-TDKALKQK-K------FIGGRELT------------VDSKGRVKVGERF------LYKKPSKGKTKT----VH
6 TelE6  95.3%  57.8%     ---------REKGSDYYKEFQKVMDNDTAKYLSEAFK------FLPSKLIK-----------------SLKQSEGFKEIIWGLAQSGEKGEKAA----TY
7 TelE7  95.7%  52.1%     SKDWMGNTIIKKSEKTVDHALDFIFG-TSKTLKKK-KSHSIVQYHGNKTVTKSFEKLMKTLGETDYSKEIRKSLKT------IEKNGLKGTSKA----KE

             cov     pid 401          .         .         .         .         :         .         .         .         .        5 500
1 TelE1 100.0% 100.0%     LYS---KGTKEYKKTVGEDF--SFKKTTLYNGRKLKANGKVGID-----WKGVGTSGKEAFKSGLKDNFKSTINPLNEFKGFKEASNFSKAGKVLGFAGT
2 TelE2 100.0%  95.5%     LYS---KGTKEYKKTVGEDF--SFKKTTLYNGRKLKANGKVGID-----WKGVGTSGKEAFKSGLKDNFKSTINPLNEFKGFKEASNFSKAGKVLGFAGT
3 TelE3  99.6%  95.1%     LYS---KGTKEYKKTVGEDF--SFKKTTLYNGRKLKANGKVGID-----WKGVGTSGKEAFKSGLKDNFKSTINPLNEFKGFKEASNFSKAGKVLGFAGT
4 TelE4  88.3%  28.6%     LYD---QETDQLFKS----------------GKDLKEASEIDISKTRYAWTAEGNINFSEMGKAGWSSFKSAIDPLSDFKGWKDASNVTKAGKVLGIAGT
5 TelE5  99.4%  95.3%     LYS---KGTKEYKKTVGEDF--SFKKTTLYNGRKLKANGKVGID-----WKGVGTSGKEAFKSGLKDNFKSTINPLNE---FKEASNFSKAGKVLGFAGT
6 TelE6  95.3%  57.8%     VL----KGLMKY-----EEFGNAIKNTKL--GTLIAGTSKVFDG-----AKKVADPIKSFVKEGLKSTKFYKAISETATAGGEVSGTLNFLGKGLKFAGK
7 TelE7  95.7%  52.1%     IAKGVLKGTKSNLISEAEE---AFNVKTVKN--VAKATKKTGQI-----TKGVKTAFKES----------------NVVTALKTGSKFTKGVAVLNVIDG

             cov     pid 501          .         .         .         .         :         .         .         .         .        6 600
1 TelE1 100.0% 100.0%     VMTIGSNIKTDFID-DRRSS------TKEKVRNFAVD-----------TTVDVVSSSSAAAAGAAVGTLVGGPVGTVVGAAVGIGFS-------------
2 TelE2 100.0%  95.5%     VMTIGSNIKTDFID-DRRSS------TKEKVRNFAVD-----------TTVDVVSSSSAAATGAAIGTLVGGPLGTVAGAIVGIAVS-------------
3 TelE3  99.6%  95.1%     VMTIGSNIKTDFID-DRKSS------TKEKVRNFAVD-----------TTVDVVSSSSAAAAGAAVGTLVGGPLGTVVGAAVGIAIS-------------
4 TelE4  88.3%  28.6%     AMTVINNVNADFINADGGIN------SAQNWGNFAVD-----------TGVDLASGAGAAAIGAAAGSLVAPPLGTVVGAGVGMLVS-------------
5 TelE5  99.4%  95.3%     VMTIGSNIKTDFID-DRRSS------TKEKVRNFAVD-----------TTVDVVSSSSAAAAGAAVGTLVGGPLGTVVGAAAGIAVS-------------
6 TelE6  95.3%  57.8%     AATV-----VTFAD-VAISG------VSGGVKEFAKTGDVGKGVIAGTFSAVKSVGPLEGATIGAAVG---GGFPGAVIGAGVGLGIYILDKTGVLD---
7 TelE7  95.7%  52.1%     VAETFDNVSKNKTQ-AKKQGLQGAEVSASVATGFAVDA---------AKATVKGVASTAAATAGAALA---EAGIGLLAGAGFTVGAPVL-ITGVAAIAI
                                                                                                 |____________________________|
                                                                                                      Glycine rich motif

             cov     pid 601          .         .         .         .    :    ] 654
1 TelE1 100.0% 100.0%     --------VAMNHRVKFLGG---------KSLTEVTK---DSLKGVAG-KFGL-
2 TelE2 100.0%  95.5%     --------EFMNTNLEILGN---------KSLTGVAK---SGLKGLAG-KFGL-
3 TelE3  99.6%  95.1%     --------KAFDHKW--GGK---------NSINDYAK---NSLKGLAG-KFGL-
4 TelE4  88.3%  28.6%     --------WGMNQDW--GGG---------KSVTTWAK---DSLKGLFN------
5 TelE5  99.4%  95.3%     --------WALNKGFFGKGK---------KSINDCAK---DGLKGLAG-KFGL-
6 TelE6  95.3%  57.8%     --------TAEDLTMKFYDKSKEVVGKAVKSVGEAIS---TTFGGIGK-ALGFG
7 TelE7  95.7%  52.1%     GALAAKGVEIADKKFNITGNIK-------STANSFIKGCGDFIKGMKGWNHGVN
```

### Figure S5

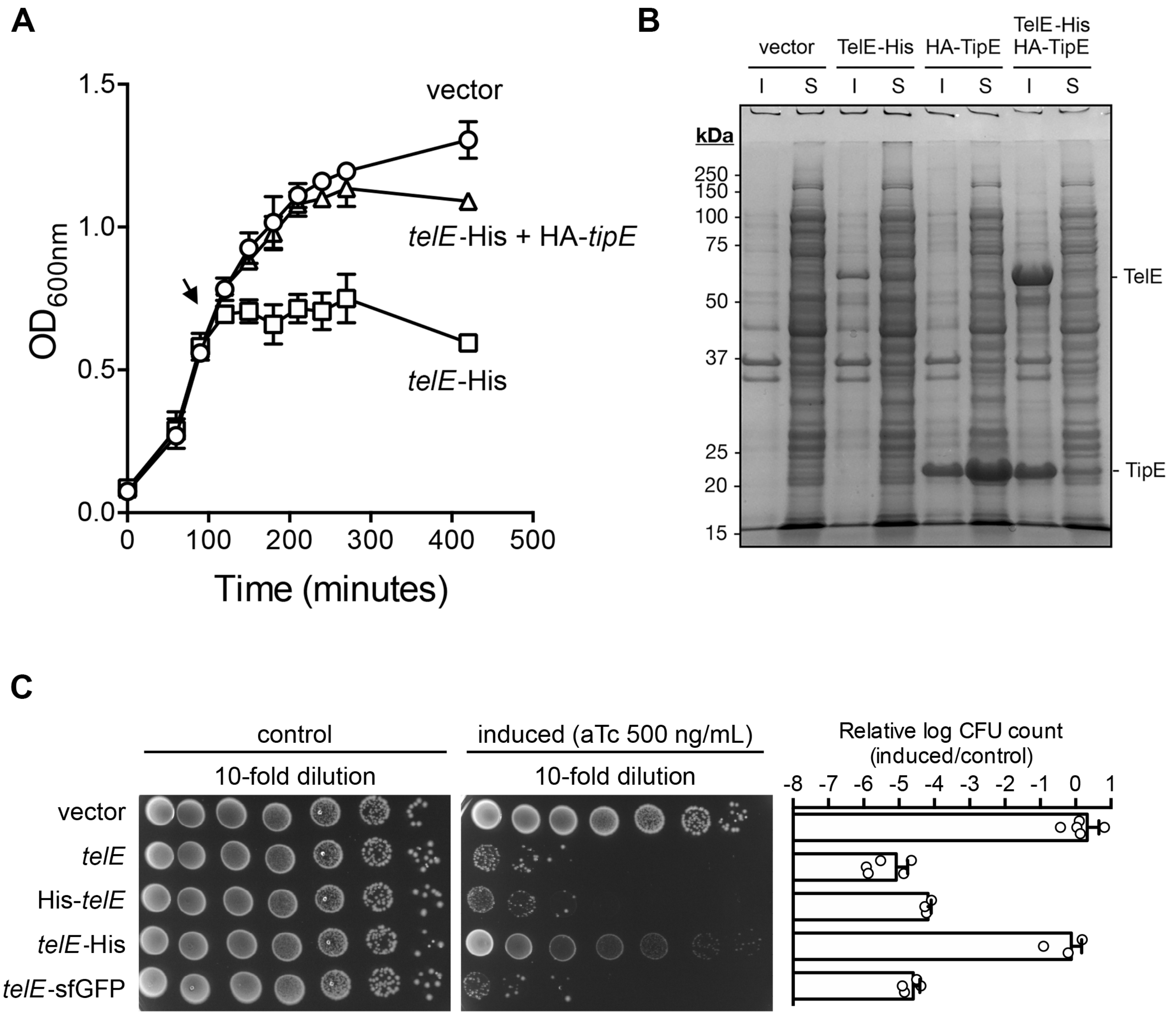
A
B
C
OD600nm
Time (minutes)
vector
telE-His + HA-tipE
telE-His
vector
TelE-His
HA-TipE
TelE-His
HA-TipE
I
S
kDa
250
150
100
75
50
37
25
20
15
TelE
TipE
control
induced (aTc 500 ng/mL)
10-fold dilution
10-fold dilution
vector
telE
His-telE
telE-His
telE-sfGFP
Relative log CFU count
(induced/control)

**Fig. S6**

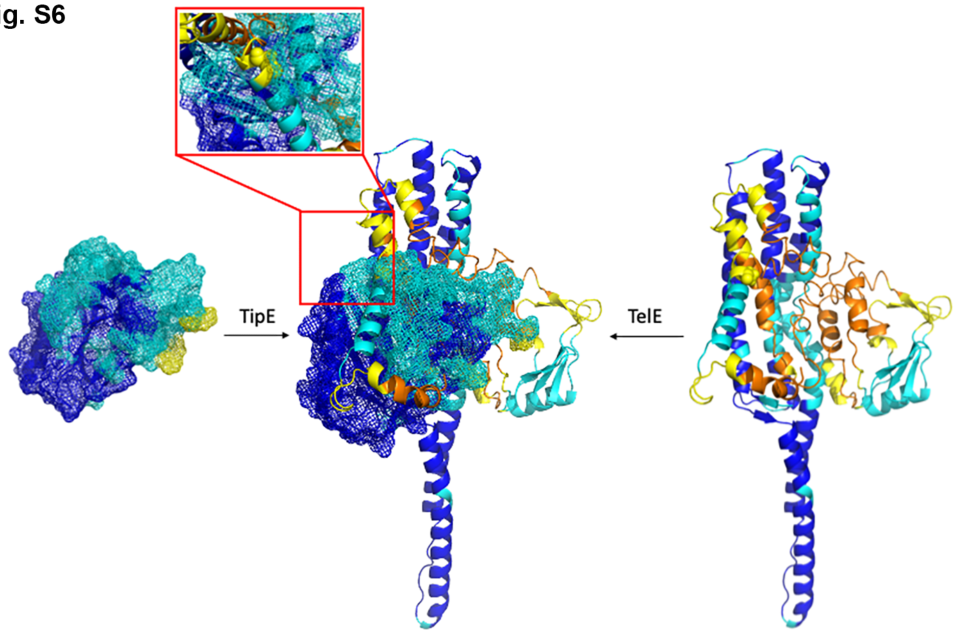

**Table S1. Strains, plasmids and primers used in this study.**

| | | |
|---|---|---|
| **Strains** | | |
| | Description | Reference |
| *Sgg* UCN34 | A clinical strain isolated from an infective endocarditis patient who later diagnosed with CRC | (1) |
| *S. agalactiae* NEM316 | MLST-23, serotype III isolated from neonate blood culture | (2) |
| *E. coli* DH5**α** | *deoR endA1 gyrA96 hsdR17* Δ(*lac*)*U169 recA1 relA1 supE44 thi-1* (ϕ80 *lacZ*ΔM15) | Lab collection |
| *E. coli* BL21(DE3) | *gal* $hsdS_B$ *ompT* | Lab collection |
| **Plasmids** | | |
| | Description | Reference |
| pG1 | Temperature sensitive vector to clone in PCR fragments at *Sma*I site; $Erm^R$ | (3) |
| pG1-*essC*KO | to generate a deletion mutant of *essC* (*gallo_0557*) in *Sgg* UCN34 | This study |
| pTCV*erm-ptetO* | *E.* coli/*Streptococcus*/*Enterococcus* shuttle vector for inducible gene expression using anhydrotetracycline; $Kan^R$ $Erm^R$ | (3) |
| pTCV*erm-ptetO-esxA* | pTCV*erm-ptetO* containing *esxA* ORF (*Gallo_0553*) at *Bam*HI / *Sph*I sites | This study |
| pTCV*erm-ptetO-gallo0559* | pTCV*erm-ptetO* containing *Gallo_0559* ORF at *Bam*HI / *Sph*I sites | This study |
| pTCV*erm-ptetO-gallo0560* | pTCV*erm-ptetO* containing *Gallo_0560* ORF at *Bam*HI / *Sph*I sites | This study |
| pTCV*erm-ptetO-telC1* | pTCV*erm-ptetO* containing *telC1* ORF (*Gallo_1068*) at *Bam*HI / *Pst*I sites | This study |
| pTCV*erm-ptetO-telC2* | pTCV*erm-ptetO* containing *telC2* ORF (*Gallo_1574*) at *Bam*HI/ *Pst*I sites | This study |
| pTCV*erm-ptetO-telE* | pTCV*erm-ptetO* containing *telE* ORF (*Gallo_0562*) at *Bam*HI / *Sph*I sites | This study |
| pTCV*erm-ptetO-telE2* | pTCV*erm-ptetO* containing *telE2* ORF (amplified from Gallo42 isolate) at *Bam*HI/ *Sph*I sites | This study |
| pTCV*erm-ptetO-telE3* | pTCV*erm-ptetO* containing *telE3* ORF (amplified from Gallo37 isolate) at *Bam*HI /*Sph*I sites | This study |
| pTCV*erm-ptetO-telE4* | pTCV*erm-ptetO* containing *telE4* ORF (amplified from Gallo47 isolate) at *Bam*HI/ *Sph*I sites | This study |
| pTCV*erm-ptetO-telE5* | pTCV*erm-ptetO* containing *telE5* ORF (amplified from Gallo49 isolate) at *Bam*HI / *Sph*I sites | This study |
| pTCV*erm-ptetO-telE6* | pTCV*erm-ptetO* containing *telE6* ORF (amplified from Gallo54 isolate) at *Bam*HI/ *Sph*I sites | This study |
| pTCV*erm-ptetO-telE7* | pTCV*erm-ptetO* containing *telE7* ORF (amplified from Gallo28 isolate) at *Bam*HI / *Sph*I sites | This study |
| pTCV*erm-ptetO-telE-G458V* | pTCV*erm-ptetO* containing *telE* ORF (*Gallo_0562*) with base substitution of G to T at nucleotide position 1373 at *Bam*HI /*Sph*I sites | This study |
| pTCV*erm-ptetO-telE-G466V* | pTCV*erm-ptetO* containing *telE* ORF (*Gallo_0562*) with base substitution of G to T at nucleotide position 1397 at *Bam*HI / *Sph*I sites | This study |

| | | |
|---|---|---|
| pTCV*erm-ptetO-telE-G470V* | pTCV*erm-ptetO* containing *telE* ORF (*Gallo_0562*) with base substitution of G to T at nucleotide position 1409 at *Bam*HI /*Sph*I sites | This study |
| pTCV*erm-ptetO-telE-G474V* | pTCV*erm-ptetO* containing *telE* ORF (*Gallo_0562*) with base substitution of G to T at nucleotide position 1421 at *Bam*HI/ *Sph*I sites | This study |
| pTCV*erm-ptetO-telE-G478V* | pTCV*erm-ptetO* containing *telE* ORF (*Gallo_0562*) with base substitution of G to T at nucleotide position 1433 at *Bam*HI / *Sph*I sites | This study |
| pTCV*erm-ptetO-telE-G480V* | pTCV*erm-ptetO* containing *telE* ORF (*Gallo_0562*) with base substitution of G to T at nucleotide position 1439 at *Bam*HI/*Sph*I sites | This study |
| pTCV*erm-ptetO-telE-His* | pTCV*erm-ptetO* containing *telE* ORF (*Gallo_0562*) with a 3' hexahistidine tag at *Bam*HI /*Sph*I sites | This study |
| pTCV*erm-ptetO-His-telE* | pTCV*erm-ptetO* containing *telE* ORF (*Gallo_0562*) with a 5' hexahistidine tag at *Bam*HI / *Sph*I sites | This study |
| pTCV*erm-ptetO-telE-SGGsfGFP* | pTCV*erm-ptetO* containing *telE* ORF (*Gallo_0562*) with a 3' *Streptococcus gallolyticus* codon-optimized superfolder GFP (SGGsfGFP) tag at *Bam*HI / *Sph*I sites | This study |
| pJN105 | *E. coli/Pseudomonas* shuttle vector for inducible gene expression using arabinose; $Gm^R$ | (4) |
| pJN105-*Gallo0563* | pJN105 containing *Gallo_0563* ORF at *Nhe*I / *Xba*I site, downstream of pBAD promoter | This study |
| pJN105-*Gallo0564* | pJN105 containing *Gallo_0564* ORF at *Nhe*I / *Xba*I site, downstream of pBAD promoter | This study |
| pJN105-*tipD* | pJN105 containing *Gallo_0565* ORF at *Nhe*I/ *Xba*I site, downstream of pBAD promoter | This study |
| pJN105-*Gallo0570* | pJN105 containing *Gallo_0570* ORF at *Nhe*I/ *Xba*I site, downstream of pBAD promoter | This study |
| pET28b | *E. coli* recombinant protein expression vector; $Kan^R$ | Lab collection |
| pET28b-*telE-His* | pET28b containing *telE* ORF without a start codon and a stop codon at *Nco*I / *Xho*I sites | This study |
| pET28b-*HA-tipD-T7-telE-His* | pET28b-*telE-His* containing N-terminally HA-tagged TipD ORF followed by a T7 promoter at *Nco*I site | This study |
| pET28b-*HA-tipD* | pET28b containing *tipD* ORF without start codon at *Nco*I */ Xho*I sites | This study |
| pET28b-*telE*$_{G470V}$-*His* | pET28b containing *telE*$_{G470V}$ ORF without a start codon and a stop codon at *Nco*I / *Xho*I sites | This study |
| **Primers** | | |
| | Sequence (5' to 3') | |
| **Knockout mutants** | | |
| essCKO_up | GATCTTACACGAGAGAC // CACCATAACGTGTACGGTGATAGTAAGTCATATTATC | This study |
| essCKO_down | GATAATATGACTTACTATCACCGTACACGTTATGGTG // GCTCACTTCAGCTTGG | This study |

| | | |
|---|---|---|
| **Gene expression in pTCV*erm-ptetO*** | | |
| BamHI/SphI_Gallo0553 | TATA**GGATCC**AGGAGATTTTTATGACC //<br>TATA**GCATGC**TTAGTTAAGTCCAAATGAAG | This study |
| BamHI/SphI_Gallo0559 | TATA**GGATCC**ATGGGAGAATGTTGTGG //<br>TATA**GCATGC**TTACATACTACCTGCTGC | This study |
| BamHI/SphI_Gallo0560 | TATA**GGATCC**TGTAAAGGGAGATTTTATGG //<br>TATA**GCATGC**TTATTTGCCTCCTTTACTAC | This study |
| BamHI/PstI_TelC1 | TATA**GGATCC**CCTCTTGCGAAAGGAG //<br>TATA**CTGCAG**TCATTAGTTGTCCTCTCC | This study |
| BamHI/PstI_TelC2 | TATA**GGATCC**CCATTTACGTTGCTAAGC //<br>TATA**CTGCAG**CTATTCTCCTACGGCTTC | This study |
| BamHI/SphI_TelE | TATA**GGATCC**GTGATGTATCCAAAGG //<br>TATA**GCATGC**TCATAATCCAAATTTACC | This study |
| BamHI/SphI_TelE2 | TATA**GGATCC**GTGATGTATCCAAAGGGAG //<br>TATA**GCATGC**TCATAATCCAAATTTCCCTGC | This study |
| BamHI/SphI_TelE3 | TATA**GGATCC**AACGATTACATTGGGAGAG //<br>TATA**GCATGC**TCATAATCCAAATTTCCCTGC | This study |
| BamHI/SphI_TelE4 | TATA**GGATCC**GAAGAGGAACGAAAGG //<br>TATA**GCATGC**TTAATTAAATAATCCTTTCAAAC | This study |
| BamHI/SphI_TelE5 | TATA**GGATCC**AACGATTACATTGGGAGAG //<br>TATA**GCATGC**TCATAACCCAAACTTACCTGC | This study |
| BamHI/SphI_TelE6 | TATA**GGATCC**AACGATTACATTGGGAGAG //<br>TATA**GCATGC**TCACCCAAAACCTAGCG | This study |
| BamHI/SphI_TelE7 | TATA**GGATCC**AACGATTACATTGGGAGAG //<br>TATA**GCATGC**TCAATTAACCCCATGGTTC | This study |
| BamHI_TelE-G458V_up | TATA**GGATCC**GTGATGTATCCAAAGG //<br>CTGCTGCTACTGCAG | This study |
| SphI_TelE-G458V_down | CTGCAGTAGCAGCAG //<br>TATA**GCATGC**TCATAATCCAAATTTACC | This study |
| BamHI_TelE-G466V_up | TATA**GGATCC**GTGATGTATCCAAAGG //<br>GGTCCAACTACTAAAG | This study |
| SphI_TelE-G466V_down | CTTTAGTAGTTGGACC //<br>TATA**GCATGC**TCATAATCCAAATTTACC | This study |

| | | |
|---|---|---|
| BamHI_TelE-G470V_up | TATA**GGATCC**GTGATGTATCCAAAGG //<br>CACAGTTACAACTGG | This study |
| SphI_TelE-G470V_down | CCAGTTGTAACTGTG //<br>TATA**GCATGC**TCATAATCCAAATTTACC | This study |
| BamHI_TelE-G474V_up | TATA**GGATCC**GTGATGTATCCAAAGG //<br>GCCACTACCACAGTT | This study |
| Sph_TelE-G474V_down | AACTGTGGTAGTGGC //<br>TATA**GCATGC**TCATAATCCAAATTTACC | This study |
| BamHI_TelE-G478V_up | TATA**GGATCC**GTGATGTATCCAAAGG //<br>CCAATAACGACTGCG | This study |
| SphI_TelE-G478V_down | CGCAGTCGTTATTGG //<br>TATA**GCATGC**TCATAATCCAAATTTACC | This study |
| BamHI_TelE-G480V_up | TATA**GGATCC**GTGATGTATCCAAAGG //<br>CTACAGAAAAGACAATAC | This study |
| Sph_TelE-G480V_down | GTATTGTCTTTTCTGTAG //<br>TATA**GCATGC**TCATAATCCAAATTTACC | This study |
| BamHI_TelE-GFP_up | TATA**GGATCC**GTGATGTATCCAAAGG //<br>CCTTTTGATAATCCAAATTTACCC | This study |
| SphI_TelE-GFP_down | GGATTATCAAAAGGTGAAGAATTG //<br>TATA**GCATGC**TTATTTATACAATTCATCCATAC | This study |
| BamHI/SphI_TelE-His | TATA**GGATCC**GTGATGTATCCAAAGG //<br>TATA**GCATGC**TCA*GTGATGATGATGATGATG*TAATCCAAATTTACCC | This study |
| BamHI/SphI_His-TelE | TATA**GGATCC**CCAAAGGGAGAGATTTATG*CATCATCATCATCATCAC*AAGATTAAGATGAGCG //<br>TATA**GCATGC**TCATAATCCAAATTTACC | This study |
| Gene expression in pJN105 | | |
| NheI/XbaI_Gallo0563 | GCAT**GCTAGC**GATAGTTTGAAAGGTGTAG //<br>GCAT**TCTAGA**CTAAAGCTCATCAATCTC | This study |
| NheI/XbaI_Gallo0564 | GCAT**GCTAGC**GAAGAGATTGATGAGC //<br>GCAT**TCTAGA**TTAGTCTACCTCCAA | This study |
| NheI/XbaI_TipD | GCAC**GCTAGC**GTATACAGTGATTTGGAG //<br>GCAC**TCTAGA**TTAATACGCATACCCAAC | This study |
| NheI/XbaI_Gallo0570 | GCAC**GCTAGC**GCAGATATTATTATTCCAG // | This study |

| | | |
|---|---|---|
| | GCAC**TCTAGA**CTACTCAAAACGAACAC | |
| Gene expression in pET28b | | |
| NcoI/XhoI_TelE-His | TATA**CCATGG**TGAAGATTAAGATGAGCGAG // TATA**CTCGAG**TAATCCAAATTTACCCGCTAC | This study |
| NcoI/XhoI_HA-TipD | GCAT**CCATGG**GC*TACCCATACGATGTTCCAGATTACGCT*GAATATACTGTAGCTGAGG // TATA**CTCGAG**TTAATACGCATACCCAAC | This study |
| NcoI_HA-TipD-T7_up | GCAT**CCATGG**GC*TACCCATACGATGTTCCAGATTACGCT*GAATATACTGTAGCTGAGG // GGATCGTTAATACGCATACCCAAC | This study |
| NcoI_HA-TipD-T7_down | GTATTAACGATCCCGCGAAATTAATAC // CTGC**CCATGG**TATATCTCC | This study |

Underlined, overlapping region for overlap extension PCR; bold, restrictive enzyme recognition sequence; italicized, epitope tag sequence

### Table S2 - Raw data of the proteome analysis comparing UCN34 WT and ΔessC mutant

| gene_locus | MWT(kDa) | Unique WT R1 | Total WT R1 | Unique WT R2 | Total WT R2 | Unique WT R3 | Total WT R3 | Unique ΔessC R1 | Total ΔessC R1 | Unique ΔessC R2 | Total ΔessC R2 | Unique ΔessC R3 | Total ΔessC R3 | Max | Sum Intensity WT R1 | Sum Intensity WT R2 | Sum Intensity WT R3 | Sum Intensity ΔessC R1 | Sum Intensity ΔessC R2 | Sum Intensity ΔessC R3 | lowest value | WT R1 | WT R2 | WT R3 | ΔessC R1 | ΔessC R2 | ΔessC R3 |
|---|---|---|---|---|---|---|---|---|---|---|---|---|---|---|---|---|---|---|---|---|---|---|---|---|---|---|---|
| Gallo_1055 | 167,69 | 67 | 864 | 69 | 821 | 75 | 1017 | 65 | 579 | 64 | 556 | 70 | 833 | 1017 | 1,60E+10 | 2,20E+10 | 2,20E+10 | 9,40E+09 | 9,40E+09 | 1,90E+10 | 9,40E+09 | 1,7 | 2,3 | 2,3 | 1,0 | 1,0 | 2,0 |
| Gallo_1413 | 59,9 | 26 | 61 | 24 | 66 | 23 | 58 | 26 | 59 | 22 | 43 | 23 | 56 | 66 | 5,10E+08 | 4,10E+08 | 3,10E+08 | 3,00E+08 | 2,50E+08 | 3,40E+08 | 2,50E+08 | 2,0 | 1,6 | 1,2 | 1,2 | 1,0 | 1,4 |
| Gallo_1412 | 59,64 | 17 | 109 | 17 | 82 | 19 | 100 | 17 | 69 | 13 | 54 | 17 | 80 | 109 | 6,50E+08 | 7,90E+08 | 7,00E+08 | 6,40E+08 | 5,10E+08 | 8,70E+08 | 5,10E+08 | 1,3 | 1,5 | 1,4 | 1,3 | 1,0 | 1,7 |
| Gallo_1057 | 171,43 | 18 | 44 | 17 | 52 | 19 | 51 | 18 | 52 | 23 | 45 | 22 | 57 | 57 | 4,50E+08 | 8,40E+08 | 5,40E+08 | 3,80E+08 | 3,70E+08 | 5,40E+08 | 3,70E+08 | 1,2 | 2,3 | 1,5 | 1,0 | 1,0 | 1,5 |
| Gallo_0479 | 81,08 | 18 | 23 | 17 | 20 | 17 | 22 | 19 | 28 | 20 | 25 | 18 | 28 | 28 | 3,60E+07 | 4,10E+07 | 3,50E+07 | 4,70E+07 | 3,20E+07 | 4,70E+07 | 3,20E+07 | 1,1 | 1,3 | 1,1 | 1,5 | 1,0 | 1,5 |
| Gallo_0558 | 102,37 | 23 | 115 | 22 | 98 | 23 | 109 | 21 | 60 | 20 | 49 | 21 | 51 | 115 | 8,80E+08 | 9,20E+08 | 7,30E+08 | 3,40E+08 | 2,90E+08 | 4,00E+08 | 2,90E+08 | 3,0 | 3,2 | 2,5 | 1,2 | 1,0 | 1,4 |
| Gallo_2047 | 34,54 | 14 | 29 | 12 | 29 | 14 | 25 | 12 | 18 | 15 | 27 | 16 | 23 | 29 | 5,80E+07 | 4,30E+07 | 2,90E+07 | 3,60E+07 | 3,00E+07 | 2,90E+07 | 2,90E+07 | 2,0 | 1,5 | 1,0 | 1,2 | 1,0 | 1,0 |
| Gallo_0814 | 45,89 | 16 | 32 | 16 | 28 | 18 | 41 | 18 | 46 | 18 | 38 | 17 | 38 | 46 | 1,40E+08 | 1,40E+08 | 1,50E+08 | 1,70E+08 | 1,70E+08 | 1,80E+08 | 1,40E+08 | 1,0 | 1,0 | 1,1 | 1,2 | 1,2 | 1,3 |
| Gallo_1577 | 85,6 | 19 | 159 | 16 | 136 | 18 | 137 | 19 | 127 | 19 | 96 | 17 | 140 | 159 | 4,40E+09 | 2,00E+09 | 2,30E+09 | 1,70E+09 | 1,40E+09 | 2,20E+09 | 1,40E+09 | 3,1 | 1,4 | 1,6 | 1,2 | 1,0 | 1,6 |
| Gallo_0019 | 47,17 | 16 | 1026 | 17 | 879 | 16 | 989 | 17 | 940 | 16 | 794 | 16 | 968 | 1026 | 1,10E+11 | 9,80E+10 | 7,90E+10 | 8,00E+10 | 5,70E+10 | 9,50E+10 | 5,70E+10 | 1,9 | 1,7 | 1,4 | 1,4 | 1,0 | 1,7 |
| Gallo_0189 | 59,99 | 12 | 17 | 9 | 12 | 14 | 17 | 12 | 12 | 11 | 14 | 13 | 17 | 17 | 1,40E+07 | 9,60E+06 | 6,80E+06 | 8,60E+06 | 8,70E+06 | 9,20E+06 | 6,80E+06 | 2,1 | 1,4 | 1,0 | 1,3 | 1,3 | 1,4 |
| Gallo_0012 | 47,45 | 17 | 60 | 15 | 50 | 15 | 56 | 16 | 58 | 17 | 50 | 17 | 59 | 60 | 7,20E+08 | 8,10E+08 | 5,00E+08 | 6,50E+08 | 4,00E+08 | 6,60E+08 | 4,00E+08 | 1,8 | 2,0 | 1,3 | 1,6 | 1,0 | 1,7 |
| Gallo_1234 | 31,73 | 13 | 40 | 12 | 33 | 15 | 35 | 12 | 36 | 14 | 31 | 13 | 37 | 40 | 2,20E+08 | 1,90E+08 | 1,30E+08 | 2,60E+08 | 1,80E+08 | 2,70E+08 | 1,30E+08 | 1,7 | 1,5 | 1,0 | 2,0 | 1,4 | 2,1 |
| Gallo_0313 | 57,38 | 11 | 22 | 11 | 15 | 9 | 19 | 10 | 16 | 11 | 16 | 9 | 18 | 22 | 5,90E+07 | 4,40E+07 | 3,30E+07 | 6,30E+07 | 3,20E+07 | 4,90E+07 | 3,20E+07 | 1,8 | 1,4 | 1,0 | 2,0 | 1,0 | 1,5 |
| Gallo_0625 | 87,81 | 15 | 19 | 14 | 16 | 13 | 15 | 14 | 19 | 12 | 13 | 15 | 21 | 21 | 2,50E+07 | 2,20E+07 | 1,60E+07 | 2,60E+07 | 1,30E+07 | 3,50E+07 | 1,30E+07 | 1,9 | 1,7 | 1,2 | 2,0 | 1,0 | 2,7 |
| Gallo_1136 | 37,24 | 15 | 89 | 14 | 58 | 16 | 73 | 14 | 60 | 16 | 54 | 13 | 72 | 89 | 5,90E+08 | 4,40E+08 | 5,10E+08 | 4,30E+08 | 3,50E+08 | 4,30E+08 | 3,50E+08 | 1,7 | 1,3 | 1,5 | 1,2 | 1,0 | 1,2 |
| Gallo_1556 | 29,45 | 15 | 44 | 13 | 37 | 15 | 49 | 13 | 36 | 14 | 35 | 14 | 42 | 49 | 4,80E+08 | 3,40E+08 | 2,70E+08 | 3,90E+08 | 3,10E+08 | 2,90E+08 | 2,70E+08 | 1,8 | 1,3 | 1,0 | 1,4 | 1,1 | 1,1 |
| Gallo_0562 | 55,19 | 13 | 34 | 12 | 42 | 16 | 39 | 0 | 0 | 0 | 0 | 0 | 0 | 42 | 2,40E+08 | 3,80E+08 | 2,40E+08 | NF | NF | NF | 2,40E+08 | 1,0 | 1,6 | 1,0 | 0,0 | 0,0 | 0,0 |
| Gallo_0737 | 56,78 | 13 | 16 | 12 | 14 | 10 | 12 | 12 | 14 | 11 | 13 | 13 | 16 | 16 | 3,00E+07 | 3,40E+07 | 2,30E+07 | 2,50E+07 | 2,20E+07 | 2,90E+07 | 2,20E+07 | 1,4 | 1,5 | 1,0 | 1,1 | 1,0 | 1,3 |
| Gallo_1197 | 99,39 | 13 | 30 | 9 | 17 | 9 | 28 | 12 | 36 | 9 | 25 | 10 | 30 | 36 | 5,40E+07 | 2,00E+07 | 2,40E+07 | 3,50E+07 | 1,80E+07 | 2,90E+07 | 1,80E+07 | 3,0 | 1,1 | 1,3 | 1,9 | 1,0 | 1,6 |
| Gallo_0757 | 78,45 | 11 | 13 | 10 | 12 | 10 | 15 | 10 | 17 | 11 | 15 | 10 | 17 | 17 | 8,90E+06 | 1,10E+07 | 7,80E+06 | 1,10E+07 | 8,00E+06 | 1,20E+07 | 7,80E+06 | 1,1 | 1,4 | 1,0 | 1,4 | 1,0 | 1,5 |
| Gallo_1632 | 83,42 | 12 | 12 | 10 | 14 | 12 | 14 | 8 | 9 | 8 | 9 | 10 | 13 | 14 | 1,00E+07 | 1,10E+07 | 1,10E+07 | 7,10E+06 | 5,60E+06 | 1,10E+07 | 5,60E+06 | 1,8 | 2,0 | 2,0 | 1,3 | 1,0 | 2,0 |
| Gallo_1995 | 41,99 | 12 | 14 | 6 | 6 | 9 | 9 | 4 | 7 | 6 | 6 | 9 | 12 | 14 | 1,50E+07 | 3,20E+06 | 8,90E+06 | 2,90E+06 | 2,40E+06 | 4,90E+06 | 2,40E+06 | 6,3 | 1,3 | 3,7 | 1,2 | 1,0 | 2,0 |
| Gallo_1574 | 62,06 | 9 | 12 | 11 | 14 | 10 | 13 | 0 | 0 | 0 | 0 | 0 | 0 | 14 | 1,70E+07 | 1,70E+07 | 1,50E+07 | NF | NF | NF | 1,50E+07 | 1,1 | 1,1 | 1,0 | 0,0 | 0,0 | 0,0 |
| Gallo_1242 | 34,62 | 9 | 159 | 11 | 127 | 12 | 152 | 12 | 139 | 13 | 130 | 11 | 134 | 159 | 1,70E+09 | 1,40E+09 | 1,10E+09 | 1,20E+09 | 1,00E+09 | 1,30E+09 | 1,00E+09 | 1,7 | 1,4 | 1,1 | 1,2 | 1,0 | 1,3 |
| Gallo_2178 | 50,74 | 11 | 27 | 10 | 33 | 9 | 23 | 9 | 28 | 10 | 27 | 9 | 27 | 33 | 8,00E+07 | 9,90E+07 | 6,00E+07 | 9,10E+07 | 8,00E+07 | 7,90E+07 | 6,00E+07 | 1,3 | 1,7 | 1,0 | 1,5 | 1,3 | 1,3 |
| Gallo_1149 | 30,73 | 10 | 12 | 8 | 9 | 10 | 11 | 10 | 12 | 10 | 14 | 6 | 10 | 14 | 2,90E+07 | 2,40E+07 | 1,90E+07 | 2,60E+07 | 2,20E+07 | 1,40E+07 | 1,40E+07 | 2,1 | 1,7 | 1,4 | 1,9 | 1,6 | 1,0 |
| Gallo_1368 | 107,12 | 10 | 12 | 8 | 10 | 7 | 9 | 9 | 13 | 9 | 9 | 12 | 14 | 14 | 3,10E+07 | 2,50E+07 | 1,70E+07 | 4,00E+07 | 1,20E+07 | 3,20E+07 | 1,20E+07 | 2,6 | 2,1 | 1,4 | 3,3 | 1,0 | 2,7 |
| Gallo_0648 | 77,56 | 10 | 11 | 9 | 10 | 7 | 7 | 7 | 7 | 6 | 6 | 7 | 9 | 11 | 1,20E+07 | 1,00E+07 | 8,20E+06 | 7,30E+06 | 4,40E+06 | 1,10E+07 | 4,40E+06 | 2,7 | 2,3 | 1,9 | 1,7 | 1,0 | 2,5 |
| Gallo_0748 | 168,43 | 8 | 8 | 8 | 8 | 7 | 9 | 5 | 6 | 8 | 10 | 8 | 8 | 10 | 6,60E+06 | 7,30E+06 | 5,90E+06 | 5,50E+06 | 9,40E+06 | 5,10E+06 | 5,10E+06 | 1,3 | 1,4 | 1,2 | 1,1 | 1,8 | 1,0 |
| Gallo_0083 | 84,01 | 6 | 7 | 5 | 5 | 6 | 7 | 12 | 13 | 5 | 7 | 7 | 7 | 13 | 3,70E+06 | 4,00E+06 | 2,80E+06 | 6,80E+06 | 1,90E+06 | 2,50E+06 | 1,90E+06 | 1,9 | 2,1 | 1,5 | 3,6 | 1,0 | 1,3 |
| Gallo_0553 | 11,08 | 11 | 443 | 13 | 360 | 11 | 402 | 5 | 14 | 5 | 10 | 5 | 8 | 443 | 1,90E+10 | 1,40E+10 | 1,50E+10 | 3,90E+07 | 1,80E+07 | 9,80E+06 | 9,80E+06 | 1938,8 | 1428,6 | 1530,6 | 4,0 | 1,8 | 1,0 |
| Gallo_0324 | 59,79 | 11 | 80 | 10 | 49 | 11 | 66 | 13 | 60 | 12 | 44 | 10 | 48 | 80 | 2,80E+08 | 2,10E+08 | 1,70E+08 | 2,40E+08 | 1,70E+08 | 1,50E+08 | 1,50E+08 | 1,9 | 1,4 | 1,1 | 1,6 | 1,1 | 1,0 |
| Gallo_1125 | 34,57 | 11 | 26 | 9 | 18 | 9 | 22 | 9 | 27 | 11 | 25 | 11 | 26 | 27 | 1,40E+08 | 8,30E+07 | 7,90E+07 | 1,00E+08 | 6,10E+07 | 9,10E+07 | 6,10E+07 | 2,3 | 1,4 | 1,3 | 1,6 | 1,0 | 1,5 |
| Gallo_0411 | 82,39 | 8 | 9 | 7 | 9 | 7 | 10 | 8 | 10 | 4 | 4 | 9 | 12 | 12 | 1,40E+07 | 1,10E+07 | 1,00E+07 | 1,20E+07 | 5,80E+06 | 1,70E+07 | 5,80E+06 | 2,4 | 1,9 | 1,7 | 2,1 | 1,0 | 2,9 |
| Gallo_1578 | 56,51 | 11 | 572 | 12 | 559 | 11 | 558 | 12 | 549 | 12 | 424 | 11 | 582 | 582 | 4,10E+10 | 3,10E+10 | 3,10E+10 | 2,90E+10 | 2,00E+10 | 3,30E+10 | 2,00E+10 | 2,1 | 1,6 | 1,6 | 1,5 | 1,0 | 1,7 |
| Gallo_0162 | 52,34 | 12 | 18 | 8 | 11 | 9 | 21 | 8 | 15 | 8 | 12 | 9 | 19 | 21 | 5,40E+07 | 4,90E+07 | 5,80E+07 | 4,80E+07 | 3,40E+07 | 5,40E+07 | 3,40E+07 | 1,6 | 1,4 | 1,7 | 1,4 | 1,0 | 1,6 |
| Gallo_1847 | 29,41 | 9 | 14 | 7 | 11 | 7 | 10 | 9 | 12 | 8 | 10 | 8 | 11 | 14 | 6,60E+07 | 9,00E+07 | 2,60E+07 | 6,10E+07 | 2,80E+07 | 4,50E+07 | 2,60E+07 | 2,5 | 3,5 | 1,0 | 2,3 | 1,1 | 1,7 |
| Gallo_0330 | 60,73 | 8 | 11 | 6 | 8 | 7 | 13 | 7 | 12 | 10 | 12 | 7 | 9 | 13 | 3,40E+07 | 4,30E+07 | 2,90E+07 | 2,90E+07 | 2,00E+07 | 2,60E+07 | 2,00E+07 | 1,7 | 2,2 | 1,5 | 1,5 | 1,0 | 1,3 |
| Gallo_2179 | 74 | 8 | 12 | 6 | 8 | 7 | 10 | 8 | 11 | 8 | 10 | 7 | 13 | 13 | 4,80E+07 | 4,00E+07 | 3,70E+07 | 3,90E+07 | 3,90E+07 | 5,10E+07 | 3,70E+07 | 1,3 | 1,1 | 1,0 | 1,1 | 1,1 | 1,4 |
| Gallo_1000 | 31,11 | 9 | 9 | 4 | 6 | 8 | 9 | 7 | 7 | 6 | 8 | 8 | 9 | 9 | 5,10E+06 | 7,90E+06 | 4,70E+06 | 4,40E+06 | 3,40E+06 | 6,20E+06 | 3,40E+06 | 1,5 | 2,3 | 1,4 | 1,3 | 1,0 | 1,8 |
| Gallo_0596 | 38,85 | 8 | 14 | 9 | 16 | 8 | 13 | 8 | 15 | 10 | 15 | 9 | 15 | 16 | 4,30E+07 | 4,80E+07 | 1,30E+08 | 1,00E+08 | 2,90E+07 | 4,70E+07 | 2,90E+07 | 1,5 | 1,7 | 4,5 | 3,4 | 1,0 | 1,6 |
| Gallo_2052 | 48,41 | 7 | 11 | 8 | 12 | 7 | 11 | 8 | 15 | 5 | 11 | 8 | 13 | 15 | 1,00E+07 | 1,50E+07 | 1,30E+07 | 1,60E+07 | 8,80E+06 | 8,70E+06 | 8,70E+06 | 1,1 | 1,7 | 1,5 | 1,8 | 1,0 | 1,0 |
| Gallo_0874 | 37,38 | 9 | 15 | 8 | 12 | 9 | 13 | 8 | 9 | 9 | 12 | 8 | 10 | 15 | 2,20E+07 | 2,10E+07 | 1,40E+07 | 1,50E+07 | 1,60E+07 | 1,30E+07 | 1,30E+07 | 1,7 | 1,6 | 1,1 | 1,2 | 1,2 | 1,0 |
| Gallo_1996 | 36,01 | 10 | 12 | 9 | 11 | 9 | 11 | 9 | 9 | 7 | 7 | 7 | 10 | 12 | 2,10E+07 | 1,90E+07 | 9,70E+06 | 1,10E+07 | 6,60E+06 | 1,20E+07 | 6,60E+06 | 3,2 | 2,9 | 1,5 | 1,7 | 1,0 | 1,8 |
| Gallo_2040 | 183,71 | 6 | 7 | 4 | 5 | 9 | 11 | 7 | 9 | 4 | 4 | 3 | 3 | 11 | 5,20E+06 | 4,80E+06 | 5,50E+06 | 8,60E+06 | 2,20E+06 | 2,60E+06 | 2,20E+06 | 2,4 | 2,2 | 2,5 | 3,9 | 1,0 | 1,2 |
| Gallo_1760 | 41,54 | 10 | 13 | 8 | 9 | 9 | 12 | 9 | 12 | 6 | 8 | 9 | 12 | 13 | 1,70E+07 | 1,30E+07 | 1,20E+07 | 1,10E+07 | 7,70E+06 | 1,40E+07 | 7,70E+06 | 2,2 | 1,7 | 1,6 | 1,4 | 1,0 | 1,8 |
| Gallo_1405 | 80,49 | 4 | 9 | 6 | 10 | 7 | 11 | 7 | 12 | 5 | 7 | 7 | 13 | 13 | 4,00E+07 | 3,50E+07 | 2,50E+07 | 3,00E+07 | 1,20E+07 | 4,50E+07 | 1,20E+07 | 3,3 | 2,9 | 2,1 | 2,5 | 1,0 | 3,8 |
| Gallo_1772 | 34,47 | 7 | 10 | 6 | 8 | 8 | 12 | 6 | 8 | 7 | 9 | 6 | 8 | 12 | 1,90E+07 | 2,50E+07 | 1,80E+07 | 1,80E+07 | 1,50E+07 | 1,70E+07 | 1,50E+07 | 1,3 | 1,7 | 1,2 | 1,2 | 1,0 | 1,1 |
| Gallo_1844 | 32,04 | 8 | 23 | 9 | 24 | 8 | 23 | 8 | 22 | 8 | 16 | 8 | 21 | 24 | 1,10E+08 | 1,10E+08 | 6,30E+07 | 8,90E+07 | 5,60E+07 | 7,10E+07 | 5,60E+07 | 2,0 | 2,0 | 1,1 | 1,6 | 1,0 | 1,3 |
| Gallo_1332 | 43,93 | 5 | 14 | 4 | 10 | 7 | 14 | 8 | 14 | 7 | 11 | 4 | 13 | 14 | 5,10E+07 | 4,10E+07 | 7,90E+07 | 3,70E+07 | 4,10E+07 | 7,70E+07 | 3,70E+07 | 1,4 | 1,1 | 2,1 | 1,0 | 1,1 | 2,1 |
| Gallo_1197 | 99,89 | 6 | 15 | 3 | 9 | 8 | 14 | 9 | 10 | 4 | 5 | 7 | 12 | 15 | 8,60E+06 | 3,90E+06 | 1,00E+07 | 1,20E+07 | 2,60E+06 | 1,50E+07 | 2,60E+06 | 3,3 | 1,5 | 3,8 | 4,6 | 1,0 | 5,8 |
| Gallo_1845 | 30,68 | 8 | 10 | 8 | 10 | 8 | 9 | 8 | 15 | 8 | 9 | 8 | 11 | 15 | 4,50E+07 | 3,50E+07 | 3,20E+07 | 6,30E+07 | 4,30E+07 | 3,70E+07 | 3,20E+07 | 1,4 | 1,1 | 1,0 | 2,0 | 1,3 | 1,2 |
| Gallo_1274 | 32,39 | 7 | 10 | 6 | 6 | 7 | 9 | 9 | 11 | 9 | 11 | 6 | 7 | 11 | 2,30E+07 | 1,00E+07 | 1,10E+07 | 2,00E+07 | 1,10E+07 | 1,40E+07 | 1,00E+07 | 2,3 | 1,0 | 1,1 | 2,0 | 1,1 | 1,4 |
| Gallo_1511 | 43,92 | 8 | 9 | 7 | 9 | 7 | 9 | 5 | 7 | 5 | 6 | 8 | 12 | 12 | 9,30E+06 | 1,20E+07 | 7,70E+06 | 4,80E+06 | 4,80E+06 | 9,50E+06 | 4,80E+06 | 1,9 | 2,5 | 1,6 | 1,0 | 1,0 | 2,0 |
| Gallo_1717 | 29,66 | 8 | 9 | 8 | 9 | 6 | 9 | 5 | 6 | 5 | 7 | 6 | 7 | 9 | 9,10E+06 | 1,30E+07 | 4,60E+06 | 2,80E+06 | 4,00E+06 | 5,60E+06 | 2,80E+06 | 3,3 | 4,6 | 1,6 | 1,0 | 1,4 | 2,0 |
| Gallo_1997 | 76,69 | 7 | 8 | 6 | 7 | 4 | 5 | 2 | 2 | 2 | 2 | 5 | 7 | 8 | 2,00E+06 | 9,70E+06 | 9,70E+05 | 3,60E+05 | 5,20E+05 | 8,90E+05 | 3,60E+05 | 5,6 | 26,9 | 2,7 | 1,0 | 1,4 | 2,5 |
| Gallo_0127 | 59,76 | 3 | 3 | 4 | 4 | 6 | 6 | 6 | 9 | 5 | 5 | 4 | 4 | 9 | 2,00E+06 | 2,30E+06 | 2,10E+06 | 3,20E+06 | 1,40E+06 | 1,80E+06 | 1,40E+06 | 1,4 | 1,6 | 1,5 | 2,3 | 1,0 | 1,3 |
| Gallo_1055 | 157,61 | 4 | 29 | 6 | 27 | 6 | 68 | 4 | 19 | 6 | 29 | 5 | 35 | 68 | 3,60E+08 | 5,00E+08 | 7,70E+08 | 2,80E+08 | 3,10E+08 | 4,80E+08 | 2,80E+08 | 1,3 | 1,8 | 2,8 | 1,0 | 1,1 | 1,7 |
| Gallo_1374 | 93,09 | 7 | 9 | 7 | 11 | 8 | 14 | 8 | 12 | 7 | 9 | 8 | 14 | 14 | 1,10E+07 | 1,70E+07 | 1,00E+07 | 1,20E+07 | 7,10E+06 | 3,00E+07 | 7,10E+06 | 1,5 | 2,4 | 1,4 | 1,7 | 1,0 | 4,2 |
| Gallo_2051 | 50,65 | 7 | 9 | 7 | 11 | 5 | 10 | 7 | 15 | 6 | 10 | 7 | 13 | 15 | 5,20E+07 | 8,30E+07 | 6,10E+07 | 7,70E+07 | 5,40E+07 | 8,50E+07 | 5,20E+07 | 1,0 | 1,6 | 1,2 | 1,5 | 1,0 | 1,6 |
| Gallo_1399 | 44,76 | 7 | 9 | 6 | 7 | 7 | 9 | 6 | 7 | 6 | 7 | 7 | 10 | 10 | 7,30E+06 | 7,30E+06 | 5,10E+06 | 7,00E+06 | 4,20E+06 | 7,30E+06 | 4,20E+06 | 1,7 | 1,7 | 1,2 | 1,7 | 1,0 | 1,7 |
| Gallo_0414 | 30,46 | 7 | 8 | 6 | 8 | 7 | 8 | 7 | 9 | 6 | 9 | 5 | 6 | 9 | 1,70E+07 | 2,60E+07 | 1,40E+07 | 2,30E+07 | 1,70E+07 | 1,30E+07 | 1,30E+07 | 1,3 | 2,0 | 1,1 | 1,8 | 1,3 | 1,0 |
| Gallo_0357 | 42,01 | 5 | 5 | 7 | 10 | 6 | 9 | 4 | 5 | 4 | 12 | 4 | 5 | 12 | 1,10E+07 | 1,60E+07 | 9,10E+06 | 5,10E+06 | 7,80E+06 | 1,30E+07 | 5,10E+06 | 2,2 | 3,1 | 1,8 | 1,0 | 1,5 | 2,5 |
| Gallo_1458 | 47,1 | 8 | 10 | 4 | 4 | 5 | 6 | 5 | 5 | 7 | 9 | 6 | 7 | 10 | 1,30E+07 | 2,90E+06 | 3,60E+06 | 3,90E+06 | 4,60E+06 | 5,00E+06 | 2,90E+06 | 4,5 | 1,0 | 1,2 | 1,3 | 1,6 | 1,7 |
| Gallo_0323 | 44,71 | 5 | 6 | 5 | 5 | 5 | 7 | 5 | 8 | 5 | 5 | 6 | 10 | 10 | 8,40E+06 | 1,00E+07 | 6,70E+06 | 9,40E+06 | 5,70E+06 | 1,20E+07 | 5,70E+06 | 1,5 | 1,8 | 1,2 | 1,6 | 1,0 | 2,1 |
| Gallo_0859 | 26,55 | 7 | 46 | 5 | 23 | 6 | 29 | 6 | 27 | 7 | 30 | 6 | 24 | 46 | 8,40E+08 | 7,70E+08 | 6,30E+08 | 7,10E+08 | 6,10E+08 | 6,60E+08 | 6,10E+08 | 1,4 | 1,3 | 1,0 | 1,2 | 1,0 | 1,1 |
| Gallo_0559 | 15,39 | 6 | 29 | 6 | 42 | 6 | 31 | 0 | 0 | 0 | 0 | 0 | 0 | 42 | 1,40E+08 | 1,70E+08 | 7,10E+07 | NF | NF | NF | 7,10E+07 | 2,0 | 2,4 | 1,0 | 0,0 | 0,0 | 0,0 |
| Gallo_0673 | 39,74 | 5 | 6 | 5 | 7 | 5 | 14 | 7 | 14 | 6 | 12 | 5 | 9 | 14 | 1,10E+07 | 2,00E+07 | 2,10E+07 | 2,60E+07 | 2,00E+07 | 1,60E+07 | 1,10E+07 | 1,0 | 1,8 | 1,9 | 2,4 | 1,8 | 1,5 |
| Gallo_1778 | 35,43 | 6 | 8 | 4 | 7 | 6 | 10 | 7 | 11 | 6 | 10 | 5 | 10 | 11 | 9,70E+06 | 4,90E+06 | 8,10E+06 | 9,10E+06 | 7,10E+06 | 9,80E+06 | 4,90E+06 | 2,0 | 1,0 | 1,7 | 1,9 | 1,4 | 2,0 |
| Gallo_0484 | 54,35 | 5 | 5 | 5 | 6 | 4 | 6 | 4 | 6 | 7 | 8 | 4 | 5 | 8 | 8,10E+06 | 1,10E+07 | 8,90E+06 | 1,50E+07 | 1,10E+07 | 1,30E+07 | 8,10E+06 | 1,0 | 1,4 | 1,1 | 1,9 | 1,4 | 1,6 |
| Gallo_2039 | 50,43 | 6 | 6 | 6 | 6 | 6 | 7 | 6 | 6 | 3 | 4 | 5 | 6 | 7 | 7,30E+06 | 6,60E+06 | 8,60E+06 | 9,30E+06 | 6,20E+06 | 7,40E+06 | 6,20E+06 | 1,2 | 1,1 | 1,4 | 1,5 | 1,0 | 1,2 |
| Gallo_1675 | 81,25 | 5 | 6 | 4 | 4 | 5 | 6 | 6 | 7 | 4 | 4 | 5 | 6 | 7 | 4,90E+06 | 4,60E+06 | 5,90E+06 | 8,30E+06 | 4,20E+06 | 7,00E+06 | 4,20E+06 | 1,2 | 1,1 | 1,4 | 2,0 | 1,0 | 1,7 |
| Gallo_0748 | 168,44 | 4 | 6 | 4 | 4 | 5 | 7 | 5 | 6 | 4 | 4 | 5 | 6 | 7 | 2,60E+06 | 3,30E+06 | 5,40E+06 | 4,70E+06 | 3,30E+06 | 7,70E+06 | 2,60E+06 | 1,0 | 1,3 | 2,1 | 1,8 | 1,3 | 3,0 |
| Gallo_1284 | 34,24 | 5 | 5 | 5 | 5 | 6 | 7 | 5 | 6 | 6 | 6 | 4 | 4 | 7 | 7,10E+06 | 7,50E+06 | 5,90E+06 | 7,40E+06 | 4,60E+06 | 3,90E+06 | 3,90E+06 | 1,8 | 1,9 | 1,5 | 1,9 | 1,2 | 1,0 |
| Gallo_1405 | 80,17 | 4 | 5 | 3 | 5 | 5 | 5 | 4 | 6 | 6 | 8 | 3 | 3 | 8 | 1,50E+06 | 1,20E+06 | 2,00E+06 | 9,80E+05 | 1,30E+06 | 6,10E+05 | 6,10E+05 | 2,5 | 2,0 | 3,3 | 1,6 | 2,1 | 1,0 |
| Gallo_2024 | 51,27 | 4 | 4 | 5 | 7 | 6 | 7 | 5 | 5 | 5 | 5 | 4 | 4 | 7 | 2,50E+06 | 7,90E+06 | 3,70E+06 | 2,50E+06 | 2,30E+06 | 1,40E+06 | 1,40E+06 | 1,8 | 5,6 | 2,6 | 1,8 | 1,6 | 1,0 |
| Gallo_1358 | 80,8 | 3 | 3 | 4 | 4 | 3 | 3 | 3 | 3 | 3 | 3 | 1 | 1 | 4 | 1,70E+06 | 1,70E+06 | 7,70E+05 | 7,90E+05 | 8,70E+05 | 4,70E+05 | 4,70E+05 | 3,6 | 3,6 | 1,6 | 1,7 | 1,9 | 1,0 |
| Gallo_2098 | 49,69 | 3 | 3 | 5 | 6 | 2 | 2 | 0 | 0 | 2 | 3 | 2 | 2 | 6 | 2,70E+05 | 1,40E+06 | 1,20E+05 | NF | 2,30E+05 | 1,90E+05 | 1,20E+05 | 2,3 | 11,7 | 1,0 | 0,0 | 1,9 | 1,6 |
| Gallo_2071 | 14,72 | 6 | 29 | 5 | 23 | 5 | 36 | 4 | 29 | 4 | 21 | 4 | 21 | 36 | 4,60E+08 | 4,00E+08 | 4,30E+08 | 3,10E+08 | 2,70E+08 | 3,70E+08 | 2,70E+08 | 1,7 | 1,5 | 1,6 | 1,1 | 1,0 | 1,4 |
| Gallo_1392 | 29,08 | 5 | 8 | 5 | 7 | 6 | 9 | 6 | 11 | 5 | 7 | 4 | 5 | 11 | 6,50E+06 | 6,50E+06 | 4,80E+06 | 1,30E+07 | 4,80E+06 | 3,80E+06 | 3,80E+06 | 1,7 | 1,7 | 1,3 | 3,4 | 1,3 | 1,0 |
| Gallo_1394 | 28,81 | 4 | 4 | 3 | 3 | 4 | 5 | 4 | 5 | 3 | 3 | 4 | 4 | 5 | 1,90E+06 | 1,50E+06 | 1,20E+06 | 1,40E+06 | 5,60E+05 | 2,30E+06 | 5,60E+05 | 3,4 | 2,7 | 2,1 | 2,5 | 1,0 | 4,1 |
| Gallo_1270 | 31,95 | 3 | 3 | 3 | 3 | 5 | 5 | 5 | 7 | 3 | 3 | 2 | 2 | 7 | 8,50E+05 | 9,30E+05 | 9,60E+05 | 1,70E+06 | 6,70E+05 | 3,60E+05 | 3,60E+05 | 2,4 | 2,6 | 2,7 | 4,7 | 1,9 | 1,0 |
| Gallo_0990 | 54,16 | 5 | 5 | 3 | 4 | 2 | 2 | 1 | 1 | 3 | 3 | 2 | 3 | 5 | 1,80E+06 | 1,20E+06 | 8,80E+05 | 4,60E+05 | 8,80E+05 | 9,90E+05 | 4,60E+05 | 3,9 | 2,6 | 1,9 | 1,0 | 1,9 | 2,2 |
| Gallo_1345 | 45,25 | 2 | 3 | 2 | 2 | 4 | 4 | 2 | 2 | 4 | 5 | 1 | 1 | 5 | 1,50E+06 | 1,60E+06 | 1,90E+06 | 1,10E+06 | 2,20E+06 | 4,80E+05 | 4,80E+05 | 3,1 | 3,3 | 4,0 | 2,3 | 4,6 | 1,0 |
| Gallo_0410 | 12,29 | 4 | 6 | 5 | 9 | 3 | 8 | 4 | 12 | 5 | 8 | 5 | 13 | 13 | 2,70E+07 | 4,90E+07 | 3,00E+07 | 4,30E+07 | 2,90E+07 | 3,50E+07 | 2,70E+07 | 1,0 | 1,8 | 1,1 | 1,6 | 1,1 | 1,3 |
| Gallo_2018 | 37,01 | 4 | 6 | 4 | 4 | 5 | 7 | 4 | 5 | 4 | 6 | 4 | 4 | 7 | 7,10E+07 | 2,60E+07 | 3,30E+07 | 4,50E+07 | 3,80E+07 | 1,10E+07 | 1,10E+07 | 6,5 | 2,4 | 3,0 | 4,1 | 3,5 | 1,0 |
| Gallo_0539 | 39,08 | 4 | 5 | 3 | 5 | 4 | 5 | 5 | 5 | 4 | 5 | 4 | 5 | 5 | 8,60E+06 | 9,10E+06 | 4,10E+06 | 5,30E+06 | 5,90E+06 | 6,20E+06 | 4,10E+06 | 2,1 | 2,2 | 1,0 | 1,3 | 1,4 | 1,5 |
| Gallo_0239 | 28,06 | 2 | 2 | 2 | 4 | 4 | 5 | 3 | 4 | 5 | 6 | 4 | 6 | 6 | 2,50E+05 | 2,00E+06 | 2,30E+06 | 2,60E+06 | 1,70E+06 | 2,60E+06 | 2,50E+05 | 1,0 | 8,0 | 9,2 | 10,4 | 6,8 | 10,4 |
| Gallo_1247 | 12,28 | 5 | 6 | 3 | 5 | 3 | 3 | 3 | 4 | 3 | 5 | 4 | 4 | 6 | 7,90E+06 | 6,60E+06 | 3,50E+06 | 4,60E+06 | 4,80E+06 | 4,70E+06 | 3,50E+06 | 2,3 | 1,9 | 1,0 | 1,3 | 1,4 | 1,3 |
| Gallo_1430 | 54,57 | 2 | 4 | 1 | 2 | 5 | 5 | 2 | 2 | 3 | 4 | 4 | 4 | 5 | 2,60E+06 | 1,10E+06 | 4,50E+06 | 9,60E+05 | 3,70E+06 | 3,40E+06 | 9,60E+05 | 2,7 | 1,1 | 4,7 | 1,0 | 3,9 | 3,5 |
| Gallo_0592 | 19,46 | 5 | 5 | 3 | 3 | 3 | 3 | 3 | 3 | 2 | 2 | 3 | 5 | 5 | 8,40E+06 | 4,00E+06 | 2,70E+06 | 2,50E+06 | 1,70E+06 | 5,00E+06 | 1,70E+06 | 4,9 | 2,4 | 1,6 | 1,5 | 1,0 | 2,9 |
| Gallo_0536 | 30,85 | 4 | 5 | 3 | 4 | 1 | 2 | 3 | 4 | 1 | 1 | 4 | 4 | 5 | 7,10E+06 | 6,60E+06 | 7,60E+06 | 4,10E+06 | 2,50E+06 | 5,50E+06 | 2,50E+06 | 2,8 | 2,6 | 3,0 | 1,6 | 1,0 | 2,2 |
| Gallo_0944 | 53,75 | 3 | 3 | 2 | 3 | 3 | 3 | 3 | 4 | 2 | 3 | 3 | 3 | 4 | 1,90E+06 | 1,30E+06 | 1,70E+06 | 2,10E+06 | 1,90E+06 | 1,60E+06 | 1,30E+06 | 1,5 | 1,0 | 1,3 | 1,6 | 1,5 | 1,2 |

| | | | | | | | | | | | | | | | | | | | | | | | | | | | |
|---|---|---|---|---|---|---|---|---|---|---|---|---|---|---|---|---|---|---|---|---|---|---|---|---|---|---|---|
| Gallo_1068 | 46,82 | 3 | 5 | 5 | 7 | 2 | 2 | 0 | 0 | 0 | 0 | 0 | 0 | 7 | 6,20E+06 | 2,80E+07 | 4,70E+05 | NF | NF | NF | 4,70E+05 | 13,2 | 59,6 | 1,0 | 0,0 | 0,0 | 0,0 |
| Gallo_2169 | 31,07 | 3 | 3 | 3 | 3 | 0 | 0 | 2 | 2 | 2 | 2 | 2 | 2 | 3 | 9,70E+05 | 1,20E+07 | NF | 6,60E+06 | 3,20E+05 | 3,20E+05 | 3,20E+05 | 3,0 | 37,5 | 0,0 | 20,6 | 1,0 | 1,0 |
| Gallo_0644 | 26,09 | 3 | 3 | 1 | 1 | 3 | 3 | 1 | 2 | 0 | 0 | 3 | 3 | 3 | 1,80E+06 | 2,20E+05 | 6,30E+05 | 2,30E+05 | NF | 7,90E+05 | 2,20E+05 | 8,2 | 1,0 | 2,9 | 1,0 | 0,0 | 3,6 |
| Gallo_0746 | 51 | 2 | 2 | 2 | 2 | 2 | 2 | 0 | 0 | 2 | 3 | 2 | 3 | 3 | 6,60E+05 | 1,60E+05 | 3,50E+05 | NF | 9,00E+05 | 3,60E+05 | 1,60E+05 | 4,1 | 1,0 | 2,2 | 0,0 | 5,6 | 2,3 |
| Gallo_0076 | 34,51 | 0 | 0 | 0 | 0 | 1 | 1 | 4 | 4 | 2 | 3 | 0 | 0 | 4 | NF | NF | 7,60E+04 | 9,50E+05 | 4,70E+05 | NF | 7,60E+04 | 0,0 | 0,0 | 1,0 | 12,5 | 6,2 | 0,0 |
| Gallo_2259 | 98,38 | 1 | 1 | 0 | 0 | 2 | 2 | 4 | 4 | 0 | 0 | 1 | 1 | 4 | 4,90E+05 | NF | 5,40E+05 | 1,50E+06 | NF | 4,10E+05 | 4,10E+05 | 1,2 | 0,0 | 1,3 | 3,7 | 0,0 | 1,0 |
| Gallo_0740 | 26,25 | 3 | 22 | 3 | 25 | 3 | 23 | 3 | 22 | 3 | 17 | 3 | 24 | 25 | 5,60E+07 | 7,60E+07 | 3,20E+07 | 4,00E+07 | 4,30E+07 | 5,30E+07 | 3,20E+07 | 1,8 | 2,4 | 1,0 | 1,3 | 1,3 | 1,7 |
| Gallo_2194 | 16,47 | 4 | 12 | 4 | 14 | 4 | 11 | 4 | 7 | 4 | 6 | 4 | 17 | 17 | 5,70E+07 | 4,60E+07 | 5,00E+07 | 3,00E+07 | 2,80E+07 | 4,80E+07 | 2,80E+07 | 2,0 | 1,6 | 1,8 | 1,1 | 1,0 | 1,7 |
| Gallo_2261 | 42,52 | 4 | 13 | 3 | 12 | 4 | 12 | 3 | 9 | 4 | 10 | 3 | 8 | 13 | 3,90E+07 | 2,80E+07 | 2,40E+07 | 2,70E+07 | 2,00E+07 | 2,70E+07 | 2,00E+07 | 2,0 | 1,4 | 1,2 | 1,4 | 1,0 | 1,4 |
| Gallo_0554 | 17,97 | 4 | 10 | 3 | 8 | 4 | 8 | 4 | 11 | 4 | 10 | 4 | 15 | 15 | 7,70E+07 | 5,50E+07 | 2,00E+07 | 5,50E+07 | 6,20E+07 | 8,70E+07 | 2,00E+07 | 3,9 | 2,8 | 1,0 | 2,8 | 3,1 | 4,4 |
| Gallo_2071 | 14,77 | 4 | 17 | 2 | 5 | 3 | 15 | 3 | 7 | 2 | 8 | 2 | 9 | 17 | 7,60E+07 | 4,10E+07 | 1,10E+08 | 5,10E+07 | 4,60E+07 | 8,20E+07 | 4,10E+07 | 1,9 | 1,0 | 2,7 | 1,2 | 1,1 | 2,0 |
| Gallo_1577 | 85,62 | 4 | 8 | 3 | 6 | 4 | 7 | 4 | 7 | 4 | 6 | 4 | 8 | 8 | 8,00E+07 | 5,50E+07 | 4,10E+07 | 4,20E+07 | 3,60E+07 | 6,40E+07 | 3,60E+07 | 2,2 | 1,5 | 1,1 | 1,2 | 1,0 | 1,8 |
| Gallo_0878 | 22,85 | 3 | 8 | 2 | 4 | 3 | 9 | 2 | 4 | 3 | 8 | 3 | 7 | 9 | 6,40E+07 | 4,20E+07 | 8,90E+07 | 4,90E+07 | 7,60E+07 | 1,10E+08 | 4,20E+07 | 1,5 | 1,0 | 2,1 | 1,2 | 1,8 | 2,6 |
| Gallo_1430 | 54,31 | 3 | 6 | 3 | 7 | 4 | 7 | 3 | 6 | 4 | 7 | 4 | 6 | 7 | 4,60E+06 | 7,30E+06 | 8,20E+06 | 7,10E+06 | 5,20E+06 | 6,60E+06 | 4,60E+06 | 1,0 | 1,6 | 1,8 | 1,5 | 1,1 | 1,4 |
| Gallo_1332 | 43,95 | 2 | 4 | 2 | 6 | 3 | 6 | 4 | 7 | 4 | 6 | 4 | 7 | 7 | 2,50E+07 | 3,30E+07 | 2,50E+07 | 2,50E+07 | 1,50E+07 | 2,10E+07 | 1,50E+07 | 1,7 | 2,2 | 1,7 | 1,7 | 1,0 | 1,4 |
| Gallo_0535 | 27,22 | 3 | 7 | 2 | 2 | 3 | 3 | 3 | 4 | 2 | 2 | 4 | 5 | 7 | 1,80E+07 | 1,40E+06 | 4,80E+06 | 7,60E+06 | 4,50E+06 | 7,20E+06 | 1,40E+06 | 12,9 | 1,0 | 3,4 | 5,4 | 3,2 | 5,1 |
| Gallo_0745 | 63,15 | 3 | 4 | 2 | 4 | 2 | 3 | 2 | 3 | 3 | 4 | 3 | 4 | 4 | 5,40E+05 | 7,90E+06 | 4,20E+05 | 2,90E+05 | 4,70E+05 | 3,70E+05 | 2,90E+05 | 1,9 | 27,2 | 1,4 | 1,0 | 1,6 | 1,3 |
| Gallo_2039 | 51,48 | 3 | 3 | 3 | 5 | 3 | 3 | 3 | 3 | 3 | 3 | 4 | 4 | 5 | 3,50E+06 | 4,50E+06 | 2,40E+06 | 2,80E+06 | 2,10E+06 | 2,90E+06 | 2,10E+06 | 1,7 | 2,1 | 1,1 | 1,3 | 1,0 | 1,4 |
| Gallo_0122 | 72,63 | 3 | 6 | 1 | 2 | 3 | 4 | 4 | 4 | 0 | 0 | 3 | 5 | 6 | 2,70E+06 | 2,20E+06 | 1,20E+06 | 1,80E+06 | NF | 1,30E+06 | 1,20E+06 | 2,3 | 1,8 | 1,0 | 1,5 | 0,0 | 1,1 |
| Gallo_1330 | 44,14 | 4 | 4 | 0 | 0 | 3 | 3 | 3 | 3 | 2 | 2 | 3 | 5 | 5 | 4,40E+06 | NF | 2,00E+06 | 3,30E+06 | 2,10E+06 | 5,00E+06 | 2,00E+06 | 2,2 | 0,0 | 1,0 | 1,7 | 1,1 | 2,5 |
| Gallo_0017 | 27,15 | 2 | 2 | 2 | 2 | 2 | 3 | 2 | 3 | 3 | 5 | 2 | 2 | 5 | 3,10E+06 | 5,20E+06 | 7,40E+06 | 5,20E+06 | 5,90E+06 | 3,70E+06 | 3,10E+06 | 1,0 | 1,7 | 2,4 | 1,7 | 1,9 | 1,2 |
| Gallo_0243 | 64,88 | 4 | 4 | 2 | 2 | 3 | 3 | 3 | 3 | 0 | 0 | 2 | 2 | 4 | 1,90E+06 | 7,80E+05 | 1,70E+06 | 1,60E+06 | NF | 1,40E+06 | 7,80E+05 | 2,4 | 1,0 | 2,2 | 2,1 | 0,0 | 1,8 |
| Gallo_2177 | 30,13 | 0 | 0 | 1 | 1 | 0 | 0 | 3 | 5 | 3 | 3 | 4 | 4 | 5 | NF | 9,30E+04 | NF | 1,70E+06 | 9,60E+05 | 1,50E+06 | 9,30E+04 | 0,0 | 1,0 | 0,0 | 18,3 | 10,3 | 16,1 |
| Gallo_1123 | 62,79 | 3 | 5 | 2 | 2 | 2 | 2 | 0 | 0 | 1 | 1 | 3 | 3 | 5 | 8,20E+05 | 3,10E+05 | 2,30E+05 | NF | 7,20E+04 | 2,90E+05 | 7,20E+04 | 11,4 | 4,3 | 3,2 | 0,0 | 1,0 | 4,0 |
| Gallo_0255 | 31,34 | 0 | 0 | 0 | 0 | 1 | 1 | 4 | 4 | 3 | 3 | 3 | 3 | 4 | NF | NF | 7,00E+05 | 3,20E+06 | 1,60E+06 | 1,50E+06 | 7,00E+05 | 0,0 | 0,0 | 1,0 | 4,6 | 2,3 | 2,1 |
| Gallo_0662 | 23,36 | 3 | 3 | 1 | 1 | 2 | 2 | 1 | 1 | 0 | 0 | 3 | 3 | 3 | 1,70E+06 | 1,40E+06 | 1,50E+06 | 9,20E+05 | NF | 1,60E+06 | 9,20E+05 | 1,8 | 1,5 | 1,6 | 1,0 | 0,0 | 1,7 |
| Gallo_0981 | 43,58 | 3 | 3 | 1 | 1 | 1 | 1 | 3 | 3 | 2 | 2 | 0 | 0 | 3 | 1,70E+06 | 4,70E+05 | 3,50E+05 | 1,30E+06 | 1,10E+06 | NF | 3,50E+05 | 4,9 | 1,3 | 1,0 | 3,7 | 3,1 | 0,0 |
| Gallo_1953 | 87,29 | 3 | 3 | 0 | 0 | 1 | 1 | 0 | 0 | 2 | 2 | 2 | 2 | 3 | 1,80E+05 | NF | 8,00E+04 | NF | 8,30E+06 | 2,10E+05 | 8,00E+04 | 2,3 | 0,0 | 1,0 | 0,0 | 103,8 | 2,6 |
| Gallo_1330 | 44,24 | 3 | 3 | 3 | 3 | 2 | 2 | 0 | 0 | 0 | 0 | 0 | 0 | 3 | 9,40E+06 | 1,10E+07 | 5,90E+06 | NF | NF | NF | 5,90E+06 | 1,6 | 1,9 | 1,0 | 0,0 | 0,0 | 0,0 |
| Gallo_0251 | 47,14 | 3 | 3 | 0 | 0 | 0 | 0 | 1 | 1 | 2 | 3 | 0 | 0 | 3 | 4,30E+05 | NF | NF | 5,10E+04 | 1,10E+05 | NF | 5,10E+04 | 8,4 | 0,0 | 0,0 | 1,0 | 2,2 | 0,0 |
| Gallo_2168 | 37,63 | 3 | 3 | 0 | 0 | 2 | 3 | 0 | 0 | 0 | 0 | 0 | 0 | 3 | 1,10E+06 | NF | 5,60E+05 | NF | NF | NF | 5,60E+05 | 2,0 | 0,0 | 1,0 | 0,0 | 0,0 | 0,0 |
| Gallo_1574 | 62,54 | 3 | 3 | 0 | 0 | 2 | 2 | 0 | 0 | 0 | 0 | 0 | 0 | 3 | 6,10E+06 | NF | 8,80E+06 | NF | NF | NF | 6,10E+06 | 1,0 | 0,0 | 1,4 | 0,0 | 0,0 | 0,0 |
| Gallo_0085 | 135,18 | 2 | 2 | 1 | 1 | 0 | 0 | 0 | 0 | 0 | 0 | 2 | 2 | 2 | 1,70E+05 | 7,50E+04 | NF | NF | NF | 2,80E+05 | 7,50E+04 | 2,3 | 1,0 | 0,0 | 0,0 | 0,0 | 3,7 |
| Gallo_2244 | 20,49 | 3 | 143 | 3 | 106 | 3 | 146 | 3 | 136 | 3 | 101 | 3 | 126 | 146 | 4,00E+09 | 2,20E+09 | 3,10E+09 | 2,80E+09 | 2,40E+09 | 2,70E+09 | 2,20E+09 | 1,8 | 1,0 | 1,4 | 1,3 | 1,1 | 1,2 |
| Gallo_0560 | 10,33 | 3 | 8 | 3 | 8 | 3 | 9 | 0 | 0 | 0 | 0 | 0 | 0 | 9 | 8,70E+07 | 1,10E+08 | 7,20E+07 | NF | NF | NF | 7,20E+07 | 1,2 | 1,5 | 1,0 | 0,0 | 0,0 | 0,0 |
| Gallo_1907 | 16,78 | 3 | 6 | 1 | 2 | 3 | 4 | 1 | 2 | 2 | 3 | 2 | 4 | 6 | 2,20E+06 | 5,60E+05 | 1,30E+06 | 6,00E+05 | 1,10E+06 | 1,60E+06 | 5,60E+05 | 3,9 | 1,0 | 2,3 | 1,1 | 2,0 | 2,9 |
| Gallo_1632 | 81,44 | 2 | 3 | 2 | 3 | 3 | 4 | 2 | 2 | 2 | 2 | 3 | 6 | 6 | 2,50E+06 | 2,10E+06 | 2,10E+06 | 1,60E+06 | 1,50E+06 | 3,70E+06 | 1,50E+06 | 1,7 | 1,4 | 1,4 | 1,1 | 1,0 | 2,5 |
| Gallo_0636 | 9,7 | 2 | 3 | 2 | 2 | 2 | 3 | 3 | 4 | 2 | 3 | 2 | 3 | 4 | 1,10E+06 | 9,50E+05 | 8,30E+05 | 9,30E+05 | 1,10E+06 | 9,40E+05 | 8,30E+05 | 1,3 | 1,1 | 1,0 | 1,1 | 1,3 | 1,1 |
| Gallo_1307 | 24,52 | 2 | 3 | 1 | 1 | 3 | 4 | 2 | 2 | 3 | 3 | 2 | 2 | 4 | 5,20E+06 | 2,10E+06 | 4,10E+06 | 2,20E+06 | 1,90E+06 | 2,30E+06 | 1,90E+06 | 2,7 | 1,1 | 2,2 | 1,2 | 1,0 | 1,2 |
| Gallo_1327 | 15,05 | 2 | 2 | 3 | 3 | 2 | 2 | 2 | 2 | 2 | 3 | 2 | 2 | 3 | 1,00E+06 | 1,60E+06 | 1,10E+06 | 9,70E+05 | 1,10E+06 | 1,20E+06 | 9,70E+05 | 1,0 | 1,6 | 1,1 | 1,0 | 1,1 | 1,2 |
| Gallo_1068 | 15,6 | 3 | 5 | 3 | 5 | 3 | 4 | 0 | 0 | 0 | 0 | 0 | 0 | 5 | 3,40E+06 | 6,10E+06 | 2,60E+06 | NF | NF | NF | 2,60E+06 | 1,3 | 2,3 | 1,0 | 0,0 | 0,0 | 0,0 |
| Gallo_0850 | 34,76 | 1 | 2 | 2 | 2 | 2 | 2 | 1 | 4 | 0 | 0 | 2 | 2 | 4 | 1,20E+06 | 9,90E+05 | 3,60E+06 | 1,40E+06 | NF | 4,10E+06 | 9,90E+05 | 1,2 | 1,0 | 3,6 | 1,4 | 0,0 | 4,1 |
| Gallo_1921 | 34,99 | 2 | 2 | 0 | 0 | 2 | 2 | 2 | 2 | 2 | 2 | 3 | 3 | 3 | 1,10E+06 | NF | 6,60E+05 | 1,30E+06 | 8,40E+05 | 1,30E+06 | 6,60E+05 | 1,7 | 0,0 | 1,0 | 2,0 | 1,3 | 2,0 |
| Gallo_0065 | 19,42 | 1 | 1 | 1 | 1 | 3 | 5 | 2 | 3 | 0 | 0 | 0 | 0 | 5 | 2,80E+05 | 3,40E+05 | 7,00E+05 | 4,80E+05 | NF | NF | 2,80E+05 | 1,0 | 1,2 | 2,5 | 1,7 | 0,0 | 0,0 |
| Gallo_0877 | 37,27 | 3 | 3 | 1 | 1 | 3 | 3 | 1 | 1 | 1 | 1 | 1 | 1 | 3 | 5,20E+05 | 2,10E+05 | 6,00E+05 | 1,40E+05 | 1,30E+05 | 1,40E+05 | 1,30E+05 | 4,0 | 1,6 | 4,6 | 1,1 | 1,0 | 1,1 |
| Gallo_1172 | 96,14 | 2 | 2 | 3 | 3 | 1 | 1 | 1 | 1 | 1 | 1 | 1 | 2 | 3 | 1,20E+06 | 1,80E+06 | 4,60E+05 | 4,80E+05 | 4,50E+05 | 1,00E+06 | 4,50E+05 | 2,7 | 4,0 | 1,0 | 1,1 | 1,0 | 2,2 |
| Gallo_1040 | 22,19 | 3 | 3 | 3 | 3 | 0 | 0 | 0 | 0 | 0 | 0 | 3 | 3 | 3 | 3,90E+06 | 4,40E+06 | NF | NF | NF | 4,10E+06 | 3,90E+06 | 1,0 | 1,1 | 0,0 | 0,0 | 0,0 | 1,1 |
| Gallo_1779 | 33,43 | 2 | 2 | 0 | 0 | 2 | 2 | 1 | 1 | 0 | 0 | 2 | 3 | 3 | 7,80E+05 | NF | 6,50E+05 | 2,90E+05 | NF | 8,70E+05 | 2,90E+05 | 2,7 | 0,0 | 2,2 | 1,0 | 0,0 | 3,0 |
| Gallo_0062 | 19,87 | 1 | 1 | 3 | 4 | 1 | 1 | 1 | 1 | 0 | 0 | 0 | 0 | 4 | 4,30E+05 | 2,00E+06 | 4,20E+05 | 5,00E+05 | NF | NF | 4,20E+05 | 1,0 | 4,8 | 1,0 | 1,2 | 0,0 | 0,0 |
| Gallo_1025 | 36,14 | 2 | 2 | 0 | 0 | 3 | 3 | 0 | 0 | 0 | 0 | 2 | 2 | 3 | 3,30E+05 | NF | 1,20E+06 | NF | NF | 9,50E+06 | 3,30E+05 | 1,0 | 0,0 | 3,6 | 0,0 | 0,0 | 28,8 |
| Gallo_0067 | 17,09 | 1 | 1 | 2 | 3 | 3 | 3 | 0 | 0 | 0 | 0 | 0 | 0 | 3 | 6,10E+05 | 1,50E+06 | 9,60E+05 | NF | NF | NF | 6,10E+05 | 1,0 | 2,5 | 1,6 | 0,0 | 0,0 | 0,0 |
| Gallo_2228 | 31,84 | 1 | 1 | 0 | 0 | 1 | 1 | 1 | 1 | 1 | 1 | 2 | 2 | 2 | 3,80E+05 | NF | 6,20E+05 | 5,00E+05 | 5,20E+05 | 1,90E+06 | 3,80E+05 | 1,0 | 0,0 | 1,6 | 1,3 | 1,4 | 5,0 |
| Gallo_1640 | 20,55 | 1 | 1 | 0 | 0 | 2 | 2 | 0 | 0 | 0 | 0 | 2 | 3 | 3 | 2,60E+05 | NF | 3,30E+05 | NF | NF | 4,30E+05 | 2,60E+05 | 1,0 | 0,0 | 1,3 | 0,0 | 0,0 | 1,7 |
| Gallo_0418 | 32,89 | 0 | 0 | 2 | 3 | 0 | 0 | 3 | 3 | 0 | 0 | 0 | 0 | 3 | NF | 6,50E+07 | NF | 1,90E+07 | NF | NF | 1,90E+07 | 0,0 | 3,4 | 0,0 | 1,0 | 0,0 | 0,0 |
| Gallo_0830 | 22,3 | 3 | 3 | 0 | 0 | 0 | 0 | 0 | 0 | 2 | 2 | 0 | 0 | 3 | 6,20E+05 | NF | NF | NF | 3,90E+05 | NF | 3,90E+05 | 1,6 | 0,0 | 0,0 | 0,0 | 1,0 | 0,0 |
| Gallo_1395 | 29,9 | 1 | 1 | 2 | 2 | 0 | 0 | 0 | 0 | 1 | 1 | 0 | 0 | 2 | 3,20E+05 | 5,70E+05 | NF | NF | 7,10E+04 | NF | 7,10E+04 | 4,5 | 8,0 | 0,0 | 0,0 | 1,0 | 0,0 |
| Gallo_0071 | 23,51 | 2 | 2 | 0 | 0 | 2 | 2 | 0 | 0 | 0 | 0 | 0 | 0 | 2 | 1,40E+05 | NF | 1,30E+05 | NF | NF | NF | 1,30E+05 | 1,1 | 0,0 | 1,0 | 0,0 | 0,0 | 0,0 |
| Gallo_1816 | 63,47 | 1 | 1 | 0 | 0 | 2 | 2 | 0 | 0 | 0 | 0 | 0 | 0 | 2 | 1,10E+05 | NF | 1,50E+06 | NF | NF | NF | 1,10E+05 | 1,0 | 0,0 | 13,6 | 0,0 | 0,0 | 0,0 |
| Gallo_1578 | 63,82 | 1 | 24 | 1 | 33 | 2 | 20 | 2 | 26 | 2 | 27 | 2 | 36 | 36 | 1,30E+07 | 3,90E+07 | 2,10E+07 | 6,00E+07 | 7,30E+07 | 5,70E+07 | 1,30E+07 | 1,0 | 3,0 | 1,6 | 4,6 | 5,6 | 4,4 |
| Gallo_1055 | 167,68 | 2 | 24 | 2 | 11 | 2 | 62 | 2 | 12 | 2 | 9 | 2 | 17 | 62 | 4,70E+06 | 1,50E+06 | 8,80E+06 | 1,10E+06 | 2,40E+06 | 1,50E+06 | 1,10E+06 | 4,3 | 1,4 | 8,0 | 1,0 | 2,2 | 1,4 |
| Gallo_1989 | 22,67 | 2 | 9 | 2 | 7 | 2 | 16 | 2 | 14 | 2 | 13 | 2 | 17 | 17 | 2,60E+08 | 2,30E+08 | 6,60E+08 | 5,50E+08 | 5,30E+08 | 6,40E+08 | 2,30E+08 | 1,1 | 1,0 | 2,9 | 2,4 | 2,3 | 2,8 |
| Gallo_1055 | 9,44 | 2 | 7 | 1 | 7 | 2 | 11 | 2 | 5 | 1 | 3 | 2 | 14 | 14 | 7,50E+07 | 5,30E+07 | 7,30E+07 | 2,20E+07 | 2,80E+07 | 7,60E+07 | 2,20E+07 | 3,4 | 2,4 | 3,3 | 1,0 | 1,3 | 3,5 |
| Gallo_0012 | 47,43 | 1 | 2 | 2 | 4 | 1 | 4 | 1 | 3 | 1 | 3 | 1 | 5 | 5 | 4,90E+07 | 9,00E+07 | 3,20E+07 | 6,70E+07 | 6,10E+07 | 4,60E+07 | 3,20E+07 | 1,5 | 2,8 | 1,0 | 2,1 | 1,9 | 1,4 |
| Gallo_1326 | 13,74 | 2 | 3 | 1 | 2 | 2 | 3 | 2 | 3 | 2 | 4 | 2 | 3 | 4 | 1,60E+06 | 2,60E+06 | 1,70E+06 | 1,70E+06 | 2,10E+06 | 1,70E+06 | 1,60E+06 | 1,0 | 1,6 | 1,1 | 1,1 | 1,3 | 1,1 |
| Gallo_1845 | 30,8 | 2 | 2 | 2 | 3 | 2 | 3 | 2 | 3 | 2 | 3 | 2 | 3 | 3 | 5,40E+06 | 7,30E+06 | 4,60E+06 | 7,00E+06 | 5,20E+06 | 7,10E+06 | 4,60E+06 | 1,2 | 1,6 | 1,0 | 1,5 | 1,1 | 1,5 |
| Gallo_2228 | 31,86 | 1 | 2 | 2 | 4 | 2 | 2 | 2 | 3 | 1 | 1 | 1 | 2 | 4 | 7,80E+05 | 2,70E+06 | 5,50E+05 | 1,40E+06 | 1,90E+05 | 8,00E+05 | 1,90E+05 | 4,1 | 14,2 | 2,9 | 7,4 | 1,0 | 4,2 |
| Gallo_0850 | 34,71 | 2 | 3 | 2 | 2 | 1 | 1 | 2 | 2 | 2 | 3 | 2 | 2 | 3 | 2,70E+06 | 2,80E+06 | 2,40E+06 | 2,50E+06 | 2,10E+06 | 3,00E+06 | 2,10E+06 | 1,3 | 1,3 | 1,1 | 1,2 | 1,0 | 1,4 |
| Gallo_1717 | 29,82 | 1 | 2 | 1 | 1 | 2 | 3 | 2 | 3 | 1 | 2 | 1 | 2 | 3 | 1,80E+07 | 1,10E+07 | 1,10E+07 | 1,50E+07 | 1,30E+07 | 1,60E+07 | 1,10E+07 | 1,6 | 1,0 | 1,0 | 1,4 | 1,2 | 1,5 |
| Gallo_2230 | 81,08 | 2 | 2 | 2 | 2 | 2 | 2 | 2 | 2 | 2 | 2 | 2 | 2 | 2 | 2,00E+06 | 2,20E+06 | 1,10E+06 | 2,20E+06 | 8,30E+05 | 1,60E+06 | 8,30E+05 | 2,4 | 2,7 | 1,3 | 2,7 | 1,0 | 1,9 |
| Gallo_0562 | 55,23 | 2 | 4 | 2 | 6 | 1 | 2 | 0 | 0 | 0 | 0 | 0 | 0 | 6 | 9,40E+06 | 1,70E+06 | 2,50E+05 | NF | NF | NF | 2,50E+05 | 37,6 | 6,8 | 1,0 | 0,0 | 0,0 | 0,0 |
| Gallo_0132 | 6,91 | 1 | 2 | 2 | 3 | 2 | 2 | 1 | 1 | 2 | 2 | 2 | 2 | 3 | 1,40E+06 | 6,00E+06 | 2,00E+06 | 1,70E+06 | 1,40E+07 | 4,30E+06 | 1,40E+06 | 1,0 | 4,3 | 1,4 | 1,2 | 10,0 | 3,1 |
| Gallo_0041 | 66,95 | 2 | 2 | 2 | 2 | 2 | 2 | 2 | 2 | 2 | 2 | 2 | 2 | 2 | 1,30E+06 | 1,70E+06 | 1,40E+06 | 1,40E+06 | 1,00E+06 | 1,40E+06 | 1,00E+06 | 1,3 | 1,7 | 1,4 | 1,4 | 1,0 | 1,4 |
| Gallo_1710 | 18,5 | 2 | 3 | 1 | 1 | 1 | 1 | 2 | 2 | 2 | 3 | 2 | 2 | 3 | 4,70E+06 | 1,50E+06 | 1,10E+06 | 3,70E+06 | 4,70E+06 | 3,40E+06 | 1,10E+06 | 4,3 | 1,4 | 1,0 | 3,4 | 4,3 | 3,1 |
| Gallo_1367 | 20,21 | 1 | 2 | 2 | 3 | 0 | 0 | 1 | 1 | 2 | 2 | 1 | 1 | 3 | 5,80E+06 | 3,30E+06 | NF | 1,60E+06 | 2,30E+06 | 1,80E+06 | 1,60E+06 | 3,6 | 2,1 | 0,0 | 1,0 | 1,4 | 1,1 |
| Gallo_0556 | 43,61 | 0 | 0 | 0 | 0 | 2 | 2 | 1 | 1 | 2 | 3 | 1 | 3 | 3 | NF | NF | 1,00E+06 | 1,80E+06 | 3,70E+06 | 3,00E+06 | 1,00E+06 | 0,0 | 0,0 | 1,0 | 1,8 | 3,7 | 3,0 |
| Gallo_1540 | 4,98 | 2 | 3 | 2 | 2 | 2 | 2 | 0 | 0 | 2 | 2 | 0 | 0 | 3 | 3,20E+05 | 4,30E+05 | 2,40E+05 | NF | 3,30E+05 | NF | 2,40E+05 | 1,3 | 1,8 | 1,0 | 0,0 | 1,4 | 0,0 |
| Gallo_0798 | 30,78 | 1 | 1 | 0 | 0 | 2 | 2 | 2 | 2 | 1 | 1 | 2 | 3 | 3 | 8,20E+05 | NF | 8,50E+05 | 9,60E+05 | 4,00E+05 | 1,50E+06 | 4,00E+05 | 2,1 | 0,0 | 2,1 | 2,4 | 1,0 | 3,8 |
| Gallo_0314 | 70,92 | 1 | 2 | 1 | 1 | 1 | 1 | 1 | 1 | 1 | 1 | 2 | 3 | 3 | 1,30E+07 | 5,90E+06 | 4,40E+06 | 3,80E+06 | 4,70E+06 | 1,00E+07 | 3,80E+06 | 3,4 | 1,6 | 1,2 | 1,0 | 1,2 | 2,6 |
| Gallo_1537 | 10,69 | 2 | 3 | 0 | 0 | 2 | 2 | 2 | 2 | 1 | 1 | 1 | 1 | 3 | 7,40E+05 | NF | 4,20E+05 | 6,70E+05 | 3,30E+05 | 2,80E+05 | 2,80E+05 | 2,6 | 0,0 | 1,5 | 2,4 | 1,2 | 1,0 |
| Gallo_1129 | 35,27 | 1 | 2 | 1 | 1 | 2 | 2 | 1 | 1 | 1 | 1 | 1 | 1 | 2 | 6,00E+05 | 2,70E+05 | 2,40E+05 | 2,40E+05 | 2,80E+05 | 2,20E+05 | 2,20E+05 | 2,7 | 1,2 | 1,1 | 1,1 | 1,3 | 1,0 |
| Gallo_1518 | 30,29 | 1 | 1 | 1 | 1 | 1 | 2 | 1 | 1 | 2 | 2 | 1 | 1 | 2 | 1,20E+06 | 1,50E+06 | 2,20E+06 | 1,40E+06 | 9,60E+05 | 1,20E+06 | 9,60E+05 | 1,3 | 1,6 | 2,3 | 1,5 | 1,0 | 1,3 |
| Gallo_0116 | 48,28 | 0 | 0 | 1 | 2 | 2 | 2 | 1 | 1 | 2 | 2 | 1 | 1 | 2 | NF | 2,20E+06 | 7,50E+05 | 3,70E+05 | 8,30E+05 | 2,10E+05 | 2,10E+05 | 0,0 | 10,5 | 3,6 | 1,8 | 4,0 | 1,0 |
| Gallo_1912 | 11,13 | 2 | 2 | 1 | 1 | 2 | 2 | 1 | 2 | 0 | 0 | 1 | 1 | 2 | 1,50E+06 | 4,80E+05 | 1,00E+06 | 1,30E+06 | NF | 8,70E+05 | 4,80E+05 | 3,1 | 1,0 | 2,1 | 2,7 | 0,0 | 1,8 |
| Gallo_0933 | 63,55 | 1 | 1 | 1 | 1 | 1 | 1 | 2 | 2 | 1 | 1 | 1 | 1 | 2 | 1,50E+05 | 6,60E+05 | 1,70E+05 | 5,20E+05 | 1,60E+05 | 1,80E+05 | 1,50E+05 | 1,0 | 4,4 | 1,1 | 3,5 | 1,1 | 1,2 |
| Gallo_0989 | 36,05 | 0 | 0 | 1 | 1 | 2 | 2 | 2 | 2 | 2 | 2 | 0 | 0 | 2 | NF | 2,40E+05 | 2,20E+05 | 3,50E+05 | 2,90E+05 | NF | 2,20E+05 | 0,0 | 1,1 | 1,0 | 1,6 | 1,3 | 0,0 |
| Gallo_0890 | 84,54 | 0 | 0 | 0 | 0 | 2 | 3 | 2 | 4 | 0 | 0 | 0 | 0 | 4 | NF | NF | 6,60E+05 | 1,20E+06 | NF | NF | 6,60E+05 | 0,0 | 0,0 | 1,0 | 1,8 | 0,0 | 0,0 |
| Gallo_1918 | 20,41 | 1 | 1 | 1 | 1 | 2 | 2 | 0 | 0 | 1 | 1 | 1 | 1 | 2 | 1,20E+06 | 8,20E+05 | 9,00E+05 | NF | 1,30E+06 | 1,20E+06 | 8,20E+05 | 1,5 | 1,0 | 1,1 | 0,0 | 1,6 | 1,5 |
| Gallo_0535 | 27,21 | 2 | 3 | 0 | 0 | 0 | 0 | 1 | 1 | 1 | 1 | 1 | 1 | 3 | 1,30E+06 | NF | NF | 4,30E+05 | 2,20E+05 | 2,90E+05 | 2,20E+05 | 5,9 | 0,0 | 0,0 | 2,0 | 1,0 | 1,3 |
| Gallo_0299 | 24,5 | 2 | 2 | 0 | 0 | 1 | 1 | 1 | 1 | 1 | 1 | 1 | 1 | 2 | 3,20E+06 | NF | 1,60E+06 | 1,60E+06 | 1,20E+06 | 2,40E+06 | 1,20E+06 | 2,7 | 0,0 | 1,3 | 1,3 | 1,0 | 2,0 |
| Gallo_1293 | 48,04 | 0 | 0 | 1 | 1 | 2 | 2 | 1 | 1 | 1 | 1 | 1 | 1 | 2 | NF | 8,00E+05 | 2,80E+05 | 3,80E+05 | 3,60E+05 | 3,20E+05 | 2,80E+05 | 0,0 | 2,9 | 1,0 | 1,4 | 1,3 | 1,1 |
| Gallo_0454 | 12,87 | 1 | 2 | 0 | 0 | 1 | 1 | 0 | 0 | 2 | 2 | 0 | 0 | 2 | 1,90E+05 | NF | 2,80E+04 | NF | 4,40E+05 | NF | 2,80E+04 | 6,8 | 0,0 | 1,0 | 0,0 | 15,7 | 0,0 |
| Gallo_0625 | 110,89 | 2 | 2 | 1 | 1 | 0 | 0 | 1 | 1 | 0 | 0 | 1 | 1 | 2 | 2,40E+06 | 1,20E+06 | NF | 1,90E+06 | NF | 7,30E+05 | 7,30E+05 | 3,3 | 1,6 | 0,0 | 2,6 | 0,0 | 1,0 |
| Gallo_2013 | 5,29 | 1 | 1 | 0 | 0 | 1 | 1 | 0 | 0 | 2 | 2 | 1 | 1 | 2 | 5,10E+06 | NF | 6,60E+06 | NF | 2,10E+07 | 2,90E+05 | 2,90E+05 | 17,6 | 0,0 | 22,8 | 0,0 | 72,4 | 1,0 |
| Gallo_2038 | 35,36 | 1 | 1 | 0 | 0 | 0 | 0 | 2 | 2 | 0 | 0 | 2 | 2 | 2 | 1,80E+05 | NF | NF | 5,50E+05 | NF | 7,10E+05 | 1,80E+05 | 1,0 | 0,0 | 0,0 | 3,1 | 0,0 | 3,9 |
| Gallo_0053 | 29,89 | 0 | 0 | 0 | 0 | 2 | 2 | 2 | 2 | 0 | 0 | 1 | 1 | 2 | NF | NF | 2,40E+05 | 4,40E+05 | NF | 3,70E+05 | 2,40E+05 | 0,0 | 0,0 | 1,0 | 1,8 | 0,0 | 1,5 |
| Gallo_0791 | 54,42 | 0 | 0 | 0 | 0 | 0 | 0 | 2 | 2 | 2 | 2 | 1 | 1 | 2 | NF | NF | NF | 3,30E+06 | 3,10E+06 | 3,70E+05 | 3,70E+05 | 0,0 | 0,0 | 0,0 | 8,9 | 8,4 | 1,0 |
| Gallo_1643 | 14,78 | 1 | 1 | 0 | 0 | 1 | 1 | 0 | 0 | 0 | 0 | 2 | 2 | 2 | 3,60E+05 | NF | 8,70E+04 | NF | NF | 4,40E+05 | 8,70E+04 | 4,1 | 0,0 | 1,0 | 0,0 | 0,0 | 5,1 |
| Gallo_1998 | 17,77 | 2 | 2 | 0 | 0 | 1 | 1 | 0 | 0 | 0 | 0 | 1 | 1 | 2 | 7,70E+05 | NF | 9,90E+04 | NF | NF | 2,00E+05 | 9,90E+04 | 7,8 | 0,0 | 1,0 | 0,0 | 0,0 | 2,0 |
| Gallo_0055 | 12,39 | 2 | 2 | 2 | 2 | 0 | 0 | 0 | 0 | 0 | 0 | 0 | 0 | 2 | 5,10E+05 | 6,50E+05 | NF | NF | NF | NF | 5,10E+05 | 1,0 | 1,3 | 0,0 | 0,0 | 0,0 | 0,0 |
| Gallo_0130 | 84,71 | 0 | 0 | 0 | 0 | 0 | 0 | 1 | 2 | 1 | 1 | 1 | 1 | 2 | NF | NF | NF | 4,20E+05 | 2,70E+05 | 2,60E+05 | 2,60E+05 | 0,0 | 0,0 | 0,0 | 1,6 | 1,0 | 1,0 |
| Gallo_0211 | 19,64 | 2 | 2 | 0 | 0 | 0 | 0 | 0 | 0 | 1 | 1 | 1 | 1 | 2 | 4,70E+05 | NF | NF | NF | 1,20E+05 | 1,40E+05 | 1,20E+05 | 3,9 | 0,0 | 0,0 | 0,0 | 1,0 | 1,2 |
| Gallo_0233 | 16,23 | 1 | 1 | 2 | 2 | 0 | 0 | 0 | 0 | 0 | 0 | 1 | 1 | 2 | 2,70E+05 | 9,80E+05 | NF | NF | NF | 2,90E+05 | 2,70E+05 | 1,0 | 3,6 | 0,0 | 0,0 | 0,0 | 1,1 |

| | | | | | | | | | | | | | | | | | | | | | | | | | | | |
|---|---|---|---|---|---|---|---|---|---|---|---|---|---|---|---|---|---|---|---|---|---|---|---|---|---|---|---|
| Gallo_0077 | 14,55 | 2 | 2 | 0 | 0 | 0 | 0 | 0 | 0 | 0 | 0 | 2 | 2 | 2 | 6,50E+05 | NF | NF | NF | NF | 6,00E+05 | 6,00E+05 | 1,1 | 0,0 | 0,0 | 0,0 | 0,0 | 1,0 |
| Gallo_0938 | 28,68 | 0 | 0 | 0 | 0 | 0 | 0 | 0 | 0 | 0 | 0 | 2 | 4 | 4 | NF | NF | NF | NF | NF | 1,50E+06 | 1,50E+06 | 0,0 | 0,0 | 0,0 | 0,0 | 0,0 | 1,0 |
| Gallo_1569 | 54,08 | 0 | 0 | 0 | 0 | 0 | 0 | 2 | 3 | 0 | 0 | 0 | 0 | 3 | NF | NF | NF | 5,30E+05 | NF | NF | 5,30E+05 | 0,0 | 0,0 | 0,0 | 1,0 | 0,0 | 0,0 |
| Gallo_1089 | 74,57 | 1 | 1 | 0 | 0 | 2 | 2 | 0 | 0 | 0 | 0 | 0 | 0 | 2 | 4,10E+05 | NF | 1,10E+06 | NF | NF | NF | 4,10E+05 | 1,0 | 0,0 | 2,7 | 0,0 | 0,0 | 0,0 |
| Gallo_1127 | 27,39 | 0 | 0 | 0 | 0 | 2 | 2 | 0 | 0 | 0 | 0 | 1 | 1 | 2 | NF | NF | 8,30E+05 | NF | NF | 8,70E+05 | 8,30E+05 | 0,0 | 0,0 | 1,0 | 0,0 | 0,0 | 1,0 |
| Gallo_0050 | 22,42 | 2 | 2 | 0 | 0 | 1 | 1 | 0 | 0 | 0 | 0 | 0 | 0 | 2 | 7,50E+05 | NF | 1,10E+05 | NF | NF | NF | 1,10E+05 | 6,8 | 0,0 | 1,0 | 0,0 | 0,0 | 0,0 |
| Gallo_2185 | 83,64 | 0 | 0 | 2 | 2 | 0 | 0 | 0 | 0 | 0 | 0 | 1 | 1 | 2 | NF | 1,20E+07 | NF | NF | NF | 1,10E+05 | 1,10E+05 | 0,0 | 109,1 | 0,0 | 0,0 | 0,0 | 1,0 |
| Gallo_0051 | 22,21 | 2 | 3 | 0 | 0 | 0 | 0 | 0 | 0 | 0 | 0 | 0 | 0 | 3 | 3,60E+07 | NF | NF | NF | NF | NF | 3,60E+07 | 1,0 | 0,0 | 0,0 | 0,0 | 0,0 | 0,0 |
| Gallo_1889 | 38,61 | 2 | 3 | 0 | 0 | 0 | 0 | 0 | 0 | 0 | 0 | 0 | 0 | 3 | 7,40E+06 | NF | NF | NF | NF | NF | 7,40E+06 | 1,0 | 0,0 | 0,0 | 0,0 | 0,0 | 0,0 |
| Gallo_1454 | 34,19 | 0 | 0 | 1 | 1 | 0 | 0 | 1 | 1 | 0 | 0 | 0 | 0 | 1 | NF | 1,20E+06 | NF | 1,30E+05 | NF | NF | 1,30E+05 | 0,0 | 9,2 | 0,0 | 1,0 | 0,0 | 0,0 |
| Gallo_1897 | 20,73 | 0 | 0 | 0 | 0 | 1 | 1 | 0 | 0 | 0 | 0 | 1 | 1 | 1 | NF | NF | 1,10E+05 | NF | NF | 3,30E+05 | 1,10E+05 | 0,0 | 0,0 | 1,0 | 0,0 | 0,0 | 3,0 |
| Gallo_0324 | 59,67 | 1 | 3 | 1 | 3 | 1 | 2 | 1 | 3 | 1 | 2 | 1 | 3 | 3 | 2,30E+07 | 2,80E+07 | 1,10E+07 | 2,50E+07 | 1,10E+07 | 1,70E+07 | 1,10E+07 | 2,1 | 2,5 | 1,0 | 2,3 | 1,0 | 1,5 |
| Gallo_1716 | 13,76 | 1 | 1 | 1 | 2 | 1 | 3 | 1 | 2 | 1 | 4 | 1 | 2 | 4 | 1,90E+06 | 3,50E+06 | 1,40E+06 | 2,10E+06 | 1,70E+06 | 1,70E+06 | 1,40E+06 | 1,4 | 2,5 | 1,0 | 1,5 | 1,2 | 1,2 |
| Gallo_0010 | 14,74 | 1 | 2 | 1 | 2 | 1 | 2 | 1 | 3 | 1 | 2 | 1 | 2 | 3 | 4,50E+06 | 7,30E+06 | 3,30E+06 | 6,90E+06 | 6,30E+06 | 5,90E+06 | 3,30E+06 | 1,4 | 2,2 | 1,0 | 2,1 | 1,9 | 1,8 |
| Gallo_1577 | 85,57 | 1 | 5 | 1 | 2 | 1 | 2 | 0 | 0 | 1 | 1 | 1 | 2 | 5 | 7,00E+07 | 2,60E+07 | 1,70E+07 | NF | 9,60E+06 | 1,50E+07 | 9,60E+06 | 7,3 | 2,7 | 1,8 | 0,0 | 1,0 | 1,6 |
| Gallo_0558 | 102,38 | 1 | 1 | 1 | 2 | 1 | 2 | 1 | 2 | 1 | 1 | 1 | 1 | 2 | 8,70E+05 | 3,10E+06 | 1,90E+06 | 2,00E+06 | 6,00E+05 | 8,40E+05 | 6,00E+05 | 1,5 | 5,2 | 3,2 | 3,3 | 1,0 | 1,4 |
| Gallo_1392 | 29,06 | 1 | 1 | 1 | 1 | 1 | 2 | 1 | 2 | 1 | 2 | 1 | 1 | 2 | 1,10E+06 | 1,50E+06 | 8,90E+05 | 1,10E+06 | 1,10E+06 | 1,20E+06 | 8,90E+05 | 1,2 | 1,7 | 1,0 | 1,2 | 1,2 | 1,3 |
| Gallo_0596 | 38,85 | 1 | 1 | 1 | 2 | 1 | 2 | 1 | 1 | 1 | 1 | 1 | 2 | 2 | 6,50E+06 | 1,50E+07 | 1,10E+07 | 6,10E+06 | 4,60E+06 | 1,50E+07 | 4,60E+06 | 1,4 | 3,3 | 2,4 | 1,3 | 1,0 | 3,3 |
| Gallo_1942 | 20,43 | 1 | 2 | 1 | 2 | 1 | 1 | 1 | 1 | 1 | 1 | 1 | 1 | 2 | 3,40E+05 | 6,70E+05 | 2,70E+05 | 4,60E+05 | 2,40E+05 | 3,30E+05 | 2,40E+05 | 1,4 | 2,8 | 1,1 | 1,9 | 1,0 | 1,4 |
| Gallo_1818 | 49,71 | 1 | 1 | 1 | 1 | 1 | 1 | 1 | 2 | 0 | 0 | 1 | 2 | 2 | 4,80E+05 | 1,40E+06 | 3,70E+05 | 1,30E+06 | NF | 7,00E+05 | 3,70E+05 | 1,3 | 3,8 | 1,0 | 3,5 | 0,0 | 1,9 |
| Gallo_2051 | 50,85 | 1 | 1 | 1 | 2 | 1 | 1 | 1 | 3 | 0 | 0 | 0 | 0 | 3 | 1,10E+06 | 4,10E+06 | 8,10E+05 | 3,70E+06 | NF | NF | 8,10E+05 | 1,4 | 5,1 | 1,0 | 4,6 | 0,0 | 0,0 |
| Gallo_1224 | 40,35 | 1 | 1 | 1 | 1 | 1 | 1 | 1 | 1 | 1 | 1 | 1 | 1 | 1 | 2,20E+05 | 3,80E+05 | 1,90E+05 | 3,00E+05 | 3,30E+05 | 3,00E+05 | 1,90E+05 | 1,2 | 2,0 | 1,0 | 1,6 | 1,7 | 1,6 |
| Gallo_0874 | 37,45 | 1 | 1 | 1 | 2 | 1 | 1 | 1 | 1 | 1 | 1 | 0 | 0 | 2 | 1,80E+05 | 2,90E+05 | 1,80E+05 | 3,50E+05 | 3,90E+05 | NF | 1,80E+05 | 1,0 | 1,6 | 1,0 | 1,9 | 2,2 | 0,0 |
| Gallo_2024 | 51,37 | 1 | 1 | 1 | 1 | 0 | 0 | 1 | 2 | 1 | 2 | 0 | 0 | 2 | 1,80E+05 | 1,80E+05 | NF | 3,00E+05 | 2,80E+05 | NF | 1,80E+05 | 1,0 | 1,0 | 0,0 | 1,7 | 1,6 | 0,0 |
| Gallo_1399 | 44,74 | 1 | 1 | 1 | 1 | 1 | 1 | 1 | 1 | 1 | 1 | 1 | 1 | 1 | 1,10E+06 | 1,40E+06 | 8,10E+05 | 7,50E+05 | 8,20E+05 | 1,30E+06 | 7,50E+05 | 1,5 | 1,9 | 1,1 | 1,0 | 1,1 | 1,7 |
| Gallo_1490 | 17,11 | 1 | 2 | 0 | 0 | 1 | 1 | 1 | 1 | 1 | 1 | 1 | 1 | 2 | 3,00E+05 | NF | 2,90E+05 | 4,20E+05 | 3,50E+05 | 3,70E+05 | 2,90E+05 | 1,0 | 0,0 | 1,0 | 1,4 | 1,2 | 1,3 |
| Gallo_0741 | 9,35 | 1 | 1 | 1 | 1 | 1 | 1 | 1 | 1 | 1 | 1 | 1 | 1 | 1 | 3,70E+05 | 5,60E+05 | 2,90E+05 | 4,70E+05 | 3,90E+05 | 3,80E+05 | 2,90E+05 | 1,3 | 1,9 | 1,0 | 1,6 | 1,3 | 1,3 |
| Gallo_2068 | 63,32 | 1 | 1 | 1 | 1 | 1 | 1 | 1 | 1 | 1 | 1 | 1 | 1 | 1 | 7,20E+05 | 1,10E+06 | 7,30E+05 | 8,90E+05 | 7,00E+05 | 7,60E+05 | 7,00E+05 | 1,0 | 1,6 | 1,0 | 1,3 | 1,0 | 1,1 |
| Gallo_1274 | 32,42 | 1 | 1 | 1 | 1 | 1 | 1 | 1 | 1 | 1 | 1 | 1 | 1 | 1 | 2,50E+06 | 1,60E+06 | 1,50E+06 | 2,00E+06 | 1,50E+06 | 1,30E+06 | 1,30E+06 | 1,9 | 1,2 | 1,2 | 1,5 | 1,2 | 1,0 |
| Gallo_1394 | 28,97 | 1 | 1 | 1 | 1 | 1 | 1 | 1 | 1 | 1 | 1 | 1 | 1 | 1 | 1,00E+06 | 1,60E+07 | 9,70E+05 | 8,20E+05 | 4,70E+05 | 9,30E+05 | 4,70E+05 | 2,1 | 34,0 | 2,1 | 1,7 | 1,0 | 2,0 |
| Gallo_0967 | 33,42 | 1 | 1 | 1 | 1 | 1 | 1 | 1 | 1 | 1 | 1 | 1 | 1 | 1 | 2,60E+05 | 3,40E+05 | 1,90E+05 | 3,00E+05 | 5,00E+05 | 3,70E+05 | 1,90E+05 | 1,4 | 1,8 | 1,0 | 1,6 | 2,6 | 1,9 |
| Gallo_1825 | 56,02 | 1 | 1 | 1 | 1 | 1 | 1 | 0 | 0 | 1 | 1 | 1 | 1 | 1 | 5,40E+05 | 5,10E+05 | 2,90E+05 | NF | 3,00E+05 | 3,40E+05 | 2,90E+05 | 1,9 | 1,8 | 1,0 | 0,0 | 1,0 | 1,2 |
| Gallo_1549 | 30,58 | 1 | 1 | 1 | 1 | 0 | 0 | 1 | 2 | 0 | 0 | 1 | 1 | 2 | 3,00E+05 | 5,00E+05 | NF | 5,40E+05 | NF | 3,20E+05 | 3,00E+05 | 1,0 | 1,7 | 0,0 | 1,8 | 0,0 | 1,1 |
| Gallo_1569 | 54,01 | 1 | 1 | 1 | 1 | 0 | 0 | 1 | 2 | 1 | 1 | 0 | 0 | 2 | 4,70E+04 | 5,40E+04 | NF | 1,70E+05 | 1,00E+05 | NF | 4,70E+04 | 1,0 | 1,1 | 0,0 | 3,6 | 2,1 | 0,0 |
| Gallo_1343 | 11,19 | 1 | 1 | 1 | 1 | 1 | 1 | 1 | 1 | 1 | 1 | 0 | 0 | 1 | 5,30E+05 | 5,30E+05 | 4,30E+05 | 4,90E+05 | 2,20E+05 | NF | 2,20E+05 | 2,4 | 2,4 | 2,0 | 2,2 | 1,0 | 0,0 |
| Gallo_1574 | 62,2 | 1 | 1 | 1 | 2 | 1 | 2 | 0 | 0 | 0 | 0 | 0 | 0 | 2 | 2,70E+05 | 1,80E+05 | 1,70E+05 | NF | NF | NF | 1,70E+05 | 1,6 | 1,1 | 1,0 | 0,0 | 0,0 | 0,0 |
| Gallo_1987 | 9,16 | 1 | 2 | 0 | 0 | 0 | 0 | 1 | 2 | 1 | 1 | 0 | 0 | 2 | 2,50E+05 | NF | NF | 2,70E+05 | 1,10E+05 | NF | 1,10E+05 | 2,3 | 0,0 | 0,0 | 2,5 | 1,0 | 0,0 |
| Gallo_0348 | 33,35 | 1 | 1 | 1 | 1 | 0 | 0 | 1 | 1 | 1 | 1 | 1 | 1 | 1 | 1,30E+06 | 1,00E+07 | NF | 1,60E+06 | 3,80E+06 | 1,20E+06 | 1,20E+06 | 1,1 | 8,3 | 0,0 | 1,3 | 3,2 | 1,0 |
| Gallo_2230 | 80,95 | 1 | 2 | 0 | 0 | 1 | 1 | 1 | 1 | 0 | 0 | 1 | 1 | 2 | 4,40E+06 | NF | 9,90E+05 | 1,60E+06 | NF | 1,40E+06 | 9,90E+05 | 4,4 | 0,0 | 1,0 | 1,6 | 0,0 | 1,4 |
| Gallo_0754 | 53,77 | 1 | 1 | 0 | 0 | 1 | 1 | 1 | 1 | 1 | 1 | 1 | 1 | 1 | 8,00E+05 | NF | 5,70E+05 | 7,10E+05 | 5,60E+05 | 6,90E+05 | 5,60E+05 | 1,4 | 0,0 | 1,0 | 1,3 | 1,0 | 1,2 |
| Gallo_0234 | 14,26 | 1 | 1 | 1 | 1 | 1 | 1 | 1 | 1 | 0 | 0 | 1 | 1 | 1 | 8,50E+05 | 8,10E+05 | 4,70E+05 | 3,50E+05 | NF | 5,00E+05 | 3,50E+05 | 2,4 | 2,3 | 1,3 | 1,0 | 0,0 | 1,4 |
| Gallo_0207 | 45,59 | 0 | 0 | 1 | 1 | 1 | 2 | 1 | 1 | 0 | 0 | 0 | 0 | 2 | NF | 2,70E+05 | 2,70E+05 | 1,60E+05 | NF | NF | 1,60E+05 | 0,0 | 1,7 | 1,7 | 1,0 | 0,0 | 0,0 |
| Gallo_2024 | 51,24 | 1 | 1 | 0 | 0 | 1 | 1 | 1 | 1 | 1 | 1 | 0 | 0 | 1 | 3,80E+05 | NF | 3,00E+05 | 4,00E+05 | 3,80E+05 | NF | 3,00E+05 | 1,3 | 0,0 | 1,0 | 1,3 | 1,3 | 0,0 |
| Gallo_0577 | 120,79 | 0 | 0 | 0 | 0 | 1 | 1 | 1 | 1 | 1 | 1 | 1 | 1 | 1 | NF | NF | 2,50E+05 | 3,40E+05 | 3,50E+05 | 3,20E+05 | 2,50E+05 | 0,0 | 0,0 | 1,0 | 1,4 | 1,4 | 1,3 |
| Gallo_1921 | 35,04 | 1 | 1 | 1 | 1 | 1 | 1 | 1 | 1 | 0 | 0 | 0 | 0 | 1 | 1,60E+06 | 1,00E+06 | 7,30E+05 | 9,70E+05 | NF | NF | 7,30E+05 | 2,2 | 1,4 | 1,0 | 1,3 | 0,0 | 0,0 |
| Gallo_1675 | 81,23 | 0 | 0 | 0 | 0 | 1 | 1 | 1 | 1 | 1 | 2 | 0 | 0 | 2 | NF | NF | 8,90E+05 | 6,00E+05 | 1,60E+06 | NF | 6,00E+05 | 0,0 | 0,0 | 1,5 | 1,0 | 2,7 | 0,0 |
| Gallo_0797 | 6,77 | 1 | 1 | 1 | 2 | 0 | 0 | 0 | 0 | 0 | 0 | 1 | 1 | 2 | 3,80E+05 | 9,00E+05 | NF | NF | NF | 3,50E+05 | 3,50E+05 | 1,1 | 2,6 | 0,0 | 0,0 | 0,0 | 1,0 |
| Gallo_0744 | 11,46 | 0 | 0 | 0 | 0 | 1 | 3 | 0 | 0 | 0 | 0 | 1 | 1 | 3 | NF | NF | 5,00E+06 | NF | NF | 2,60E+06 | 2,60E+06 | 0,0 | 0,0 | 1,9 | 0,0 | 0,0 | 1,0 |
| Gallo_0596 | 38,88 | 1 | 2 | 0 | 0 | 0 | 0 | 0 | 0 | 0 | 0 | 1 | 2 | 2 | 6,10E+06 | NF | NF | NF | NF | 5,80E+06 | 5,80E+06 | 1,1 | 0,0 | 0,0 | 0,0 | 0,0 | 1,0 |
| Gallo_1066 | 10,11 | 1 | 1 | 1 | 1 | 1 | 1 | 0 | 0 | 0 | 0 | 0 | 0 | 1 | 8,80E+05 | 1,40E+06 | 8,80E+05 | NF | NF | NF | 8,80E+05 | 1,0 | 1,6 | 1,0 | 0,0 | 0,0 | 0,0 |
| Gallo_2204 | 54,55 | 0 | 0 | 1 | 1 | 1 | 1 | 1 | 1 | 0 | 0 | 0 | 0 | 1 | NF | 2,30E+05 | 1,30E+05 | 2,30E+05 | NF | NF | 1,30E+05 | 0,0 | 1,8 | 1,0 | 1,8 | 0,0 | 0,0 |
| Gallo_0146 | 34,7 | 1 | 1 | 0 | 0 | 0 | 0 | 0 | 0 | 1 | 1 | 1 | 1 | 1 | 1,00E+05 | NF | NF | NF | 8,90E+04 | 1,40E+05 | 8,90E+04 | 1,1 | 0,0 | 0,0 | 0,0 | 1,0 | 1,6 |
| Galo_2258 | 61,03 | 1 | 1 | 0 | 0 | 1 | 1 | 0 | 0 | 1 | 1 | 0 | 0 | 1 | 8,50E+04 | NF | 6,00E+04 | NF | 6,60E+04 | NF | 6,00E+04 | 1,4 | 0,0 | 1,0 | 0,0 | 1,1 | 0,0 |
| Gallo_0060 | 13,01 | 0 | 0 | 1 | 1 | 1 | 1 | 1 | 1 | 0 | 0 | 0 | 0 | 1 | NF | 4,40E+05 | 2,60E+05 | 2,00E+05 | NF | NF | 2,00E+05 | 0,0 | 2,2 | 1,3 | 1,0 | 0,0 | 0,0 |
| Gallo_0617 | 25,26 | 0 | 0 | 0 | 0 | 1 | 3 | 0 | 0 | 0 | 0 | 0 | 0 | 3 | NF | NF | 1,20E+07 | NF | NF | NF | 1,20E+07 | 0,0 | 0,0 | 1,0 | 0,0 | 0,0 | 0,0 |
| Gallo_0367 | 73,62 | 1 | 1 | 1 | 1 | 0 | 0 | 0 | 0 | 0 | 0 | 0 | 0 | 1 | 2,50E+05 | 2,30E+05 | NF | NF | NF | NF | 2,30E+05 | 1,1 | 1,0 | 0,0 | 0,0 | 0,0 | 0,0 |
| Gallo_1057 | 163,31 | 1 | 1 | 0 | 0 | 1 | 1 | 0 | 0 | 0 | 0 | 0 | 0 | 1 | 1,10E+06 | NF | 1,30E+06 | NF | NF | NF | 1,10E+06 | 1,0 | 0,0 | 1,2 | 0,0 | 0,0 | 0,0 |
| Gallo_1359 | 13,64 | 1 | 2 | 0 | 0 | 0 | 0 | 0 | 0 | 0 | 0 | 0 | 0 | 2 | 1,30E+05 | NF | NF | NF | NF | NF | 1,30E+05 | 1,0 | 0,0 | 0,0 | 0,0 | 0,0 | 0,0 |
| Gallo_1559 | 22,68 | 1 | 1 | 1 | 1 | 0 | 0 | 0 | 0 | 0 | 0 | 0 | 0 | 1 | 1,70E+05 | 1,20E+05 | NF | NF | NF | NF | 1,20E+05 | 1,4 | 1,0 | 0,0 | 0,0 | 0,0 | 0,0 |
| Gallo_1573 | 24,11 | 0 | 0 | 0 | 0 | 0 | 0 | 1 | 1 | 1 | 1 | 0 | 0 | 1 | NF | NF | NF | 1,70E+05 | 1,80E+05 | NF | 1,70E+05 | 0,0 | 0,0 | 0,0 | 1,0 | 1,1 | 0,0 |
| Gallo_2134 | 47,65 | 0 | 0 | 1 | 1 | 0 | 0 | 1 | 1 | 0 | 0 | 0 | 0 | 1 | NF | 2,00E+05 | NF | 1,50E+05 | NF | NF | 1,50E+05 | 0,0 | 1,3 | 0,0 | 1,0 | 0,0 | 0,0 |
| Gallo_0817 | 26,34 | 0 | 0 | 0 | 0 | 0 | 0 | 1 | 1 | 1 | 1 | 0 | 0 | 1 | NF | NF | NF | 1,10E+05 | 1,60E+05 | NF | 1,10E+05 | 0,0 | 0,0 | 0,0 | 1,0 | 1,5 | 0,0 |
| Gallo_1140 | 9,05 | 0 | 0 | 1 | 1 | 0 | 0 | 0 | 0 | 0 | 0 | 1 | 1 | 1 | NF | 7,80E+05 | NF | NF | NF | 7,10E+05 | 7,10E+05 | 0,0 | 1,1 | 0,0 | 0,0 | 0,0 | 1,0 |
| Gallo_0307 | 37,25 | 1 | 1 | 0 | 0 | 1 | 1 | 0 | 0 | 0 | 0 | 0 | 0 | 1 | 6,60E+04 | NF | 4,10E+04 | NF | NF | NF | 4,10E+04 | 1,6 | 0,0 | 1,0 | 0,0 | 0,0 | 0,0 |
| Gallo_1067 | 12,22 | 0 | 0 | 0 | 0 | 0 | 0 | 0 | 0 | 1 | 2 | 0 | 0 | 2 | NF | NF | NF | NF | 6,40E+05 | NF | 6,40E+05 | 0,0 | 0,0 | 0,0 | 0,0 | 1,0 | 0,0 |
| Gallo_2141 | 10,43 | 0 | 0 | 0 | 0 | 1 | 2 | 0 | 0 | 0 | 0 | 0 | 0 | 2 | NF | NF | 6,30E+05 | NF | NF | NF | 6,30E+05 | 0,0 | 0,0 | 1,0 | 0,0 | 0,0 | 0,0 |
| Gallo_1985 | 16,66 | 0 | 0 | 0 | 0 | 1 | 2 | 0 | 0 | 0 | 0 | 0 | 0 | 2 | NF | NF | 4,10E+05 | NF | NF | NF | 4,10E+05 | 0,0 | 0,0 | 1,0 | 0,0 | 0,0 | 0,0 |
| Gallo_1571 | 24,69 | 0 | 0 | 0 | 0 | 0 | 0 | 0 | 0 | 1 | 1 | 0 | 0 | 1 | NF | NF | NF | NF | 1,40E+05 | NF | 1,40E+05 | 0,0 | 0,0 | 0,0 | 0,0 | 1,0 | 0,0 |
| Gallo_0725 | 67,72 | 0 | 0 | 1 | 1 | 0 | 0 | 0 | 0 | 0 | 0 | 0 | 0 | 1 | NF | 7,60E+04 | NF | NF | NF | NF | 7,60E+04 | 0,0 | 1,0 | 0,0 | 0,0 | 0,0 | 0,0 |
| Gallo_0603 | 46,33 | 0 | 0 | 0 | 0 | 0 | 0 | 0 | 0 | 0 | 0 | 1 | 1 | 1 | NF | NF | NF | NF | NF | 1,20E+05 | 1,20E+05 | 0,0 | 0,0 | 0,0 | 0,0 | 0,0 | 1,0 |
| Gallo_1489 | 42,95 | 1 | 1 | 0 | 0 | 0 | 0 | 0 | 0 | 0 | 0 | 0 | 0 | 1 | 5,30E+04 | NF | NF | NF | NF | NF | 5,30E+04 | 1,0 | 0,0 | 0,0 | 0,0 | 0,0 | 0,0 |
| Gallo_1899 | 30,81 | 0 | 0 | 0 | 0 | 0 | 0 | 1 | 1 | 0 | 0 | 0 | 0 | 1 | NF | NF | NF | 2,00E+05 | NF | NF | 2,00E+05 | 0,0 | 0,0 | 0,0 | 1,0 | 0,0 | 0,0 |
| Gallo_0340 | 43,83 | 0 | 0 | 0 | 0 | 0 | 0 | 1 | 1 | 0 | 0 | 0 | 0 | 1 | NF | NF | NF | 1,00E+05 | NF | NF | 1,00E+05 | 0,0 | 0,0 | 0,0 | 1,0 | 0,0 | 0,0 |
| Gallo_1248 | 17,55 | 1 | 1 | 0 | 0 | 0 | 0 | 0 | 0 | 0 | 0 | 0 | 0 | 1 | 1,10E+05 | NF | NF | NF | NF | NF | 1,10E+05 | 1,0 | 0,0 | 0,0 | 0,0 | 0,0 | 0,0 |
| Gallo_0609 | 105,29 | 0 | 0 | 1 | 1 | 0 | 0 | 0 | 0 | 0 | 0 | 0 | 0 | 1 | NF | 7,60E+04 | NF | NF | NF | NF | 7,60E+04 | 0,0 | 1,0 | 0,0 | 0,0 | 0,0 | 0,0 |
| Gallo_1510 | 26,57 | 0 | 0 | 0 | 0 | 0 | 0 | 1 | 1 | 0 | 0 | 0 | 0 | 1 | NF | NF | NF | 5,00E+05 | NF | NF | 5,00E+05 | 0,0 | 0,0 | 0,0 | 1,0 | 0,0 | 0,0 |
| Gallo_1917 | 55,21 | 1 | 1 | 0 | 0 | 0 | 0 | 0 | 0 | 0 | 0 | 0 | 0 | 1 | 3,90E+05 | NF | NF | NF | NF | NF | 3,90E+05 | 1,0 | 0,0 | 0,0 | 0,0 | 0,0 | 0,0 |
| Gallo_0704 | 36,88 | 0 | 0 | 0 | 0 | 0 | 0 | 0 | 0 | 0 | 0 | 1 | 1 | 1 | NF | NF | NF | NF | NF | 1,20E+05 | 1,20E+05 | 0,0 | 0,0 | 0,0 | 0,0 | 0,0 | 1,0 |
| Gallo_0494 | 67,6 | 0 | 0 | 0 | 0 | 1 | 1 | 0 | 0 | 0 | 0 | 0 | 0 | 1 | NF | NF | 2,40E+05 | NF | NF | NF | 2,40E+05 | 0,0 | 0,0 | 1,0 | 0,0 | 0,0 | 0,0 |
| Gallo_0748 | 168,46 | 0 | 0 | 0 | 0 | 0 | 0 | 0 | 0 | 0 | 0 | 1 | 1 | 1 | NF | NF | NF | NF | NF | 3,80E+05 | 3,80E+05 | 0,0 | 0,0 | 0,0 | 0,0 | 0,0 | 1,0 |
| Gallo_1821 | 48,73 | 1 | 1 | 0 | 0 | 0 | 0 | 0 | 0 | 0 | 0 | 0 | 0 | 1 | 5,90E+04 | NF | NF | NF | NF | NF | 5,90E+04 | 1,0 | 0,0 | 0,0 | 0,0 | 0,0 | 0,0 |
| Gallo_1873 | 52,6 | 1 | 1 | 0 | 0 | 0 | 0 | 0 | 0 | 0 | 0 | 0 | 0 | 1 | 1,80E+05 | NF | NF | NF | NF | NF | 1,80E+05 | 1,0 | 0,0 | 0,0 | 0,0 | 0,0 | 0,0 |
| Gallo_1404 | 75,94 | 0 | 0 | 0 | 0 | 1 | 1 | 0 | 0 | 0 | 0 | 0 | 0 | 1 | NF | NF | 6,90E+04 | NF | NF | NF | 6,90E+04 | 0,0 | 0,0 | 1,0 | 0,0 | 0,0 | 0,0 |
| Gallo_0056 | 24,18 | 0 | 0 | 0 | 0 | 0 | 0 | 1 | 1 | 0 | 0 | 0 | 0 | 1 | NF | NF | NF | 2,50E+05 | NF | NF | 2,50E+05 | 0,0 | 0,0 | 0,0 | 1,0 | 0,0 | 0,0 |
| Gallo_0719 | 51,32 | 0 | 0 | 0 | 0 | 1 | 1 | 0 | 0 | 0 | 0 | 0 | 0 | 1 | NF | NF | 3,30E+05 | NF | NF | NF | 3,30E+05 | 0,0 | 0,0 | 1,0 | 0,0 | 0,0 | 0,0 |
| Gallo_0608 | 29,34 | 1 | 1 | 0 | 0 | 0 | 0 | 0 | 0 | 0 | 0 | 0 | 0 | 1 | 1,60E+05 | NF | NF | NF | NF | NF | 1,60E+05 | 1,0 | 0,0 | 0,0 | 0,0 | 0,0 | 0,0 |
| Gallo_1992 | 50,13 | 1 | 1 | 0 | 0 | 0 | 0 | 0 | 0 | 0 | 0 | 0 | 0 | 1 | 1,50E+05 | NF | NF | NF | NF | NF | 1,50E+05 | 1,0 | 0,0 | 0,0 | 0,0 | 0,0 | 0,0 |
| Gallo_2055 | 57,4 | 0 | 0 | 0 | 0 | 0 | 0 | 0 | 0 | 0 | 0 | 1 | 1 | 1 | NF | NF | NF | NF | NF | 7,30E+04 | 7,30E+04 | 0,0 | 0,0 | 0,0 | 0,0 | 0,0 | 1,0 |
| Gallo_2232 | 22,97 | 1 | 1 | 0 | 0 | 0 | 0 | 0 | 0 | 0 | 0 | 0 | 0 | 1 | 4,50E+05 | NF | NF | NF | NF | NF | 4,50E+05 | 1,0 | 0,0 | 0,0 | 0,0 | 0,0 | 0,0 |
| Gallo_1152 | 49,3 | 1 | 1 | 0 | 0 | 0 | 0 | 0 | 0 | 0 | 0 | 0 | 0 | 1 | 1,80E+05 | NF | NF | NF | NF | NF | 1,80E+05 | 1,0 | 0,0 | 0,0 | 0,0 | 0,0 | 0,0 |
| Gallo_0064 | 14,75 | 1 | 1 | 0 | 0 | 0 | 0 | 0 | 0 | 0 | 0 | 0 | 0 | 1 | 3,10E+05 | NF | NF | NF | NF | NF | 3,10E+05 | 1,0 | 0,0 | 0,0 | 0,0 | 0,0 | 0,0 |
| Gallo_1607 | 32,48 | 0 | 0 | 0 | 0 | 0 | 0 | 1 | 1 | 0 | 0 | 0 | 0 | 1 | NF | NF | NF | 2,80E+05 | NF | NF | 2,80E+05 | 0,0 | 0,0 | 0,0 | 1,0 | 0,0 | 0,0 |
| Gallo_1498 | 34,29 | 0 | 0 | 0 | 0 | 1 | 1 | 0 | 0 | 0 | 0 | 0 | 0 | 1 | NF | NF | 2,50E+07 | NF | NF | NF | 2,50E+07 | 0,0 | 0,0 | 1,0 | 0,0 | 0,0 | 0,0 |
| Gallo_0334 | 16,92 | 0 | 0 | 0 | 0 | 0 | 0 | 1 | 1 | 0 | 0 | 0 | 0 | 1 | NF | NF | NF | 3,90E+05 | NF | NF | 3,90E+05 | 0,0 | 0,0 | 0,0 | 1,0 | 0,0 | 0,0 |
| Gallo_0706 | 97,96 | 0 | 0 | 0 | 0 | 1 | 1 | 0 | 0 | 0 | 0 | 0 | 0 | 1 | NF | NF | 2,10E+05 | NF | NF | NF | 2,10E+05 | 0,0 | 0,0 | 1,0 | 0,0 | 0,0 | 0,0 |
| Gallo_0127 | 59,79 | 0 | 0 | 0 | 0 | 0 | 0 | 1 | 1 | 0 | 0 | 0 | 0 | 1 | NF | NF | NF | 6,40E+05 | NF | NF | 6,40E+05 | 0,0 | 0,0 | 0,0 | 1,0 | 0,0 | 0,0 |
| Gallo_2189 | 10,41 | 0 | 0 | 0 | 0 | 1 | 1 | 0 | 0 | 0 | 0 | 0 | 0 | 1 | NF | NF | 4,00E+05 | NF | NF | NF | 4,00E+05 | 0,0 | 0,0 | 1,0 | 0,0 | 0,0 | 0,0 |
| Gallo_0306 | 17,86 | 0 | 0 | 0 | 0 | 0 | 0 | 0 | 0 | 1 | 1 | 0 | 0 | 1 | NF | NF | NF | NF | 2,20E+07 | NF | 2,20E+07 | 0,0 | 0,0 | 0,0 | 0,0 | 1,0 | 0,0 |
| Gallo_0084 | 26,16 | 0 | 0 | 0 | 0 | 1 | 1 | 0 | 0 | 0 | 0 | 0 | 0 | 1 | NF | NF | 2,00E+06 | NF | NF | NF | 2,00E+06 | 0,0 | 0,0 | 1,0 | 0,0 | 0,0 | 0,0 |
| Gallo_1347 | 41,54 | 0 | 0 | 0 | 0 | 0 | 0 | 1 | 1 | 0 | 0 | 0 | 0 | 1 | NF | NF | NF | 7,50E+04 | NF | NF | 7,50E+04 | 0,0 | 0,0 | 0,0 | 1,0 | 0,0 | 0,0 |
